# A Plug-and-Play System for Polycyclic Tetramate Macrolactam Expression and Functionalization

**DOI:** 10.1101/2024.10.15.618458

**Authors:** Anna Glöckle, Sebastian Schuler, Manuel Einsiedler, Tobias A. M. Gulder

## Abstract

**Background:** The biosynthesis of the natural product family of the polycyclic tetramate macrolactams (PoTeMs) employs an uncommon iterative polyketide synthase/non-ribosomal peptide synthetase (iPKS/NRPS). This machinery produces a universal PoTeM biosynthetic precursor that contains a tetramic acid moiety connected to two unsaturated polyene side chains. The enormous structural and hence functional diversity of PoTeMs is enabled by pathway-specific tailoring enzymes, particularly cyclization-catalyzing oxidases that process the polyene chains to form distinct ring systems, and further modifying enzymes.

**Results:** Ikarugamycin is the first discovered PoTeM and is formed by the three enzymes IkaABC. Utilizing the iPKS/NRPS IkaA, we established a genetic plug-and-play system by screening eight different strong promoters downstream of *ikaA* to facilitate high-level heterologous expression of PoTeMs in different *Streptomyces* host systems. Furthermore, we applied the system on three different PoTeM modifying genes (*ptmD, ikaD*, and *cftA*), showing the general utility of this approach to study PoTeM post PKS/NRPS processing of diverse tailoring enzymes.

**Conclusion:** By employing our plug-and-play system for PoTeMs, we reconstructed the ikarugamycin biosynthesis and generated five derivatives of ikarugamycin. This platform will generally facilitate the investigation of new PoTeM biosynthetic cyclization and tailoring reactions in the future.

## Background

Polycyclic tetramate macrolactams (PoTeMs) constitute a growing class of complex and structurally diverse natural products mostly isolated from diverse microorganisms.^1^Depending on their individual structures, PoTeMs exhibit a great number of different pharmacological effects such as antibiotic, antifungal, and cytotoxic properties. Some important representatives are ikarugamycin (**1**) from *Streptomyces phaeochromogenes* var. *ikaruganensis*,^2^ alteramide A (**2**) from *Alteromonas* sp.,^3^HSAF (heat-stable antifungal factor, **3**) from *Lysobacter enzymogenes* strain C3,^4,5^and frontalamide A (**4**) from *Streptomyces* sp. SPB78 (Figure 1a).^6^

**Figure 1.**
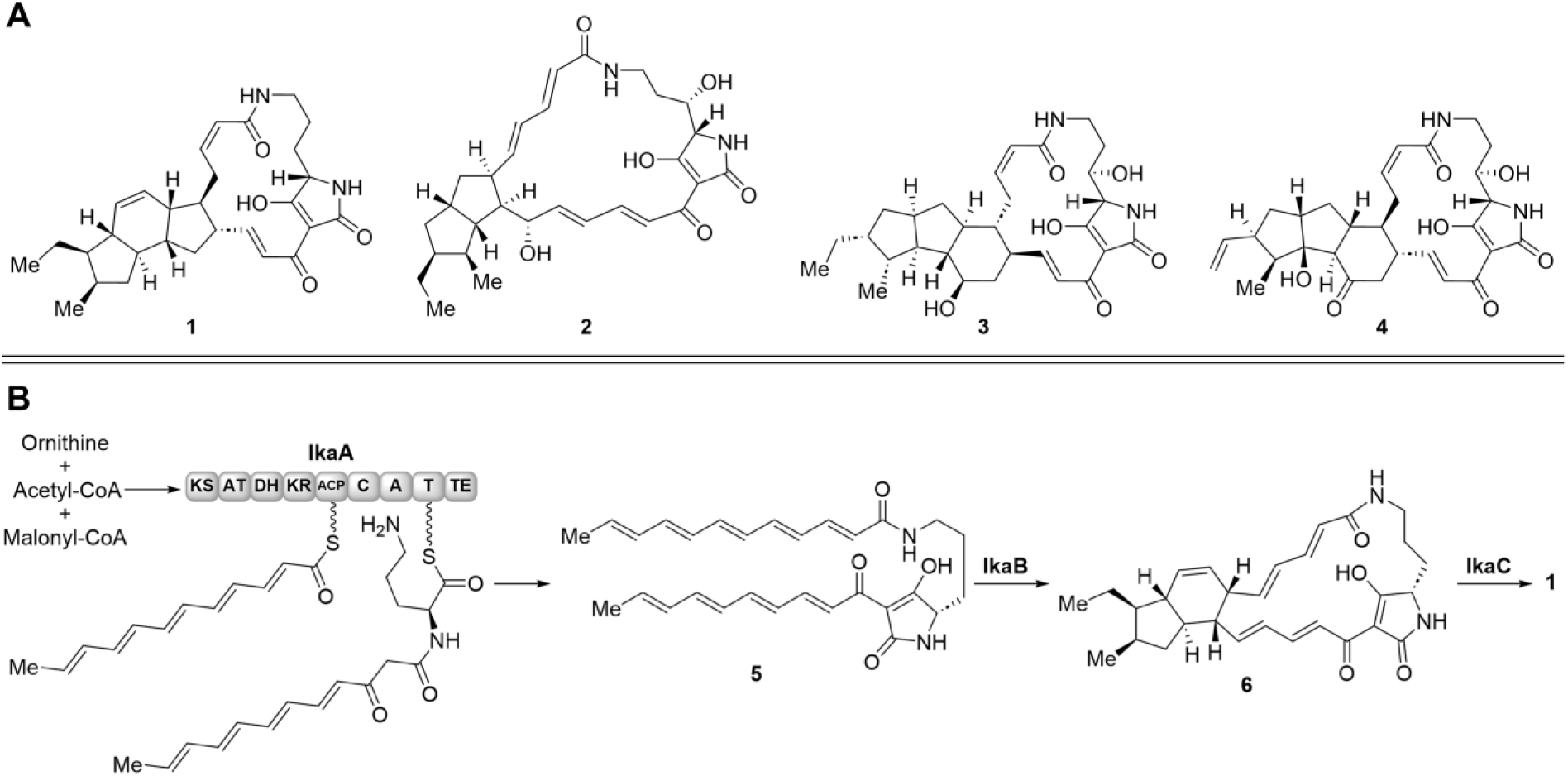
Examples of PoTeMs and their biosynthesis. **A.** Prominent PoTeM examples ikarugamycin (**1**), alteramide A (**2**), HSAF (**3**), and frontalamide A (**4**). **B**. Biosynthesis of **1**: the common intermediate lysobacterene A (**5**) derived of the iPKS/NPRS IkaA is cyclized in a stepwise manner to bicyclic **6** by IkaB and to the final 5-6-5-cyclization pattern in **1** by IkaC.

The general organization of PoTeM biosynthetic gene clusters (BGCs) is highly conserved. The central part is a hybrid iPKS/NRPS that generates lysobacterene A (**5**), the common tetramic acid precursor of all PoTeMs.^1^ Downstream of this core biosynthetic machinery, flavin-dependent phytoene dehydrogenase homologs (PhyDHs) and nicotinamide-cofactor-dependent alcohol dehydrogenase (ADH) homologs are located.^1^ These catalyze formation of the polycyclic ring system typical to PoTeMs. For example, in the biosynthesis of **1**, the PhyDH IkaB catalyzes formation of the two outer rings (one of which putatively formed by a Diels-Alder reaction), while the ADH homolog IkaC installs the inner ring (Figure 1b). In addition, several PoTeM BGCs encode further modifying enzymes, such as cytochrome P450 enzymes (e.g., *ikaD, cftA*, and *ftdF*)^7^and/or a PoTeM hydroxylases (e.g., *SD, ftdA*)^8^for the (oxidative) decoration of the PoTeM core structure.

With the growing knowledge on PoTeM biosynthesis,^1, 5, 6, 7, 9, 10, 11, 12, 13, 14, 15, 16, 17, 18, 19, 20, 21, 22, 23, 24^ targeted approaches to directly identify new PoTeMs were established. Early approaches employed degenerate primers targeting the iPKS/NRPS system to screen unsequenced bacterial strains for the presence of PoTeM BGCs, which led, i.a., to the identification of the BGC encoding **1**^12^ as well as to the discovery of the clifednamides and their pathway.^25^ Most recent work applied genome mining on sequenced genomes, for example leading to the discovery of pseudoamides A−C from *Pseudoalteromonas elyakovii* ATCC 700519,^20^ sahamides A−F from *Saccharopolyspora hirsuta* DSM 44795,^21^ koyanamide A from *S. koyangensis* SCSIO 5802,^26^ clifednamides D−J from *Kitasatospora* sp. S023,^27^ somamycins A−D from *S. somaliensis* SCSIO ZH66,^28^ combamides A−D from *Streptomyces* sp. S10,^29^ pactamides A−F from *S. pactum* SCSIO 02999,^17^ compound D from *S. griseus* IFO 13350,^30^, and others.^18, 31^ These approaches applied, i.a., promoter engineering and heterologous expression to activate silent BGCs.

The increasing number of sequenced genomes and their analysis with genome mining tools such as AntiSMASH^32^ reveals an unexpectedly high abundance of PoTeM BGCs in both, Gram-positive and Gram-negative bacteria.^1^ Since many of these pathways are still unstudied and remain silent during cultivation of the natural producer under standard laboratory expression conditions, we aimed at developing a plug-and-play platform for heterologous expression to facilitate the production of PoTeMs. The main goals were to generate a platform that (1) is applicable to all PoTeMs, (2) enables the production of PoTeMs in sufficiently high yields, (3) enables a fast and effective cloning, and (4) reliably expresses all genes of interest, particularly focusing on PoTeM tailoring genes.

## Methods

### Reagents, primers, and bacterial strains

The listed commercial materials were purchased from the following manufacturers: LB medium, TB medium, CASO bouillon, soluble starch, nalidixic acid (NA), kanamycin sulfate (kan), chloramphenicol (cam), dipotassium phosphate (K_2_HPO_4_), tryptone, yeast extract, mannitol, and agar from Carl Roth (Karlsruhe, Germany); SERVA DNA stain Clear G from Serva (Heidelberg, Germany); dry magnesium sulfate (MgSO_4_), sodium chloride (NaCl), ammonium sulfate ((NH_4_)_2_SO_4_), calcium carbonate (CaCO_3_), iron sulfate (FeSO_4_ × 7 H_2_O), manganese chloride (MnCl_2_ × 4 H_2_O), zinc sulfate (ZnSO_4_ × 7 H_2_O) from Grüssing (Filsum, Germany); apramycin sulfate (apra) from Glentham life science (Wiltshire, UK); PeqGOLD Plasmid Miniprep Kit I (C-Line), and chemical solvents (methanol, ethylacetate, acetonitrile) from VWR (Darmstadt, Germany); deoxynucleotides (dNTPs), Q5 High-Fidelity DNA polymerase, Antarctic phosphatase, restriction enzyme (if available high fidelity), NEBuilder^®^ HiFi DNA Assembly Cloning Kit, T-4 DNA polymerase, and Monarch^®^ DNA Gel Extraction Kit from NEB (Frankfurt am Main, Germany); T4-DNA ligase and PCR Purification Kit from Jena Bioscience (Jena, Germany). All kits and enzymes were used according to the manufacturer’s recommended protocol unless otherwise stated. Primers were purchased from Sigma Aldrich (Darmstadt, Germany) at quality “deprotected and desalted”. All used bacterial strains are listed in Table S6.

### Primer design and PCR

Primers were designed using the NEBuilder assembly tool and evaluated using Oligo Calc. Gibson homology arms were composed of at least 18 nt with a calculated melting temperature of ≥ 50 °C. HiFi DNA assemblies were simulated using SnapGene, in this way excluding multiple binding sites of the primers. Primers used in this study are listed in Table S7. PCRs for insert generation were performed using Q5 High Fidelity DNA Polymerase (NEB, Germany). Colony-screening-PCR was performed using Taq DNA polymerase (NEB, Germany). Clones were picked and resuspended in 50 µL dH_2_O. 5 µL of cell suspension were directly used as template DNA. Cycling was conducted in T100 Thermal Cycler (Biorad, Germany) or a LifeECO Thermal Cycler (Biozym, Germany).

### DNA assembly

All DNA assembly methods were conducted in a 20 µL reaction batch containing 0.02 pmol vector DNA and 0.1 pmol insert DNA. For conventional cloning, 1× T4-ligase reaction buffer and 0.12 U T4-ligase were added and incubated at 16 °C over night. Ligation reactions were heat-inactivated at 65 °C for 10 min. For HiFi-DNA assembly 1x HiFi DNA assembly master mix was added and incubated at 50 °C for 1 h. For SLIC 1× NEBBuffer 2.1 and 1 µL T4-polymerase were added on ice, incubated 10 min at room temperature and stored on ice for 10 min.

### Cloning strategy

Expression plasmids were obtained in four individual cloning steps based on the Direct Pathway Cloning (DiPaC) approach (Figure S2).^33, 34^ In a first step, the iPKS/NRPS gene *ikaA* was added to the expression plasmid pSET152-ermE. The vector was digested with StuI and *ikaA* (9.4 kb) was amplified by PCR from pSET152-ermE*::ikaABC* adding homology arms. The two DNA fragments were assembled using HiFi DNA assembly. The different tested promoters were ordered as synthetic genes from Eurofins Genomics in a pEX-A2 vector (Table S7). The promoters were excised from the plasmids by digestion with StuI and XbaI, dephosphorylated, and subsequently ligated into pSET152-ermE::*ikaA*, which in turn was linearized with StuI and XbaI, to obtain the basic expression constructs for the new plug-and-play system.

For this study, the two cyclizing genes from the ikarugamycin BGC, *ikaB* and *ikaC*, and three different tailoring genes (*ikaD, ptmD*, and *cftA*) were added to the plug-and-play platform. Therefore, *ikaBC* was amplified as one piece (2.9 kb) by PCR simultaneously adding homology arms and assembled with the StuI digested and dephosphorylated basic expression constructs using HiFi DNA assembly. The three modifying genes were amplified by PCR using either gDNA (*ikaD*) or synthesized genes (*ptmD* and *cftA*) as templates. Genes were introduced by digestion of the expression vectors and inserts with XbaI, dephosphorylation, and ligation to obtain the final expression constructs for this study. *CftA* was amplified with homology arms and integrated using SLIC.^35^ All cloning steps were verified by analytical restriction digests and sequencing (cf. ESI, chapter 8).

### Heterologous expression in *Streptomyces* sp. and extraction of PoTeMs

The expression vectors were individually integrated into *S. albus* DSM 40313, *S. lividans* TK24, and *S. coelicolor* M1154 using intergenetic conjugation with *E. coli* ET12567/pUZ8002. Therefore, expression constructs were transformed into *E. coli* ET12567/pUZ8002 by electroporation. PCR-verified clones were grown in LB medium containing apra, cam, and kan (37 °C, 200 rpm) until an OD_600_ of 0.4–0.6 was reached. Cells were washed twice with 10 mL ice-cold LB without antibiotic (4000 rpm, 5 min) and resuspended in 500 µL LB without antibiotic. Spores were resuspended in 500 µL 2xYT (16 g/L tryptone, 10 g/L yeast extract, and 5 g/L NaCl, pH 7.0) and heat-activated at 50 °C for 10 min. Cells and spores were combined, harvested (4000 rpm, 2 min), plated on MS agar (20 g/L mannitol, 20 g/L soya flour, 20 g/L agar) supplemented with 10 mM MgCl_2_ and 60 mM CaCl_2_, and cultivated at 30 °C. After 16–20 h, conjugation plates were overlaid with 1 mg NA and 1 mg apra. Colonies were obtained after 4–7 days (30 °C) and verified by colony-screening-PCR.

*Streptomyces* sp. were first grown in CASO supplemented with apra and NA for 3–5 days (28 °C, 200 rpm). After upscaling to a 20 mL CASO culture, 5 mL of a dense culture were used to inoculate 50 mL ISP-4 (10 g/L soluble starch, 1 g/L K_2_PO_4_, 1 g/L MgSO_4_, 1 g/L NaCl, 2 g/L (NH_4_)_2_SO_4_, 2 g/L CaCO_3_, 1 mg/L FeSO_4_ × 7 H_2_O, 1 mg/L MnCl_2_ × 4H_2_O, 1 mg/L ZnSO_4_ × 7 H_2_O) without antibiotics. The main culture was harvested (6000 rpm, 10 min) after 7 days incubation (28 °C and 200 rpm). Pellet and supernatant were extracted individually. The pH of the supernatant was set to 5 using 1 M HCl_aq_ and the samples extracted three times with ethyl acetate. The combined organic phases were dried over MgSO_4_, filtered, and solvent was removed *in vacuo*. The cells were resuspended in 20 mL methanol/acetone (1:1), sonicated for 30 min in a sonicating bath, and centrifuged (10 min, 6000 rpm). The supernatant was dried *in vacuo*.

### HPLC-MS analysis

Organic extracts were analyzed using a computer controlled Jasco HPLC System composed of a MD-2010 Plus Multiwavelength detector, a DG-2080-53 3-Line Degasser, two PU-2086 Plus Intelligent Pumps, a AS-2055 Plus Intelligent Sampler, a MIKA 1000 dynamic mixing chamber, a 100 µL sample loop (Portmann Instruments AG Biel-Benken), and a LCNetII/ADC. For LC-MS measurements, this system was coupled to an Expression LCMS instrument (Advion) containing a single-quadrupole mass analyzer used in combination with a N118LA nitrogen generator (Peak Scientific) and an RV12 high vacuum pump (Edwards). For chromatographic separation a Eurosphere C8 column with precolumn (Knauer 10XE084E2J, 100 × 3 mm, 100-5 C8 A) was utilized. H_2_O (A) and acetonitrile (B), both supplemented with 0.05% TFA, with a flowrate of 1 mL/min were used as solvents. The gradient was as followed: 0–2 min 100% A, 2–10 min 100–55% A, 10–20 min 55% A, 20–24 min 0% A, 24–28 min 100% A.

### HR-MS

For high resolution mass spectrometry and MS/MS measurements after HPLC separation, a Bruker UHPLC consisting of an Elute autosampler and a HPG 1300 pump was used. This was coupled to an Impact II mass spectrometer with ESI source and Q-TOF mass analyzer manufactured by Bruker. The following parameters were used: solvents: A = H_2_O + 0.05% formic acid (FA), B = ACN + 0.05% FA; separation method: 0–2 min: 95% A, 2–25 min: 95–5% A, 25–28 min: 5% A, 28–30 min: 95% A; flow rate: 0.3 mL/min; column: Intensity Solo 2 C18, 100 × 2.1 mm (in column oven: 40 °C). For MS/MS, auto-MS/MS mode with 20–50 eV collision energy (N_2_) was used. The system was controlled by Bruker Compass^®^ HyStar software; analysis was conducted with Bruker Compass^®^ Data Analysis software.

### Compound Purification

The PoTeMs were purified from raw extracts by semi-preparative HPLC, either using a system made by Jasco (composed of an UV-1575 Intelligent UV/VIS detector, two PU-2086 Intelligent HPLC pumps, a MIKA 1000 dynamic mixing chamber, a 5000 µL sample loop (Portman Instruments AG Biel-Benken), a LC-NetII/ADC, and a Rheodyne injection valve; Galaxie software) or from Knauer (consisting of a degasser, a P 6.1L pump, a 1000 µL injection port, and a Smartline UV detector 2500; ClarityChrom) with detection at 220 nm. H_2_O (A) and acetonitrile (B), both supplemented with 0.05% TFA, were used as solvents. Individual separation conditions were: Capsimycin G (8): column: Eurospher II, 100-5 C18A, 250 × 16 mm; gradient: 0–2 min: 75% A, 2–22 min: 75–25% A, 22–32 min: 0% A; flow rate 5 mL/min. Butremycin (9): column: Eurospher II, 100-5 C8A, 250 × 8 mm; gradient: 0–2 min: 95% A, 2–13 min: 95–50% A, 13–20 min: 50% A, 20–24 min: 0% A, 24–28 min: 95% A; flow rate 3 mL/min. Clifednamides A (10) and C (11): column Eurospher II 100-5 C18A, 250 × 8 mm; gradient: 0–2 min: 75% A, 2–37 min: 75–40% A, 37–50 min: 0% A; flow rate 5 mL/min.

### NMR

Nuclear Magnetic Resonance Spectra (NMR) were recorded on Bruker AVANCE II 300, ASCEND 500 and AVANCE III 600 spectrometers at ambient temperature. The chemical shifts are given in δ-values (ppm) relative to TMS (^1^H, ^13^C). ^1^H and ^13^C spectra were referenced internally using the residual proton resonances (DMSO-*d*_6_: δ_H_ = 2.50 ppm, δ_C_ = 39.52 ppm, pyridine-*d*_5_: δ_H_ = 7.22 ppm, δ_C_ = 123.87 ppm;). The coupling constants *J* are given in Hertz [Hz] and determined assuming first-order spin-spin coupling. The following abbreviations were used for the allocation of signal multiplicities: s – singlet, bs – broad singlet, d – doublet, bd – broad doublet, t – triplet, bt – broad triplet, q – quartet, qnt – quintet, sxt – sextet, spt – septet, m – multiplet, or any combination thereof.

## Results

### Establishing a plug-and-play system for PoTeM expression

To establish an efficient recombinant expression platform, we decided to use *Streptomyces* sp. as heterologous hosts, since the majority of PoTeM BGCs natively derive from this species.^1^ As all currently known PoTeM biosynthetic pathways rely on the prototypical iPKS/NRPS system to provide the common precursor molecule **5**, we installed *ikaA* from *Streptomyces* sp. Tü6239^12, 36^ under the control of the strong constitute promoter ermE^*37^ (P1) into a pSET152 vector backbone by Direct Pathway Cloning (DiPaC)^34^ to form the basis of the plug-and-play platform (Figure 2A). We selected *ikaA* as previous studies had shown successful heterologous expression of **1** in *E. coli*^12^ and *Streptomyces*^14^ using the *ikaABC* gene cassette, proving that the precursor **5** should be produced in sufficient amounts. The resulting construct was tested by expression in *S. albus* DSM 40313 and the production of **5** was confirmed by HPLC-MS (Figure S1). Downstream of *ikaA*, we installed two unique restriction sites (StuI and XbaI) flanking a second promoter (P2). The StuI site can be utilized to integrate genes encoding cyclization enzymes of choice. For establishing our system, we exemplarily used the ikarugamycin tailoring genes *ikaBC*, which were integrated by HiFi DNA assembly.^38^ The XbaI site was designed to readily integrate genes involved for late-stage PoTeM functionalization by ligation cloning or SLIC.^35^ Since promoter activities are moderately studied in *Streptomyces*,^39^ we integrated eight different promoters to determine optimal expression conditions for individual genes (Figure 2). These included gapdhP(EL), which is located upstream of the gapdh operon in *Eggerthella lenta*, and rpsLP(XC), which is located upstream of the operon coding for the 30S ribosomal proteins S12 and S7 and elongation factors in *Xylanimonas cellulosilytica*, since they were reported as most active ones among a set of various promoters derived from house-keeping genes of different organisms.^40^ KasOP^*^ was previously generated by targeted and random mutagenesis of a promoter positioned upstream of a PKS in *S. coelicolor*.^41^ It showed an upregulated expression of up to five times compared to ermE.^41^ The phage *I19* promoter SF14P was isolated from a *Streptomyces* strain and also showed a high expression level.^42^ The last four promoters P-2, P-6, P-15, and P-31 were discovered in a large study that compared non-transcribed intergenic regions in *S. albus* J1074 and showed significant activating effect.^43^

**Figure 2.**
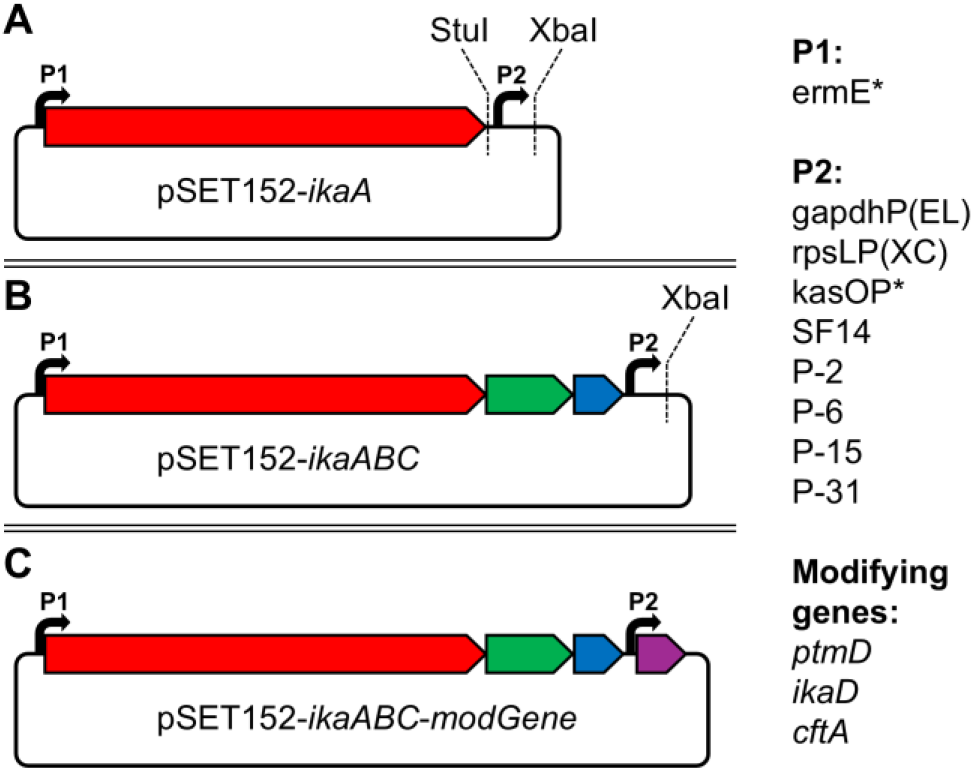
Plug-and-play system for the heterologous expression of PoTeMs. The system is based on a pSET152 vector and designed for *Streptomyces* expression. **A.** The basic expression vector contains the iPKS/NRPS *ikaA* under control of an ermE^*^ promoter (P1). **B**. The expression vector contains all essential genes *ikaABC* for the biosynthesis of **1**. The construct can be used to investigate the influence of modifying enzymes on **1. C**. Plug-and-play system containing either *ptmD, ikaD*, or *cftA* under the individual control of eight different, strong constitutive promoters (P2). Red: *ikaA*, green: *ikaB*, blue: *ikaC*, and purple tailoring genes *ptmD*/*ikaD*/*cftA*.

### Bispecific monooxygenase IkaD

As we focused on enabling investigations on late-stage tailoring enzymes in this study, we integrated *ikaBC* into the StuI site by ligation cloning (Figure 2B, Figure S2), thus delivering the ikarugamycin core structure **1** in high yields upon heterologous expression in *Streptomyces* (Figure S1). This construct was further used to probe the influence of different modifying genes acting on PoTeM core structures, exemplarily for **1**, under the control of the eight selected constitutive promoters. As proof-of-concept, *ikaD* was chosen as the first modifying enzyme. This bispecific P450 monooxygenase was initially characterized from the gene cluster of **1** from *S. xiamenensis* 318,^44^ and was intensively investigated *in vivo* and *in vitro*. It adds an epoxide at the C-13/14 of its native substrate **1**, resulting in capsimycin B (**7**).^7, 44^ Additionally, IkaD hydroxylates **7** at C-29 resulting in capsimycin G (**8**).^7, 44^ The activity of IkaD acting on the 5-5-cyclic pattern of combamides was also elucidated *in vivo* and *in vitro*.^16, 23^

*IkaD* was added to the expression platform containing *ikaABC* under the control of the eight different second promoters. The presence of the restriction site (XbaI) enabled a fast insertion by classical ligation cloning. The constructs were expressed in three different *Streptomyces* strains (*S. albus* DSM 40313, *S. lividans* TK24, and *S. coelicolor* M1154) under the conditions resulting in highest yields of **1** in previous expression experiments (see above). For every promoter, biological triplicates were cultured, extracted, and analyzed by LC-MS for PoTeM quantification. In *S. albus*, **1** was almost completely converted to the two oxidatively modified compounds **7** and **8** (Figure 3E). The major product in the cells was the double-modified ikarugamycin derivative **8** (*m/z* 511.2835), whereas in the medium, the single-modified intermediate **7** (*m/z* 495.2850) was also present in comparable amounts. Additionally, trace amounts of further PoTeM decomposition products were detected in the extract of the medium. These compounds were generated under the acidic extraction conditions, which led to a nucleophilic epoxide-opening by methanol or water.^44^

**Figure 3.**
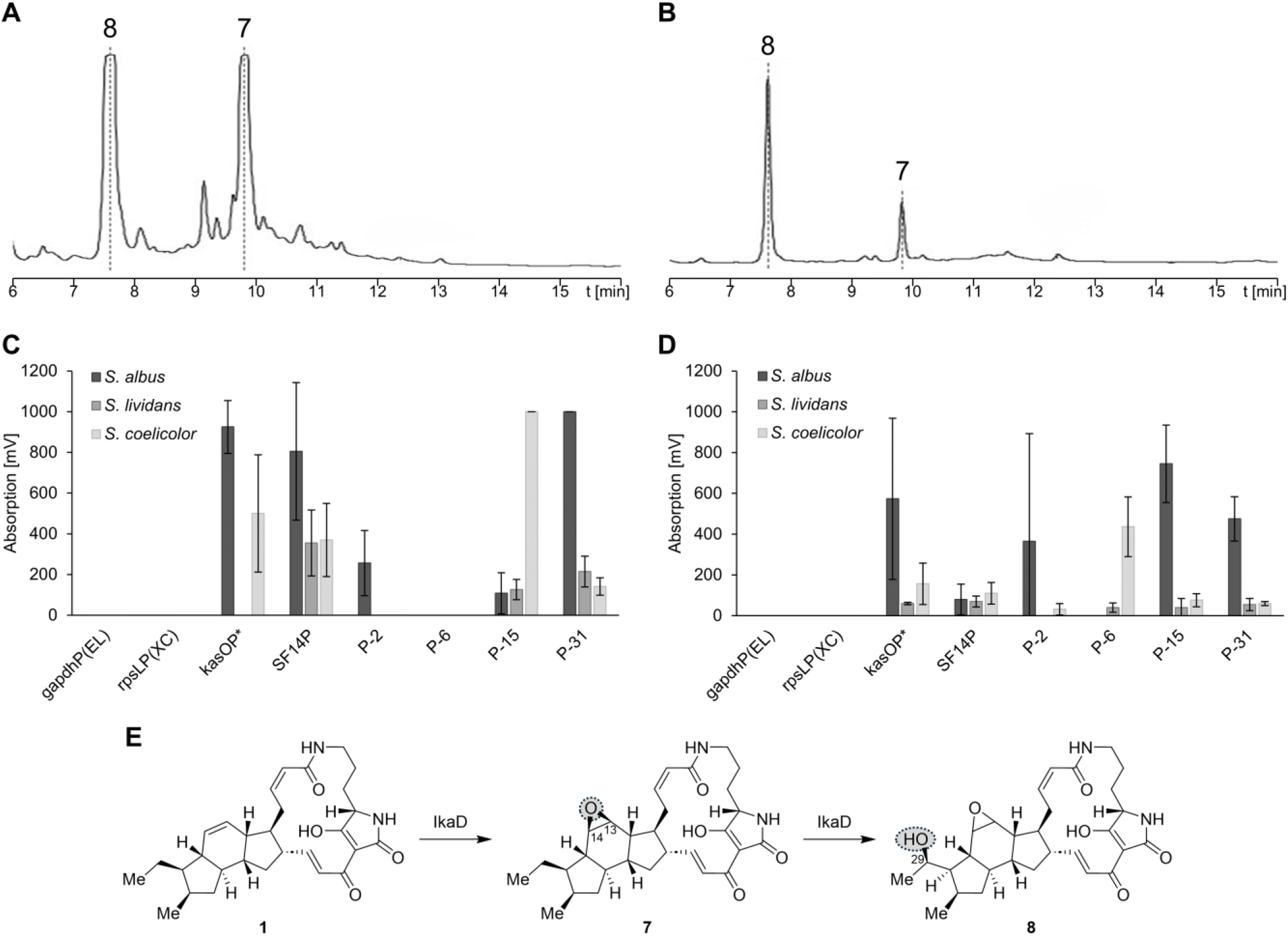
Conversion of **1** to **7**/**8** by IkaD using different promoters and *Streptomyces* expression hosts. HPLC-UV (220 nm) chromatogram of the extracts of **A.** the medium and **B**. the cells, exemplarily shown for expression with *S. albus* DSM 40313 equipped with pSET152-ermE-*ikaABC*-kasOP^*^-*ikaD*. Amount of capsimycin G (**8**) in **C**. the medium and **D**. the cells, with tailoring gene expression under control of eight different promoters in three different stains: *S. albus* DSM 40313 (black), *S. lividans* TK24 (dark grey), and *S. coelicolor* M1154 (light grey). **E**. Stepwise modification of **1** to **7** and **8** by IkaD.

The yield of **8** significantly depended on the used strain and promoter (Figure 3CD). Generally, the expression worked best in *S. albus* and barely in *S. lividans*. In *S. albus*, the utilization of kasOP^*^ (Figure 3AB), SF14P, and P-31 as second promoter resulted in the highest production of **8**, which was preferably located in the cells. The use of promoter P-15 led to a high yield of **8** in the cells, but the PoTeM was barely transported to the medium. Promoter P-2 still enabled production of **8**, but in low amounts. With the remaining promoters (gapdhP(EL), rpsLP(XC), and P-6) no conversion of **1** was observed, which indicates that these were completely inactive. The expression in *S. coelicolor* resulted in significantly lower yields, but again kasOP^*^ and SF14P belonged to the best-performing promoters. The best promoter in *S. coelicolor* was P-15. In *S. lividans*, PoTeM expression was generally very low. The most active promoters were SF14P and P-31.

For structural validation, large scale fermentation (2 L) was performed with *S. albus* and the well-performing expression construct containing the kasOP^*^ promoter. Following compound isolation from the fermentation cell pellet, the structure of compound **8** was validated by 1D and 2D NMR (cf. ESI, Table S2, Fig. S14–17). Consistent with literature, IkaD introduces two modifications to **1**: an epoxide at the position of the former double bond (C-13/14) and a hydroxy group at C-29 (Figure 3E).^7, 44^ In addition, based on the HRMS data and the higher retention time, **7** was unambiguously identified as the single-modified product of IkaD.

### PoTeM hydroxylase PtmD

We next investigated a modifying enzyme that naturally acts on 5-5-6-cyclic PoTeMs to determine whether it can also be applied for modifications on 5-6-5-cyclic **1**. We selected *ptmD* from the pactamide gene cluster, described as a hydroxylase whose regioselectivity has not been fully characterized.^17,18^ The *ptmD* gene was efficiently inserted downstream of the eight different second promoters using ligation cloning and the expression was again performed in the three different *Streptomyces* strains. The analysis of the extracts revealed that **1** remained the primary product under all tested conditions. However, a new PoTeM (**9**) with a shorter retention time and an *m/z* 495.2850 was also detected, consistent with a hydroxylated derivative of **1** ([M+H]^+^ calculated *m/z* 495.2853; Figure 4AB). As observed for IkaD above, *S. albus* yielded the highest amounts of PoTeMs (Figure 4CD). The conversion of **1** in the two other strains was even lower than during experiments with IkaD. Interestingly, the promoters gapdhP(EL) and rpsLP(XC), which were essentially inactive with IkaD, yielded the highest conversion rates for PtmD. Except for P-6 and P-15, all other promoters also yielded acceptable conversion rates. For compound isolation, the fermentation of the construct containing the gapdhP(EL) promoter was scaled up (2 L).

**Figure 4.**
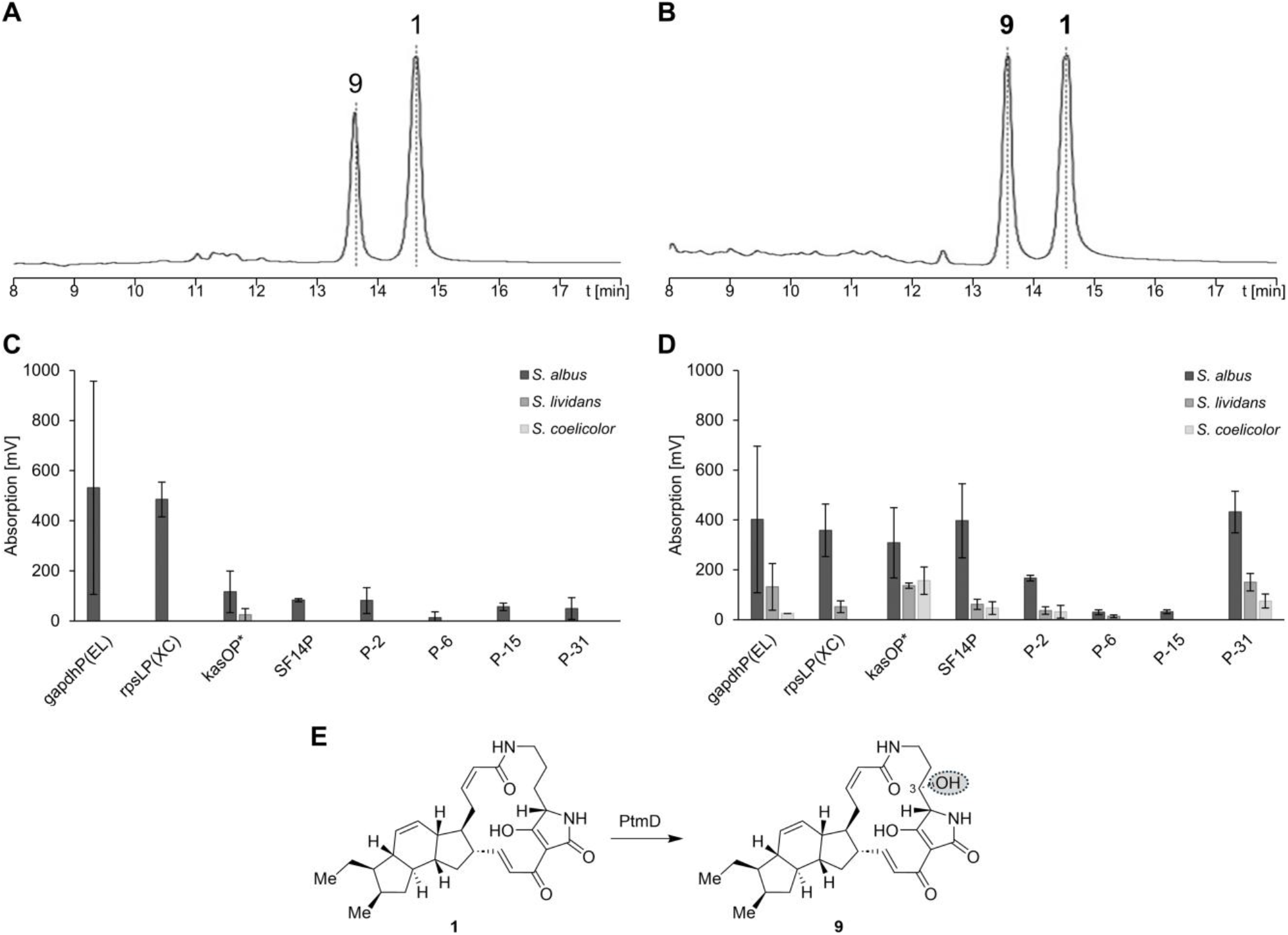
Conversion of **1** to **9** by PtmD using different promoters and *Streptomyces* expression hosts. HPLC-UV (220 nm) chromatogram of the extracts of **A.** the medium and **B**. the cells, exemplarily shown for expression with *S. albus* DSM 40313 equipped with pSET152-ermE-*ikaABC*-gapdhP(EL)-*ptmD*. Amount of butremycin (**9**) in **C**. the medium and **D**. the cells, with tailoring gene expression under control of eight different promoters in three different strains: *S. albus* DSM 40313 (black), *S. lividans* TK24 (dark grey), and *S. coelicolor* M1154 (light grey). **E**. Modification of **1** to **9** by PtmD.

Initially, *ptmD* was presumed to function as a hydroxylase at position C-31 of pactamide A.^17^ However, it was later hypothesized to act as a PoTeM hydroxylase, adding a hydroxyl group at the C-3 position.^18^ This is also supported by its sequence similarity to PoTeM hydroxylases (over 50% amino acid identity) and its location upstream of the iPKS/NRPS, where commonly PoTeM hydroxylases can be found.^17^ However, no detailed investigation had been conducted so far. A comparison of 1D NMR data of **9** and **1** revealed differences in the region of the ornithine-derived molecular portion. The signals of the two hydrogens at C-3, which appear in the region between d_H_ 1.69–1.88 ppm in **1**, were missing. Instead, new signals at 3.80 and 5.09 ppm appeared. Together with 2D NMR analysis (cf. ESI, Fig. S18–21), the hydroxyl group was localized at C-3 (Figure S4). Thus, PtmD was confirmed to act as a PoTeM hydroxylase that converts **1** to butremycin (**9**; Figure 4E).^8^ Furthermore, it was shown that PtmD, which derives from a BGC encoding a 5-5-6-ring PoTeM, can also function on the 5-6-5-carbacyclic system.

### Applying the plug-and-play system for PoTeM modification

To apply the gained information about our plug-and-play system, we tested a third modifying gene found in PoTeM BGCs. *CftA* derives from the clifednamide gene cluster and is a P450 monooxygenase.^45^ For the investigation of its function, we only applied a single expression condition (Figure 5). We used *S. albus*, which was the most productive strain during both previous screenings, and used gapdhP(EL) as second promoter, since this construct yielded the highest conversion with PtmD. The analysis of the corresponding cell extracts showed complete conversion of **1** to two new products **10** (*m/z* 493.2692) and **11** (*m/z* 509.2631). Due to the high expression level, both compounds were readily isolated, and their structures were elucidated by 1D and 2D NMR (cf. ESI, Table S4/5, Figure S22–S31). In both compounds, a ketone was located at the ethyl moiety at C-29 (Scheme 1). Compound **11** additionally possessed a neighboring hydroxyl group at C-30. Bifunctionality and the regioselectivity of CftA, resulting in clifednamide A (**10**) and clifednamide C (**11**), were thus in accordance with previous studies *in vivo*^27, 46^ and *in vitro*.^7, 46^

**Figure 5.**
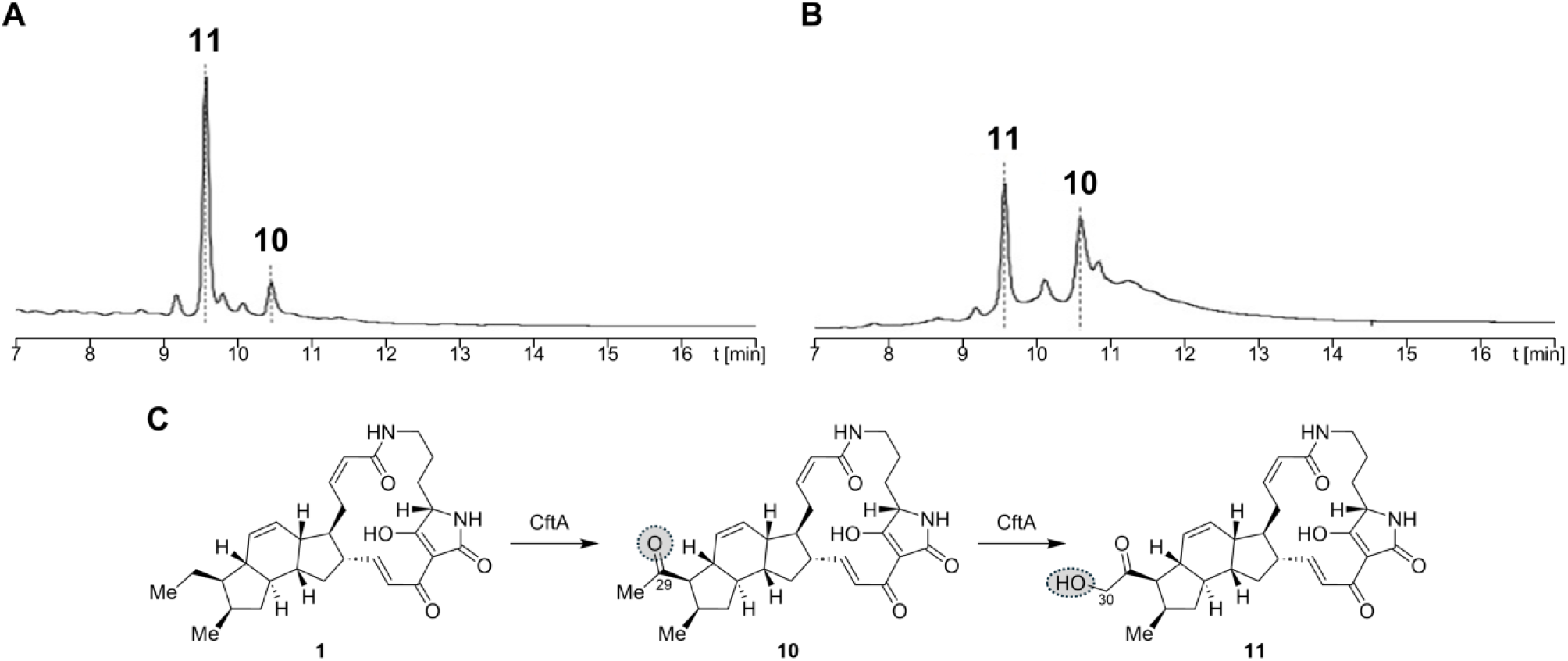
Conversion of **1** to **10** and **11** by CftA. HPLC-UV (220 nm) chromatogram of the extracts of **A.** medium and **B**. the cells from *S. albus* DSM 40313. **1** was efficiently converted by CftA to clifednamide A (**10**) and clifednamide C (**11**) by using gapdhP(EL) as second promoter. **C**. Stepwise modification of **1** to **10** and **11** by CftA.

## Discussion

To discover the full diversity of PoTeMs, heterologous expression became irreplaceable, since many BGCs remain silent under laboratory conditions in their native hosts. To set the stage for investigations of new PoTeM BGCs, particularly to study functions of tailoring enzymes, our generated plug-and-play system contains *ikaA* for a reliable production of the uniform PoTeM precursor molecule **5**. Oxidoreductases from all kinds of PoTeM BGCs can be easily introduced and tested for their activity *in vivo*, as exemplarily shown for production of **1** using *ikaBC*. Furthermore, modifying enzymes can be introduced to this vector downstream of a second promoter. Thereby, the functionality of the bifunctional monooxygenases IkaD and CftA were confirmed and the function of PtmD was reassigned as C-3 PoTeM hydroxylase (Scheme 1). Previous studies had already shown that it can be beneficial to replace the native iPKS/NRPS by a well-studied homologous version to ensure production of a sufficient amount of the central precursor **5**.^20, 21, 23^ Within this work, we additionally show that different promoters controlling genes encoding tailoring enzymes can dramatically influence modified PoTeM production yields. Besides the broadly used kasOP^*^, P-2 and P-31 generally yielded good gene expression of PoTeM modifying genes. Surprisingly, we observed differences between the expression of the monooxygenase *ikaD* and the hydroxylase *ptmD*. For *ikaD*, also the promoters gapdhP(EL), rpsLP(XC), SF14P, and P-15 yielded high conversion of **1** to **7** and **8** (Scheme 1). Compound **1** was almost completely converted by the expression of the monooxygenases through our system. Furthermore, the plug-and-play system was successfully used in different *Streptmyces* strains. *S. albus* DSM40313 was found to be the most productive strain, but *S. coelicolor* M1154 also yielded decent amounts and had slightly less side-products, which potentially facilitates product isolation from complicated product mixtures.

**Scheme 1.**
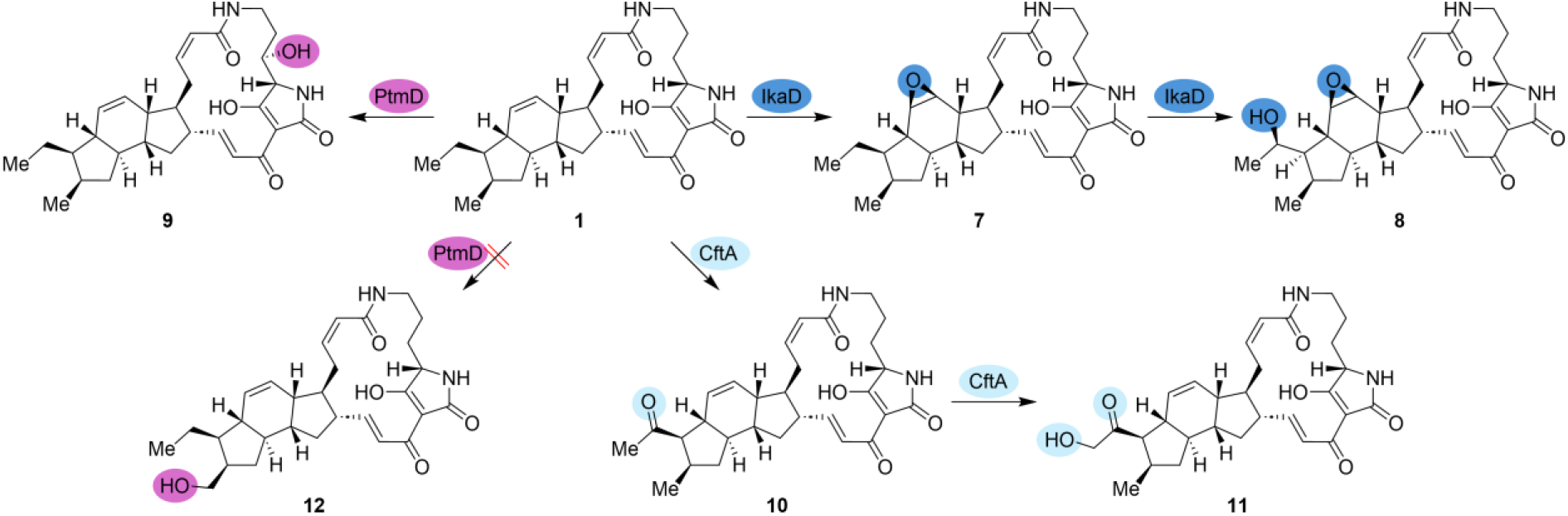
Conversion of 1 by PtmD, IkaD, and CftA. The natural products butremycin (**9**), capsimycin B (**7**) and capsimycin G (**8**), as well as clifednamide A (**10**) and clifednamide C (**11**), were produced by the three modifying enzymes, respectively. Hydroxylation at position C-31 by PtmD was not observed.

## Conclusions

Overall, we established a genetic plug-and-play system that readily facilitated the production of oxidatively modified PoTeM analogs by late-stage functionalization of **1** using IkaD (products **7** and **8**), CftA (**10** and **11**), and PtmD (**9**) (Scheme 1). This system and our insights into production levels in three different *Streptomyces* sp. host strains combined with several constitutive promoters can now be further applied to the investigation of new PoTeM BGCs and/or modifying enzymes, making novel PoTeM derivatives accessible for biological activity screenings.

## Supporting information

ESI File

## Declarations

### Ethics approval and consent to participate

Not applicable.

### Consent for publication

Not applicable.

### Availability of data and materials

All data generated or analyzed in this study are included in this published article [and its supplementary information files]. All raw data files are available from the corresponding author on request.

### Authors’ contributions

AG performed the cloning, expression, and analysis of the different expression constructs. SS performed upscaling experiments, compound isolation, and structure elucidation. ME conducted structural data analysis and revision, as well as discussion of results. TAMG developed the research program, secured funding, and supervised the project. All authors contributed to writing the manuscript.

## Acknowledgements

The authors acknowledge Dr. Tilo Lübken (Chair of Organic Chemistry I, TU Dresden) for his support with NMR analyses of functionalized PoTeMs.

## References

1. Harper, C. P.; Day, A.; Tsingos, M.; Ding, E.; Zeng, E.; Stumpf, S. D.; Qi, Y.; Robinson, A.; Greif, J.; Blodgett, J. A., Critical analysis of polycyclic tetramate macrolactam biosynthetic gene cluster phylogeny and functional diversity. Appl. Environ. Microbiol. 2024, 90, e00600–24.

2. Jomon, K.; Kuroda, Y.; Ajisaka, M.; Sakai, H., A new antibiotic, ikarugamycin. J. Antibiot. 1972, 25, 271–280.

3. Shigemori, H.; Bae, M. A.; Yazawa, K.; Sasaki, T.; Kobayashi, J., Alteramide A, a new tetracyclic alkaloid from a bacterium Alteromonas sp. associated with the marine sponge Halichondria okadai. J. Org. Chem. 1992, 57, 4317–4320.

4. Sullivan, R.; Holtman, M.; Zylstra, G.; White Jr, J.; Kobayashi, D. Y., Taxonomic positioning of two biological control agents for plant diseases as Lysobacter enzymogenes based on phylogenetic analysis of 16S rDNA, fatty acid composition and phenotypic characteristics. J. Appl. Microbiol. 2003, 94, 1079–1086.

5. Yu, F.; Zaleta-Rivera, K.; Zhu, X.; Huffman, J.; Millet, J. C.; Harris, S. D.; Yuen, G.; Li, X.-C.; Du, L., Structure and biosynthesis of heat-stable antifungal factor (HSAF), a broad-spectrum antimycotic with a novel mode of action. Antimicrob. Agents Chemother. 2007, 51, 64–72.

6. Blodgett, J. A.; Oh, D. C.; Cao, S.; Currie, C. R.; Kolter, R.; Clardy, J., Common biosynthetic origins for polycyclic tetramate macrolactams from phylogenetically diverse bacteria. Proc. Natl. Acad. Sci. U. S. A. 2010, 107, 11692–11697.

7. Jiang, P.; Jin, H.; Zhang, G.; Zhang, W.; Liu, W.; Zhu, Y.; Zhang, C.; Zhang, L., A mechanistic understanding of the distinct regio-and chemoselectivity of multifunctional P450s by structural comparison of IkaD and CftA complexed with common substrates. Angew. Chem. 2023, 135, e202310728.

8. Greunke, C.; Antosch, J.; Gulder, T. A. M., Promiscuous hydroxylases for the functionalization of polycyclic tetramate macrolactams - conversion of ikarugamycin to butremycin. Chem. Commun. 2015, 51, 5334–5336.

9. Lou, L.; Qian, G.; Xie, Y.; Hang, J.; Chen, H.; Zaleta-Rivera, K.; Li, Y.; Shen, Y.; Dussault, P. H.; Liu, F.; Du, L. Biosynthesis of HSAF, a tetramic acid-containing macrolactam from Lysobacter enzymogenes. J. Am. Chem. Soc. 2011, 133, 643–645.

10. Li, Y.; Chen, H.; Ding, Y.; Xie, Y.; Wang, H.; Cerny, R. L.; Shen, Y.; Du, L., Iterative assembly of two separate polyketide chains by the same single-module bacterial polyketide synthase in the biosynthesis of HSAF. Angew. Chem. Int. Ed. 2014, 53, 7524–7530.

11. Chen, H.; Du, L., Iterative polyketide biosynthesis by modular polyketide synthases in bacteria. Appl. Microbiol. Biotechnol. 2016, 100, 541–557.

12. Antosch, J.; Schaefers, F.; Gulder, T. A. M., Heterologous reconstitution of ikarugamycin biosynthesis in E. coli. Angew. Chem. Int. Ed. 2014, 53, 3011–3014.

13. Zhang, G.; Zhang, W.; Zhang, Q.; Shi, T.; Ma, L.; Zhu, Y.; Li, S.; Zhang, H.; Zhao, Y.-L.; Shi, R.; Zhang, C. Mechanistic insights into polycycle formation by reductive cyclization in ikarugamycin biosynthesis. Angew. Chem. Int. Ed. 2014, 53, 4840–4844.

14. Greunke, C.; Glöckle, A.; Antosch, J.; Gulder, T. A. M., Biocatalytic total synthesis of ikarugamycin. Angew. Chem. Int. Ed. 2017, 56, 4351–4355.

15. Li, Y.; Wang, H.; Liu, Y.; Jiao, Y.; Li, S.; Shen, Y.; Du, L., Biosynthesis of the polycyclic system in the antifungal HSAF and analogues from Lysobacter enzymogenes. Angew. Chem. Int. Ed. 2018, 57, 6221–6225.

16. Jin, H.; Zhang, W.; Zhang, G.; Zhang, L.; Liu, W.; Zhang, C., Engineered biosynthesis of 5/5/6 type polycyclic tetramate macrolactams in an ikarugamycin (5/6/5 type)-producing chassis. Org. Lett. 2020, 22, 1731–1735.

17. Saha, S.; Zhang, W.; Zhang, G.; Zhu, Y.; Chen, Y.; Liu, W.; Yuan, C.; Zhang, Q.; Zhang, H.; Zhang, L.; Zhang, W.; Zhang, C. Activation and characterization of a cryptic gene cluster reveals a cyclization cascade for polycyclic tetramate macrolactams. Chem. Sci. 2017, 8, 1607–1612.

18. Liu, W.; Zhang, W.; Jin, H.; Zhang, Q.; Chen, Y.; Jiang, X.; Zhang, G.; Zhang, L.; Zhang, W.; She, Z.; Zhang, C. Genome mining of marine-derived Streptomyces sp. SCSIO 40010 leads to cytotoxic new polycyclic tetramate macrolactams. Mar. Drugs 2019, 17, 663.

19. Chen, Y.; Jin, H.; Xiong, W.; Fang, Z.; Sun, W.; Zhu, Y.; Zhang, L.; Zhang, Y.; Zhang, W.; Zhang, C., Discovery of aburatubolactams reveals biosynthetic logic for distinct 5/5-type polycyclic tetramate macrolactams. Org. Lett. 2024, 26, 1677–1682.

20. Li, X.; Liu, Q.; Zou, H.; Luo, J.; Jiao, Y.; Wang, H.; Du, L.; Shen, Y.; Li, Y., Discovery and biosynthesis of pseudoamides reveal enzymatic cyclization of the polyene precursor to 5–5 bicyclic tetramate macrolactams. ACS Catal. 2023, 13, 4760–4767.

21. Zou, H.; Xia, X.; Xu, Q.; Wang, H.; Shen, Y.; Li, Y., Discovery of oxidized polycyclic tetramate macrolactams bearing one or two rings through combinatorial pathway reassembly. Org. Lett. 2022, 24, 6515–6519.

22. Luo, J.; Li, X.; Wang, H.; Du, L.; Shen, Y.; Li, Y., Identification and characterization of the 28-N-methyltransferase involved in HSAF analogue biosynthesis. Biochemistry 2022, 61, 2879–2883.

23. Yan, Y.; Wang, H.; Song, Y.; Zhu, D.; Shen, Y.; Li, Y., Combinatorial biosynthesis of oxidized combamides using cytochrome P450 enzymes from different polycyclic tetramate macrolactam pathways. ACS Synth. Biol. 2021, 10, 2434–2439.

24. Li, X.; Wang, H.; Shen, Y.; Li, Y.; Du, L., OX4 is an NADPH-dependent dehydrogenase catalyzing an extended Michael addition reaction to form the six-membered ring in the antifungal HSAF. Biochemistry 2019, 58, 5245–5248.

25. Cao, S.; Blodgett, J. A.; Clardy, J., Targeted discovery of polycyclic tetramate macrolactams from an environmental Streptomyces strain. Org. Lett. 2010, 12, 4652–4654.

26. Ding, W.; Tu, J.; Zhang, H.; Wei, X.; Ju, J.; Li, Q., Genome mining and metabolic profiling uncover polycyclic tetramate macrolactams from Streptomyces koyangensis SCSIO 5802. Mar. Drugs 2021, 19, 440.

27. Jiao, Y.-J.; Liu, Y.; Wang, H.-X.; Zhu, D.-Y.; Shen, Y.-M.; Li, Y.-Y., Expression of the clifednamide biosynthetic pathway in Streptomyces generates 27, 28-seco-derivatives. J. Nat. Prod. 2020, 83, 2803–2808.

28. Hou, L.; Liu, Z.; Yu, D.; Li, H.; Ju, J.; Li, W., Targeted isolation of new polycyclic tetramate macrolactams from the deepsea-derived Streptomyces somaliensis SCSIO ZH66. Bioorg. Chem. 2020, 101, 103954.

29. Liu, Y.; Wang, H.; Song, R.; Chen, J.; Li, T.; Li, Y.; Du, L.; Shen, Y., Targeted discovery and combinatorial biosynthesis of polycyclic tetramate macrolactam combamides A–E. Org. Lett. 2018, 20, 3504–3508.

30. Luo, Y.; Huang, H.; Liang, J.; Wang, M.; Lu, L.; Shao, Z.; Cobb, R. E.; Zhao, H., Activation and characterization of a cryptic polycyclic tetramate macrolactam biosynthetic gene cluster. Nat. Commun. 2013, 4, 2894–2901.

31. Olano, C.; García, I.; González, A.; Rodriguez, M.; Rozas, D.; Rubio, J.; Sánchez-Hidalgo, M.; Braña, A. F.; Méndez, C.; Salas, J. A., Activation and identification of five clusters for secondary metabolites in Streptomyces albus J1074. Microb. Biotechnol. 2014, 7, 242–256.

32. Blin, K.; Shaw, S.; Augustijn, H. E.; Reitz, Z. L.; Biermann, F.; Alanjary, M.; Fetter, A.; Terlouw, B. R.; Metcalf, W. W.; Helfrich, E. J., antiSMASH 7.0: new and improved predictions for detection, regulation, chemical structures and visualisation. Nucleic Acids Res. 2023, 51, W46–W50.

33. D’Agostino, P. M.; Gulder, T. A. M., Direct pathway cloning combined with sequence- and ligation-independent cloning for fast biosynthetic gene cluster refactoring and heterologous expression. ACS Synth. Biol. 2018, 7, 1702–1708.

34. Greunke, C.; Duell, E. R.; D’Agostino, P. M.; Glöckle, A.; Lamm, K.; Gulder, T. A. M., Direct pathway cloning (DiPaC) to unlock natural product biosynthetic potential. Metab. Eng. 2018, 47, 334–345.

35. Li, M. Z.; Elledge, S. J., Harnessing homologous recombination in vitro to generate recombinant DNA via SLIC. Nat. Methods 2007, 4, 251–256.

36. Bertasso, M.; Holzenkaempfer, M.; Zeeck, A.; Stackebrandt, E.; Beil, W.; Fiedler, H.-P., Ripromycin and other polycyclic macrolactams from Streptomyces sp. Tü 6239. J. Antibiot. 2003, 56, 364–371.

37. Bibb, M. J.; White, J.; Ward, J. M.; Janssen, G. R., The mRNA for the 23S rRNA methylase encoded by the ermE gene of Saccharopolyspora erythraea is translated in the absence of a conventional ribosome-binding site. Mol. Microbiol. 1994, 14, 533–545.

38. Gibson, D. G.; Young, L.; Chuang, R.-Y.; Venter, J. C.; Hutchison III, C. A.; Smith, H. O., Enzymatic assembly of DNA molecules up to several hundred kilobases. Nat. Methods 2009, 6, 343–345.

39. Myronovskyi, M.; Luzhetskyy, A., Native and engineered promoters in natural product discovery. Nat. Prod. Rep. 2016, 33, 1006–1019.

40. Shao, Z.; Rao, G.; Li, C.; Abil, Z.; Luo, Y.; Zhao, H., Refactoring the silent spectinabilin gene cluster using a plug-and-play scaffold. ACS Synth. Biol. 2013, 2, 662–669.

41. Wang, W.; Li, X.; Wang, J.; Xiang, S.; Feng, X.; Yang, K., An engineered strong promoter for Streptomycetes. Appl. Environ. Microbiol. 2013, 79, 4484–4492.

42. Labes, G.; Bibb, M.; Wohlleben, W., Isolation and characterization of a strong promoter element from the Streptomyces ghanaensis phage I19 using the gentamicin resistance gene (aacC1) of Tn1696 as reporter. Microbiol. 1997, 143, 1503–1512.

43. Luo, Y.; Zhang, L.; Barton, K. W.; Zhao, H., Systematic identification of a panel of strong constitutive promoters from Streptomyces albus. ACS Synth. Biol. 2015, 4, 1001–1010.

44. Yu, H.-L.; Jiang, S.-H.; Bu, X.-L.; Wang, J.-H.; Weng, J.-Y.; Yang, X.-M.; He, K.-Y.; Zhang, Z.-G.; Ao, P.; Xu, J., Structural diversity of anti-pancreatic cancer capsimycins identified in mangrove-derived Streptomyces xiamenensis 318 and post-modification via a novel cytochrome P450 monooxygenase. Sci. Rep. 2017, 7, 1–14.

45. Qi, Y.; Ding, E.; Blodgett, J. A., Native and engineered clifednamide biosynthesis in multiple Streptomyces spp. ACS Synth. Biol. 2017, 7, 357–362.

46. Yang, J.; Qi, Y.; Blodgett, J. A.; Wencewicz, T. A., Multifunctional P450 monooxygenase CftA diversifies the clifednamide pool through tandem C–H bond activations. J. Nat. Prod. 2022, 47–55.

