## Supplementary material for "A Plug-and-Play System for Polycyclic Tetramate Macrolactam Expression and Functionalization": ESI File

**Contents**

1. Analysis of the basic plug-and-play system ... 2
2. Cloning strategy to establish the plug-and-play system ... 3
3. Analysis of capsimycin B (7) production by IkaD ... 4
4. Structure elucidation of compound 9 ... 5
5. HR-MS analysis of the produced PoTeMs ... 6
6. NMR analysis of the produced PoTeMs ... 7
   Ikarugamycin (1) ... 7
   Capsimycin G (8) ... 11
   Butremycin (9) ... 14
   Clifednamide A (10) ... 17
   Clifednamide C (11) ... 21
7. Bacterial strains, primer, plasmids, and sequences ... 25
   Strains ... 25
   Primers ... 25
   Plasmids ... 26
   Gene sequences ... 28
   Sequences of the used promoters ... 32
8. Cloning ... 34
9. Literature ... 50

#### 1. Analysis of the basic plug-and-play system

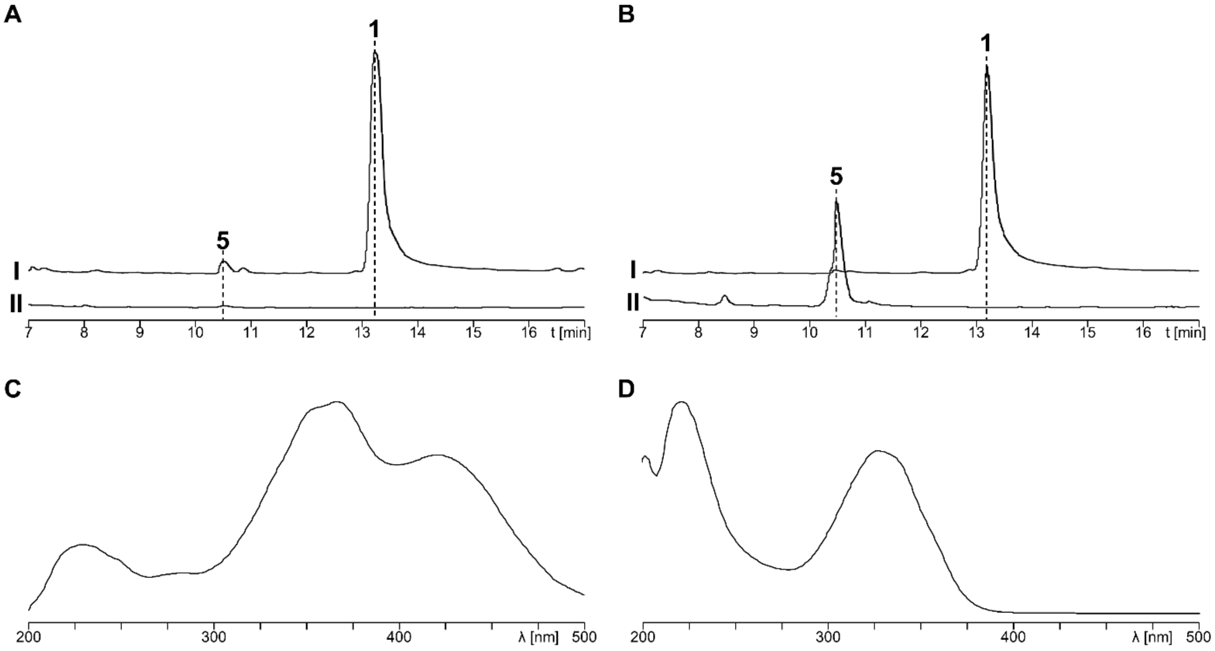


**Figure S1.** Analysis of the PoTeMs produced by the basic plug-and-play constructs. The plasmids pSET152-ermE*::ikaABC* gapdhP(EL) (**I**) and pSET152-ermE*::ikaA*-gapdhP(EL) (**II**) were expressed in *S. albus* DSM 40313. The extracts of **A.** the medium and **B.** the cells were analyzed by HPLC-UV (**I**: 220 nm and **II**: 365 nm). The UV spectra of **C.** lysobacterene A (**5**) and **D.** ikarugamycin (**1**) are shown.

#### 2. Cloning strategy to establish the plug-and-play system

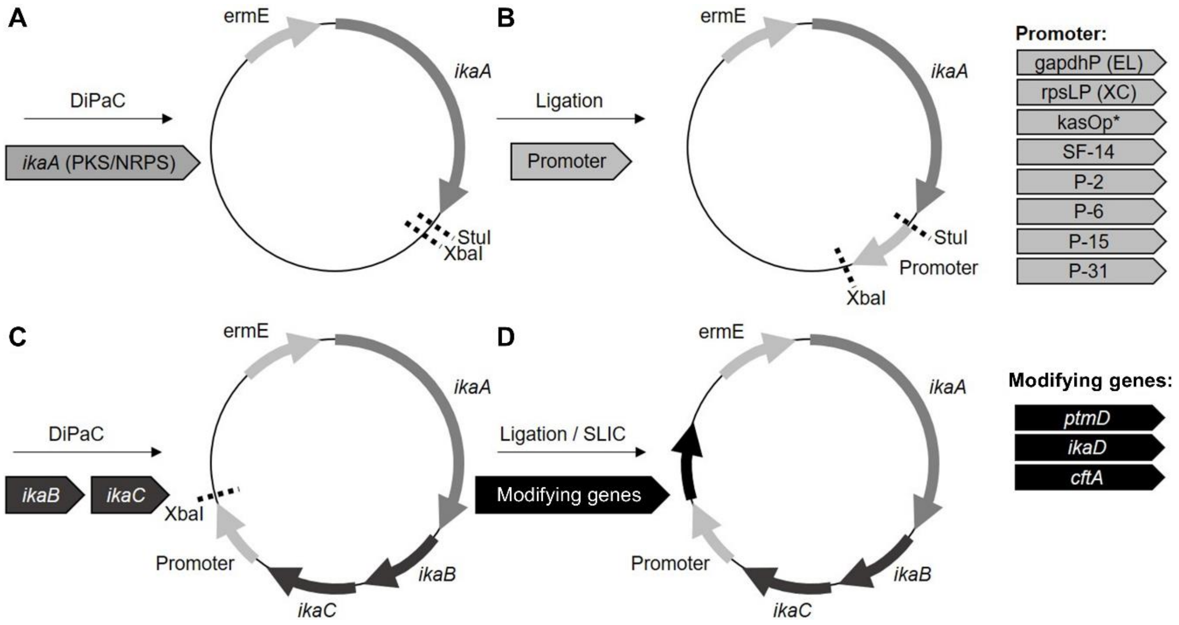


**Figure S2.** Cloning strategy to establish the PoTeM plug-and-play system. **A.** The iPKS/NRPS gene *ikaA* was introduced into pSET152-ermE by DiPaC. **B.** Eight different promoters were added to build the basic eight expression vectors for this plug-and-play system by ligation cloning. **C.** The two additional genes from the biosynthetic gene cluster encoding **1** (*ikaBC*) were added by DiPaC. **D.** The expression system was finalized by adding three different modifying genes (*ptmD*, *ikaD*, and *cftA*) by ligation cloning.

#### 3. Analysis of capsimycin B (7) production by IkaD

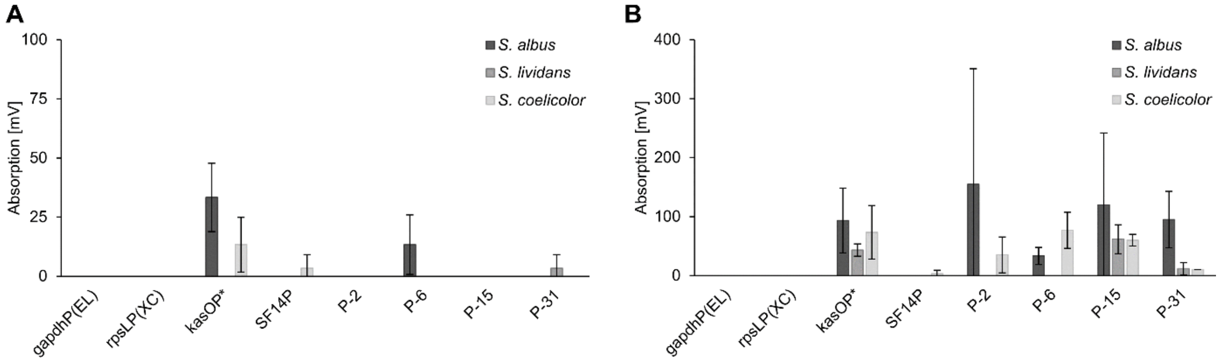


**Figure S3.** Conversion of **1** to **7** by IkaD using different promoters and *Streptomyces* expression hosts. Amount of **7** in **A.** the medium and **B.** the cells, with tailoring gene expression under control of eight different promoters in three different stains: *S. albus* DSM 40313 (black), *S. lividans* TK24 (dark grey), and *S. coelicolor* M1154 (light grey).

#### 4. Structure elucidation of compound 9

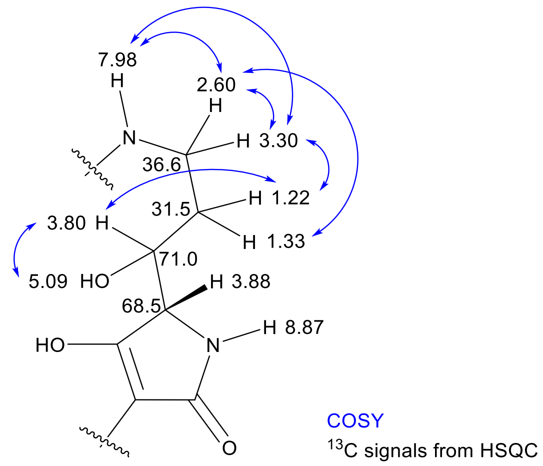


**Figure S4.** Structure elucidation of compound **9**. The section depicts the region with divergent NMR signals compared to **1**. Compound **9** was analyzed by 1D and 2D NMR and the position of the hydroxy group was localized at C-3. Crucial COSY interactions are indicated by blue double-arrows.

#### 5. HR-MS analysis of the produced PoTeMs

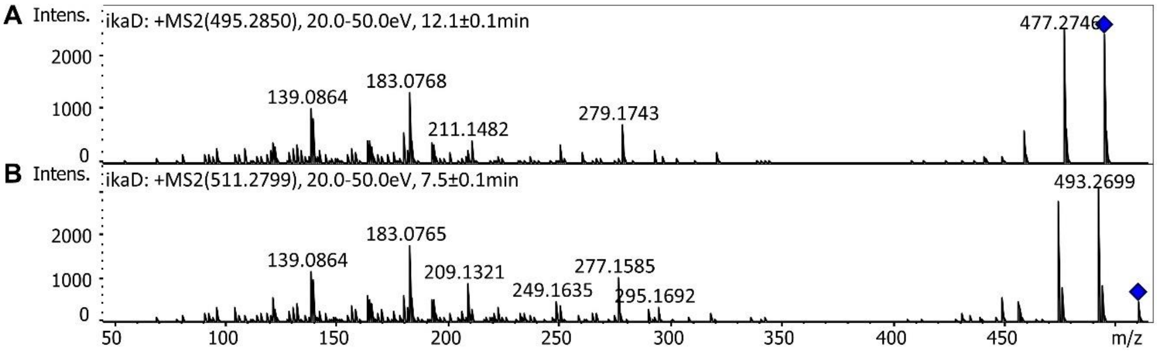


**Figure S5.** MS/MS analysis of the PoTeM modified by IkaD. **A.** An *m/z* 495.2850 was observed fitting **7** ($[M+H]^+$ calculated *m/z* 495.2853). **B.** An *m/z* 511.2799 was observed fitting **8** ($[M+H]^+$ calculated *m/z* 511.2803).

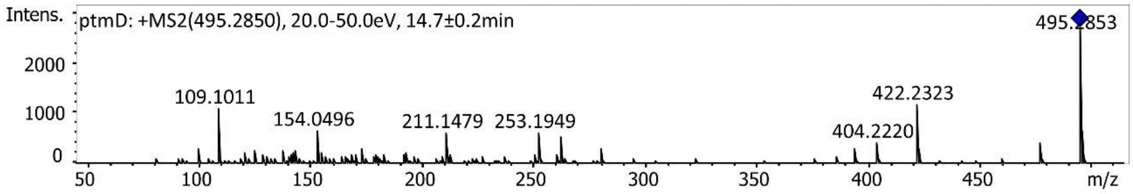


**Figure S6.** MS/MS analysis of the PoTeM modified by PtmD. An *m/z* 495.2850 was observed fitting **9** ($[M+H]^+$ calculated *m/z* 495.2853).

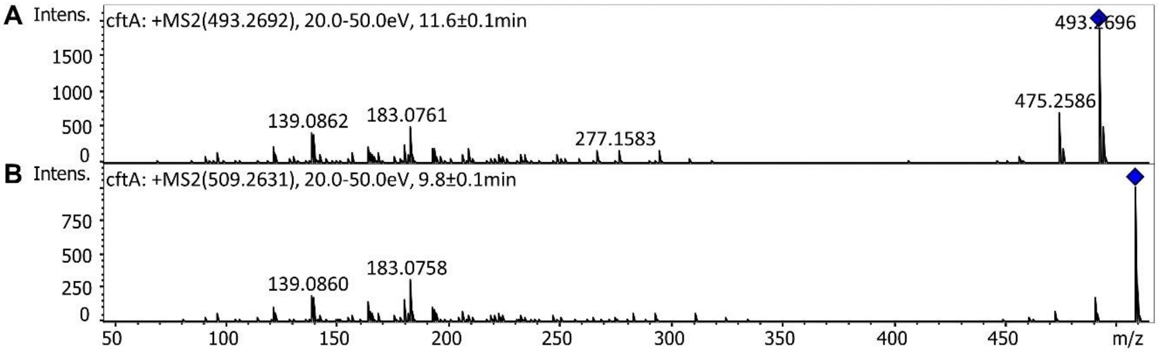


**Figure S7.** MS/MS analysis of the PoTeM modified by CftA. **A.** An *m/z* 493.2692 was observed fitting **10** ($[M+H]^+$ calculated *m/z* 493.2697). **B.** An *m/z* 509.2631 was observed fitting **11** ($[M+H]^+$ calculated *m/z* 509.2646).

#### 6. NMR analysis of the produced PoTeMs

##### Ikarugamycin (1)

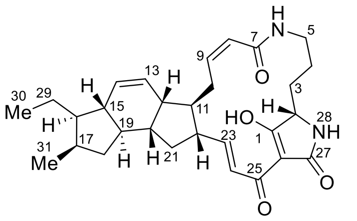


**Table S1.** [1]H, [13]C NMR (600 MHz, DMSO-$d_6$) of ikarugamycin (**1**).

| | $\delta_C$ | $\delta_H$ (m, *J*) | COSY |
|---|---|---|---|
| 1 | 195.9 | / | |
| 2 | 61.1 | 3.83 (d, 5.9 Hz) | 1, 3 |
| 3 | 26.8 | 1.69–1.75 (m), 1.81–1.88 (m) | 2, 3, 4 |
| 4 | 20.4 | 1.03–1.10 (m), 1.35–1.40 (m) | 3, 4, 5 |
| 5 | 38.2 | 2.42–2.48 (m), 3.20–3.25 (m) | 4, 5, 6 |
| 6 | / | 7.87 (t, 5.6 Hz) | 5 |
| 7 | 165.5 | / | |
| 8 | 124.3 | 5.76 (d, 12.7 Hz) | 9, 10 |
| 9 | 139.3 | 5.98 (td, 11.1, 3.8 Hz) | 8, 10 |
| 10 | 24.8 | 2.19–2.28 (m), 3.51–3.56 (m) | 8, 9, 10, 11 |
| 11 | 48.2 | 1.45–1.55 (m) | 10, 11, 22 |
| 12 | 42.6 | 2.49–2.53 (m) | 12, 13, 14, 20 |
| 13 | 128.8 | 5.73 (dt, 9.9, 2.8 Hz) | 12, 14 |
| 14 | 130.6 | 5.91 (dt, 2.1 Hz) | 12, 13, 15 |
| 15 | 46.7 | 1.50–1.56 (m) | 13, 14, 16, 19 |
| 16 | 46.5 | 1.28–1.34 (m) | 15, 17 |
| 17 | 32.5 | 2.19–2.28 (m) | 16, 18, 31 |
| 18 | 38.1 | 0.66 (td, 12.0, 6.9 Hz),<br>2.09 (dt, 12.8, 7.8 Hz) | 17, 18, 19 |
| 19 | 48.3 | 1.11–1.20 (m) | 15, 18, 20 |
| 20 | 41.2 | 2.00–2.06 (m) | 12, 19 |
| 21 | 36.1 | 1.19–1.28 (m), 2.00–2.06 (m) | 21, 22 |
| 22 | 49.6 | 2.32–2.41 (m) | 11, 21, 23 |
| 23 | 150.3 | 6.64 (dd, 15.5, 10.2 Hz) | 22, 24 |
| 24 | 122.0 | 6.98 (d, 15.5 Hz) | 23 |
| 25 | 171.4 | / | |
| 26 | 100.8 | / | |
| 27 | 175.2 | / | |
| 28 | / | 8.69 (s) | |
| 29 | 21.1 | 1.28–1.34 (m), 1.42–1.50 (m) | 30 |
| 30 | 13.2 | 0.91 (t, 7.2 Hz) | 29 |
| 31 | 17.7 | 0.86 (d, 7.1 Hz) | 17 |
| 32 | / | / | |

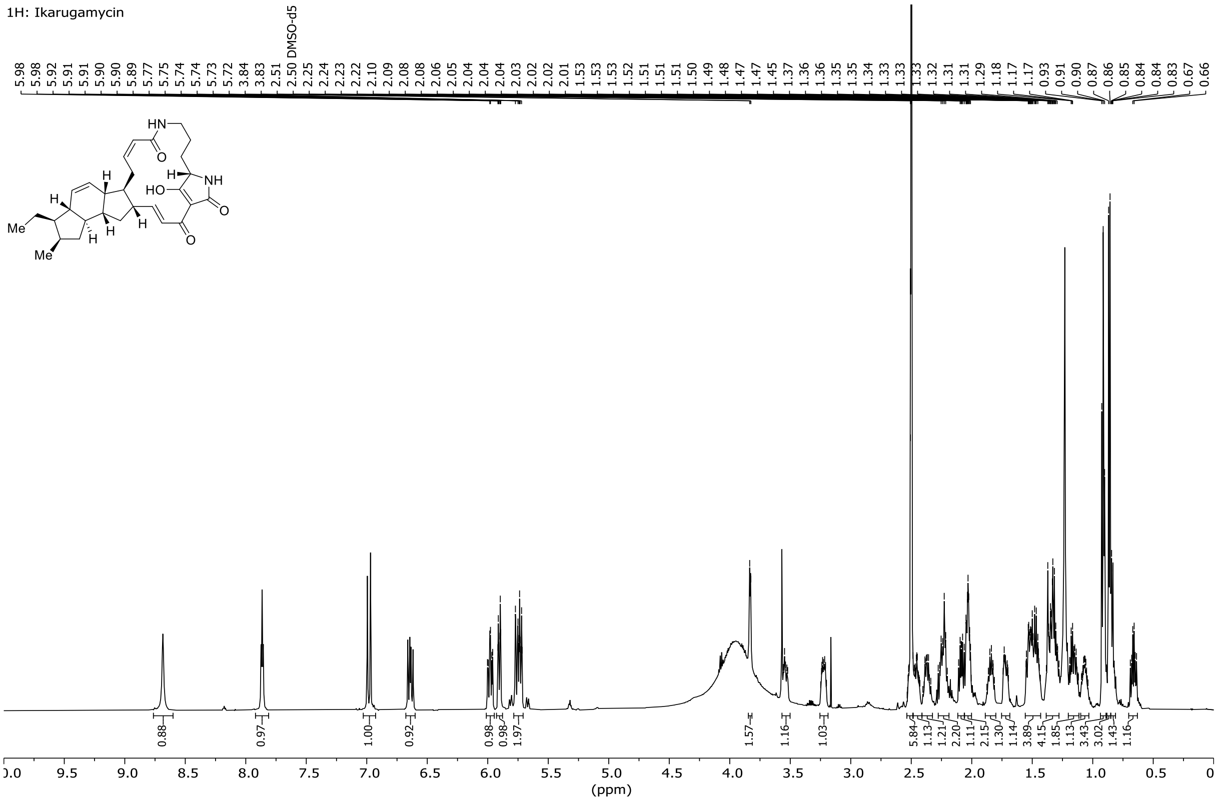


**Figure S8.** $^{1}$H NMR of ikarugamycin (**1**).

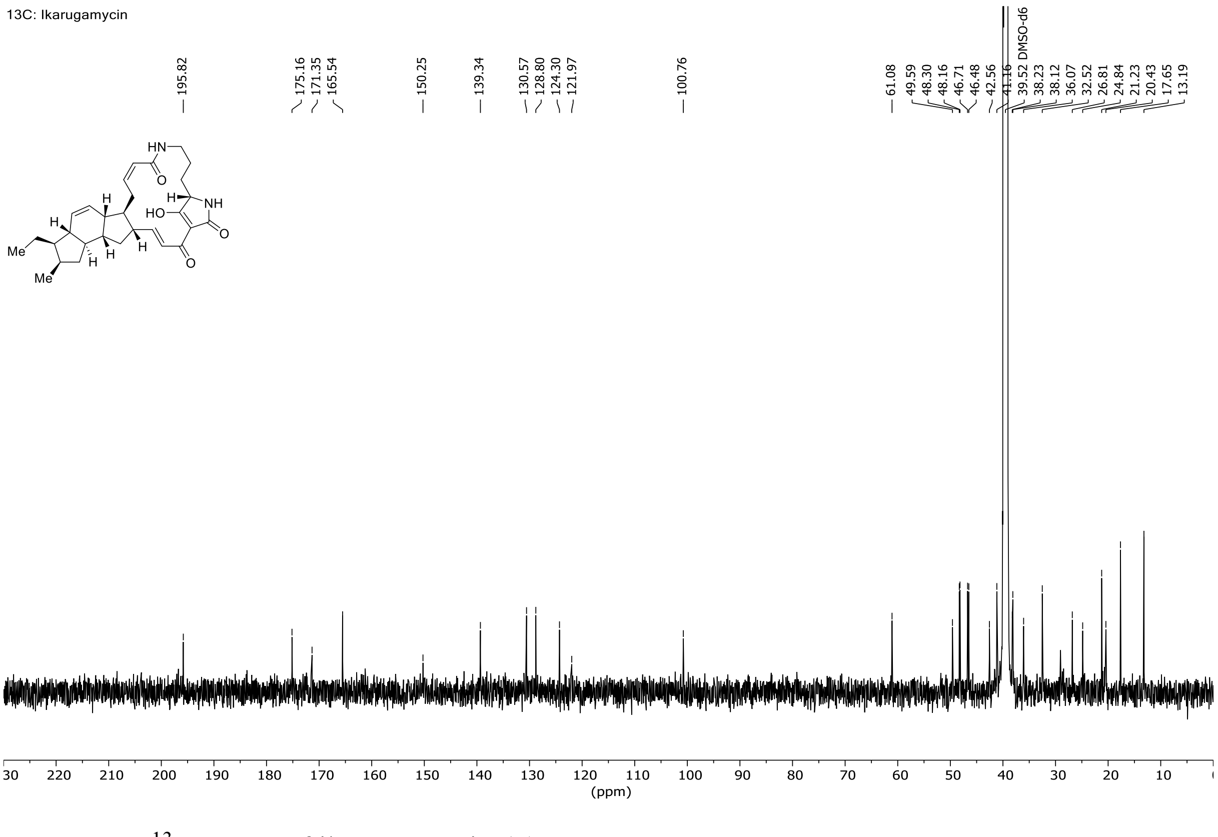


**Figure S9.** $^{13}$C NMR of ikarugamycin (**1**).

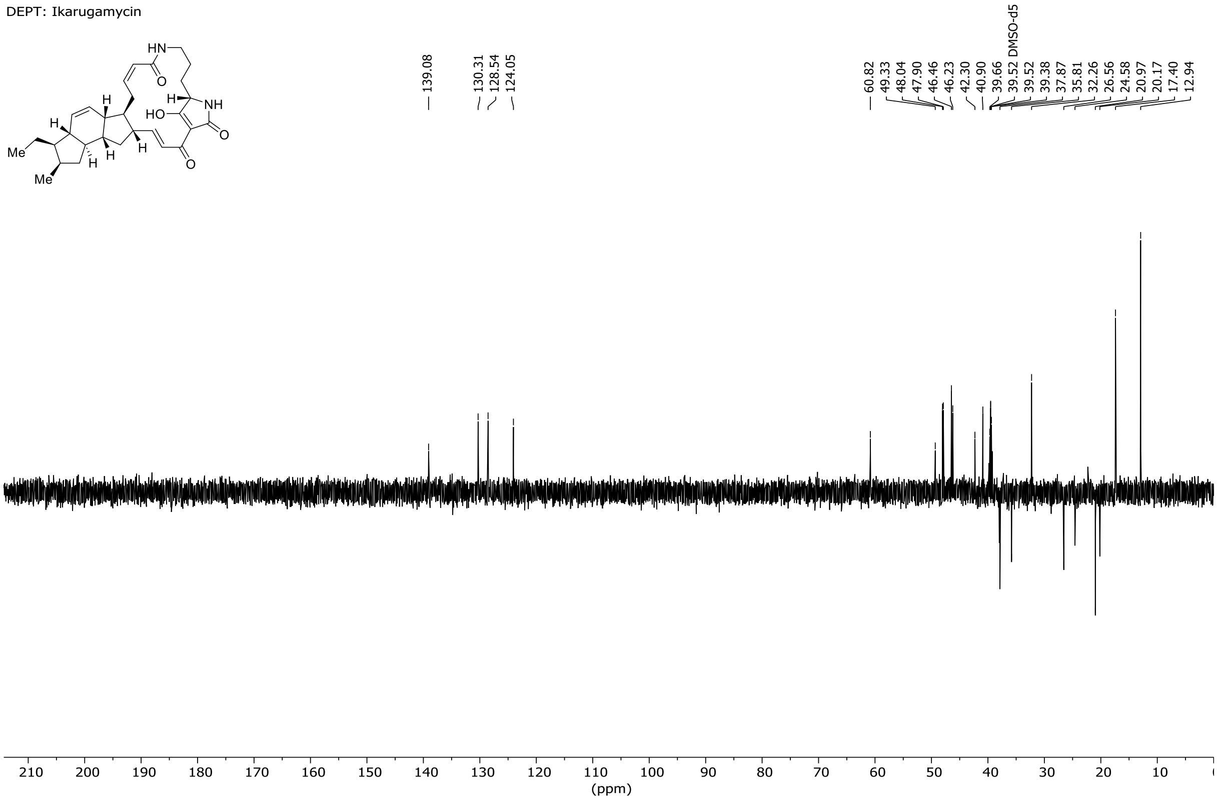


**Figure S10.** DEPT NMR of ikarugamycin (**1**).

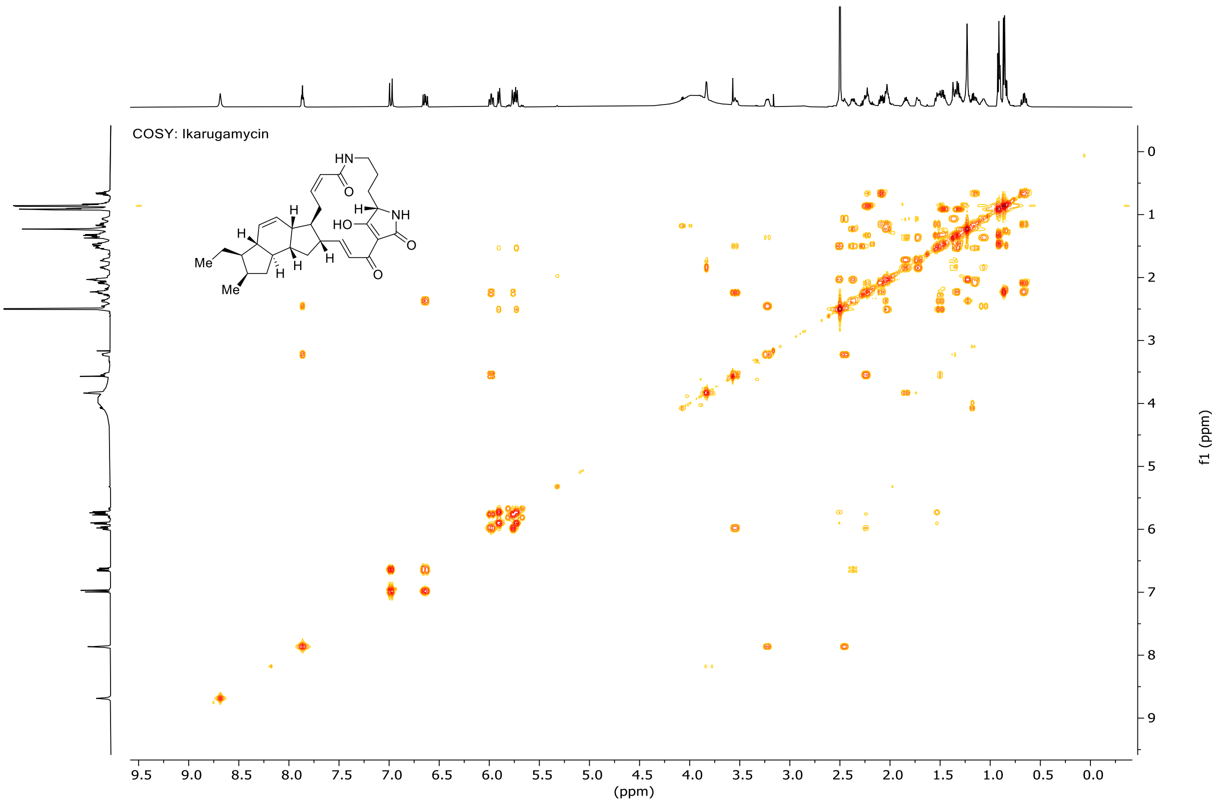


**Figure S11.** COSY NMR of ikarugamycin (**1**).

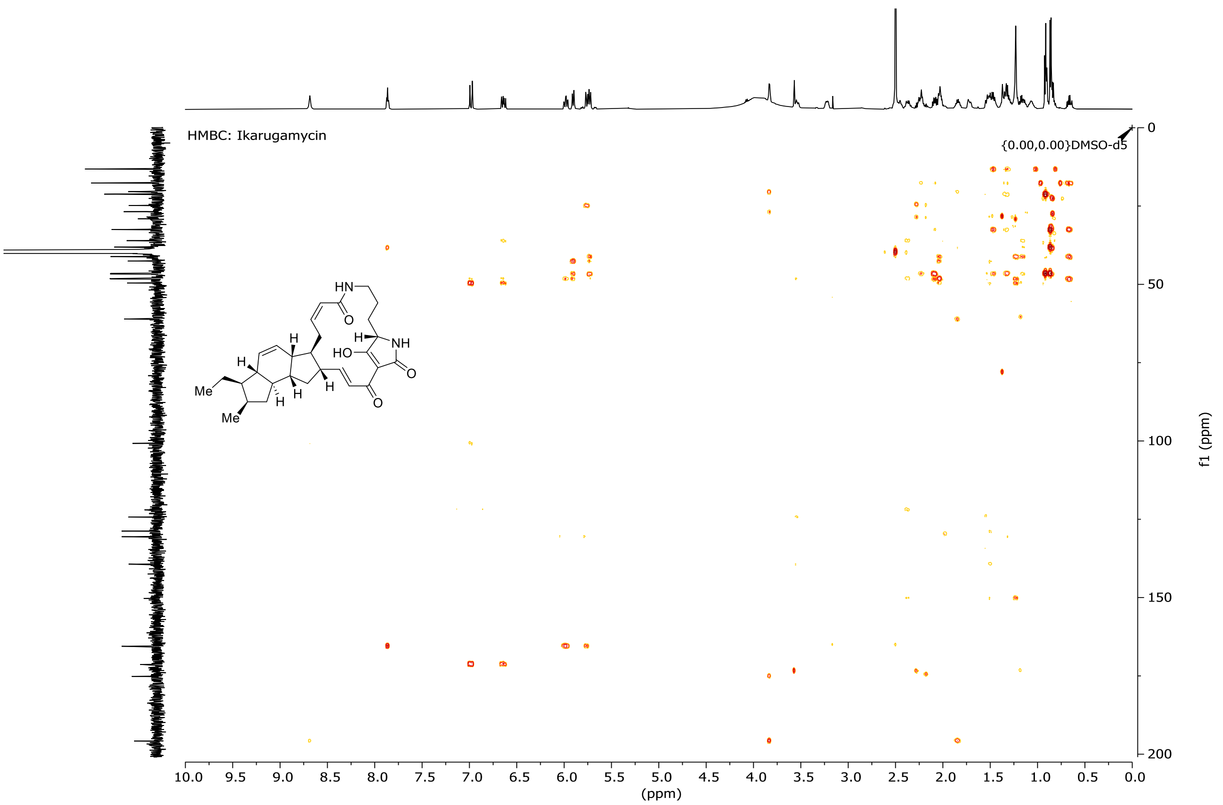


**Figure S12.** HMBC NMR of ikarugamycin (**1**).

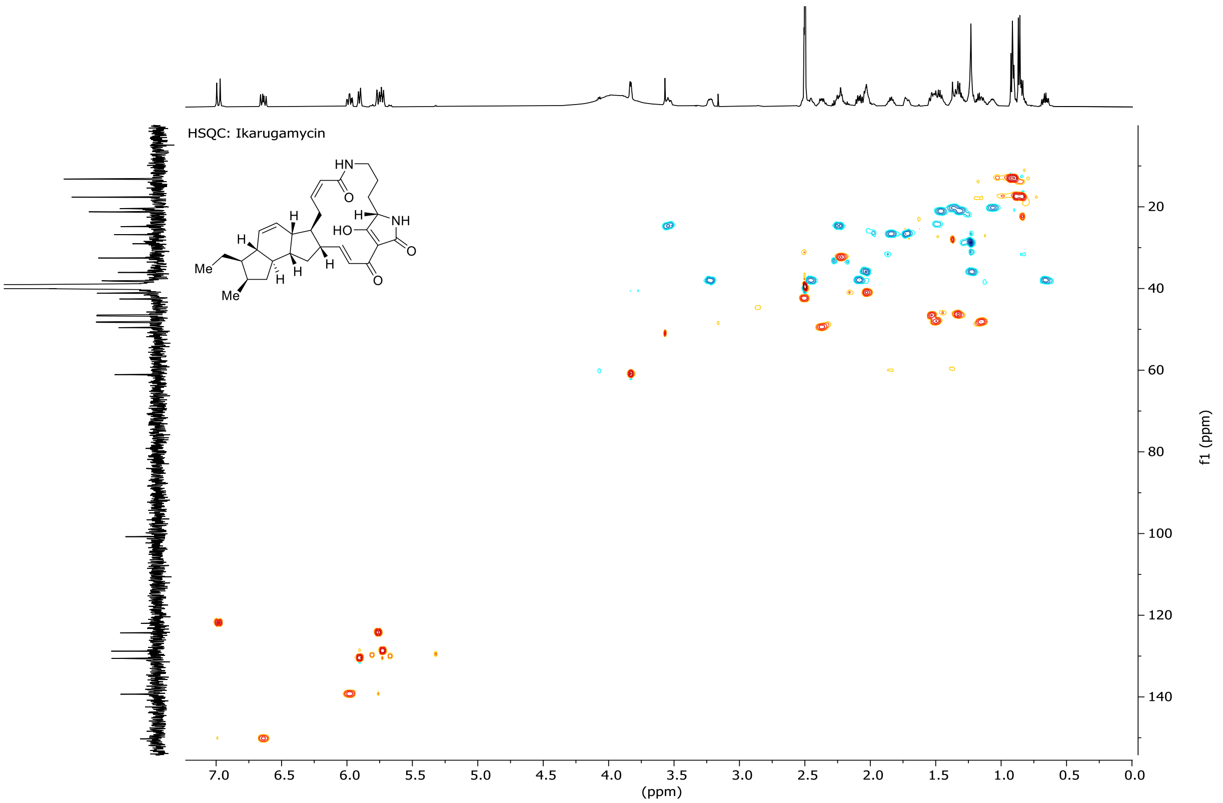


**Figure S13.** HSQC NMR of ikarugamycin (**1**).

**Capsimycin G (8)**

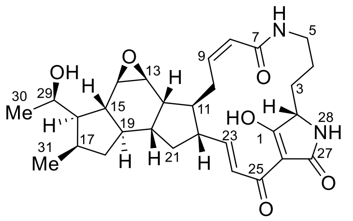


**Table S2.** $^{1}$H, $^{13}$C NMR (600 MHz, DMSO-$d_6$) of capsimycin G (**8**).

| | $\delta_C$ | $\delta_H$ (m, $J$) | COSY |
|---|---|---|---|
| 1 | 195.9 | / | |
| 2 | 61.1 | 3.83 (d, 5.6 Hz) | 3 |
| 3 | 26.8 | 1.85 (m), 1.73 (m) | 2, 4 |
| 4 | 20.4 | 1.37 (m), 1.07 (m) | 3, 5 |
| 5 | 38.2 | 3.24 (m), 2.45 (m) | 4, 6 |
| 6 | / | 7.89 (t, 5.6 Hz) | 5 |
| 7 | 165.5 | / | |
| 8 | 124.4 | 5.79 (d, 11.6 Hz) | 9, 10 |
| 9 | 139.0 | 6.03 (td, 11.0, 3.5 Hz) | 8, 10 |
| 10 | 25.2 | 3.64 (m), 2.31 (m) | 8, 9, 11 |
| 11 | 45.4 | 1.67 (m) | 10, 22 |
| 12 | 40.0 | 2.29 (m) | 11, 20 |
| 13 | 53.0 | 2.90 (d, 4.0 Hz) | 12, 14 |
| 14 | 57.4 | 3.18 (dd, 4.0, 1.7 Hz) | 13, 15 |
| 15 | 46.7 | 0.87 (td, 11.9, 2.0 Hz) | 16, 19 |
| 16 | 51.4 | 1.76 (dt, 11.4, 9.4 Hz) | 15 |
| 17 | 33.2 | 2.26 (m) | 18, 31 |
| 18 | 38.7 | 1.98 (m), 0.58 (m) | 17, 19 |
| 19 | 47.1 | 1.12 (qd, 11.6, 6,6) | 15, 18, 20 |
| 20 | 40.7 | 1.65 (m) | 12, 19 |
| 21 | 36.3 | 1.99 (m), 1.16 (m) | 22 |
| 22 | 48.9 | 2.34 (qd, 10.7, 7.7 Hz) | 11, 21, 23 |
| 23 | 149.8 | 6.64 (dd, 15.5, 10.3 Hz) | 22, 24 |
| 24 | 122.2 | 7.00 (d, 15.6 Hz) | 23 |
| 25 | 171.1 | / | |
| 26 | 100.9 | / | |
| 27 | 175.1 | / | |
| 28 | / | 8.69 (bs) | |
| 29 | 66.5 | 3.70 (dq, 8.8, 6.2 Hz) | |
| 30 | 23.4 | 1.22 (d, 6.2 Hz) | |
| 31 | 18.2 | 1.01 (d, 7.1 Hz) | 17 |

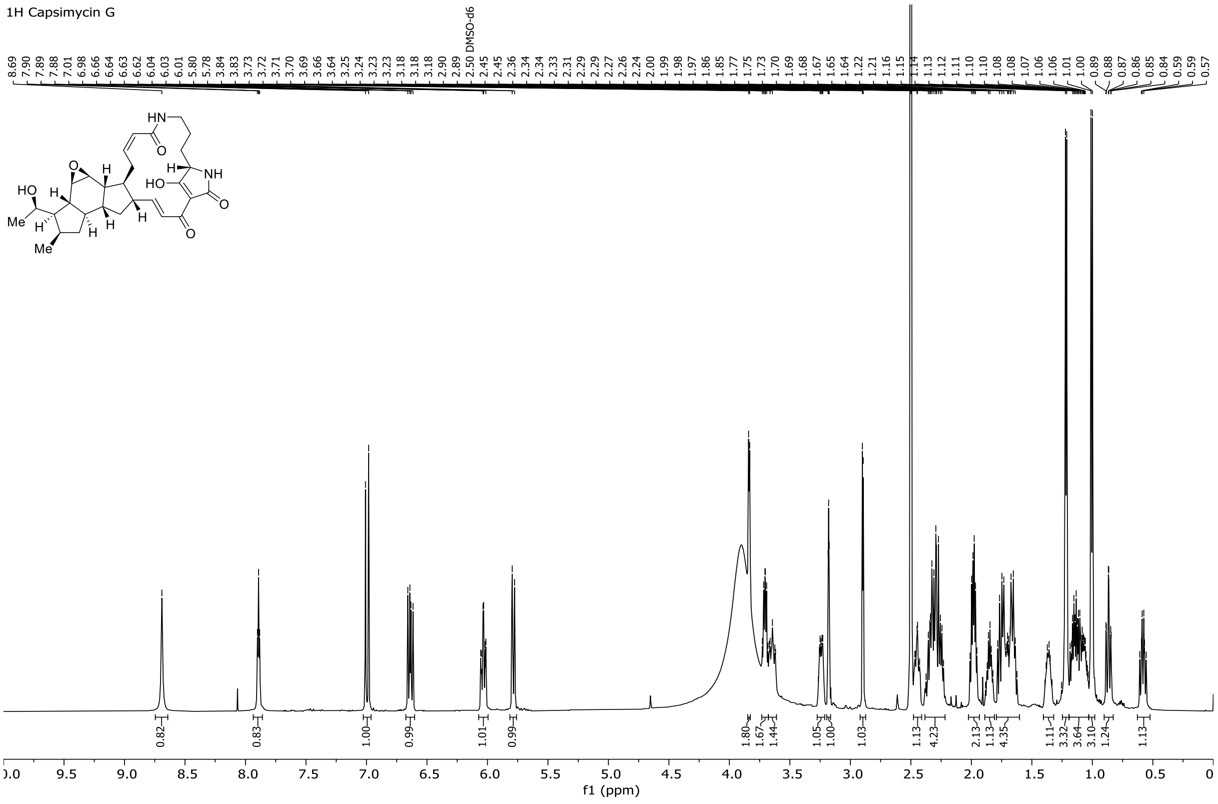


**Figure S14.** $^{1}$H NMR of capsimycin G (**8**)’

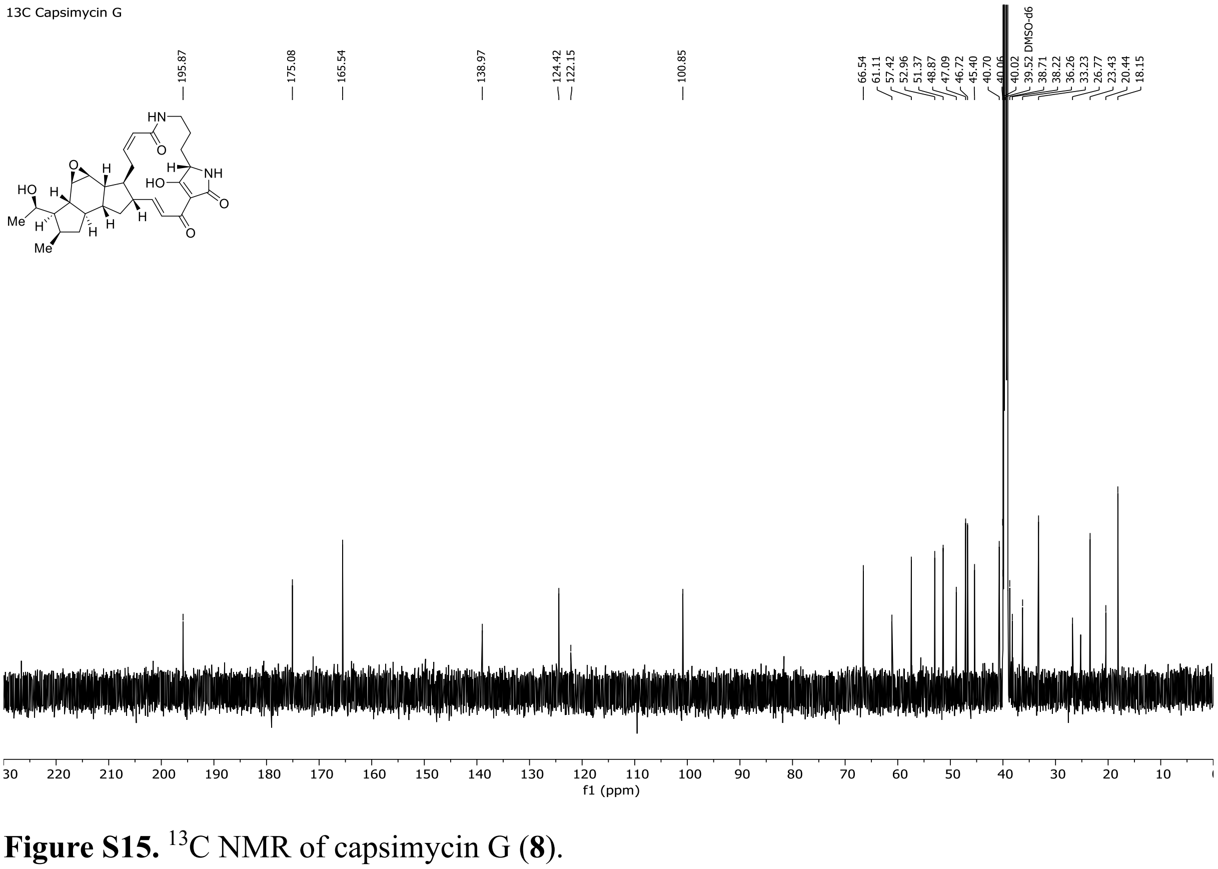


**Figure S15.** $^{13}$C NMR of capsimycin G (**8**).

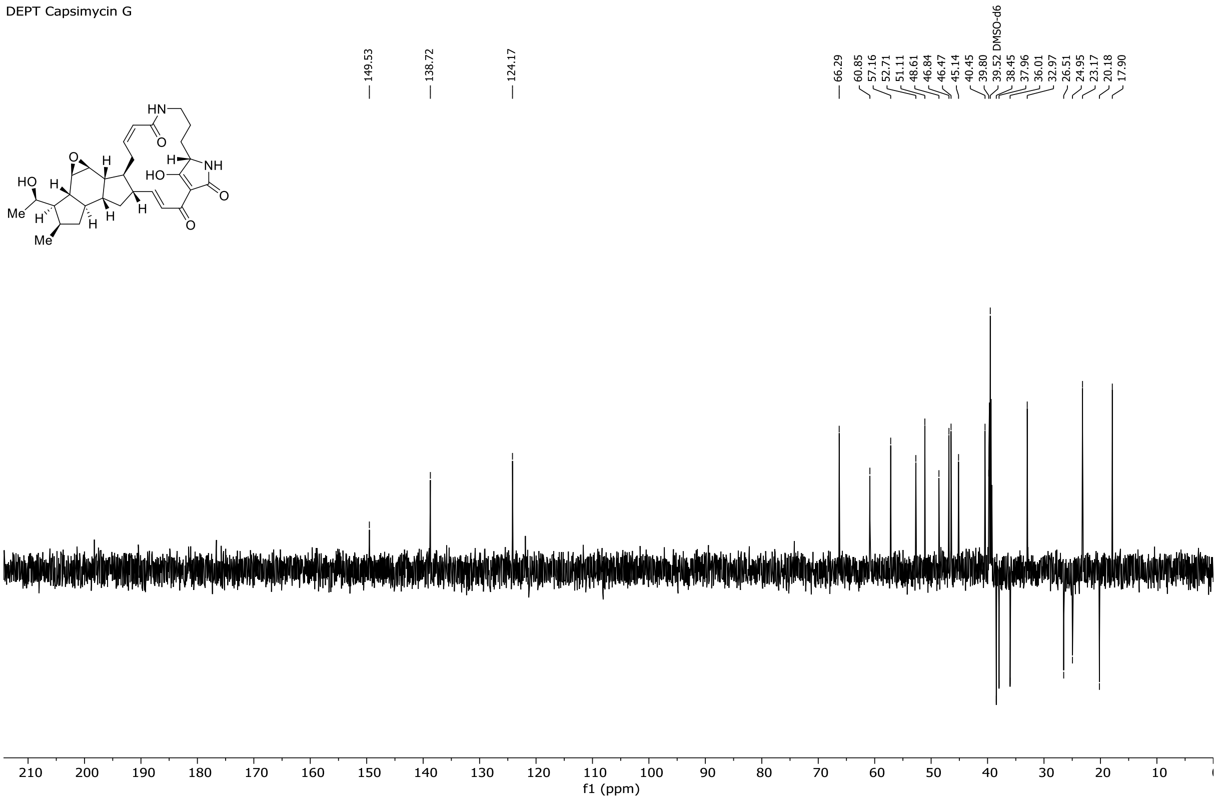


**Figure S16.** DEPT NMR of capsimycin G (**8**).

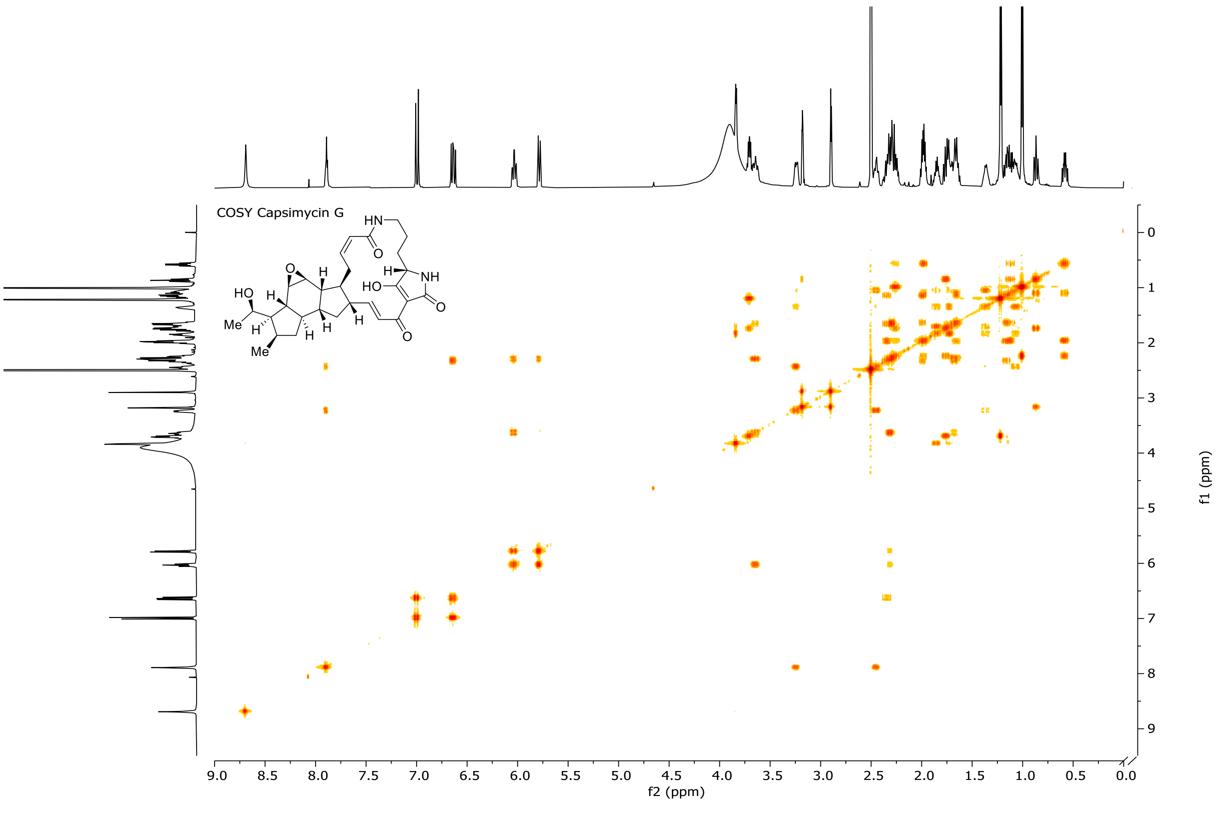


**Figure S17.** COSY NMR of capsimycin G (**8**).

**Butremycin (9)**

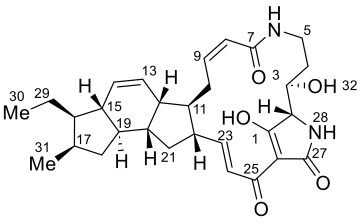


**Table S3.** $^{1}$H, $^{13}$C NMR (500 MHz, DMSO-$d_6$) of butremycin (**9**).

| | $\delta_C$ | $\delta_H$ (m, $J$) | COSY |
|---|---|---|---|
| 1 | n/a | / | |
| 2 | 68.7* | 3.86–3.89 (bs) | |
| 3 | 70.9* | 3.78–3.82 (m) | 4, 32 |
| 4 | 31.0* | 1.28–1.38 (m), 2.21–2.29 (m) | 3, 5 |
| 5 | 36.3* | 2.56–2.64 (m), 3.25–3.34 (m) | 4, 5, 6 |
| 6 | / | 7.98 (t, 5.7 Hz) | 5 |
| 7 | 165.1** | / | |
| 8 | 124.3* | 5.75 (d, 10.3 Hz) | 9, 10 |
| 9 | 139.2* | 6.01 (td, 11.2, 3.0 Hz) | 8, 10 |
| 10 | 24.5* | 2.08–2.17 (m), 3.60–3.70 (m) | 8, 9, 11 |
| 11 | 48.0* | 1.44–1.56 (m) | 10 |
| 12 | 42.1* | 2.50–2.56 (m) | 13, 14, 20 |
| 13 | 128.9* | 5.72 (dt, 9.9, 3.0 Hz) | 12, 14, 15 |
| 14 | 130.5* | 5.90 (d, 9.9 Hz) | 12, 13, 15 |
| 15 | 46.8* | 1.44–1.56 (m) | 13, 14, 16 |
| 16 | 46.5* | 1.28–1.38 (m) | 15, 17 |
| 17 | 32.5* | 2.21–2.26 (m) | 16, 31 |
| 18 | 38.2* | 0.67 (td, 12.0, 6.8 Hz), 2.06–2.16 (m) | 18, 19 |
| 19 | 48.3* | 1.11–1.19 (m) | 18 |
| 20 | 41.2* | 1.97–2.06 (m) | 12 |
| 21 | 36.1* | 1.28–1.38 (m), 1.97–2.06 (m) | 22 |
| 22 | 49.3* | 2.31–2.38 (m) | 21, 23 |
| 23 | 150.4* | 6.60–6.67 (m) | 22, 24 |
| 24 | 122.0* | 6.91 (d, 15.5 Hz) | 23 |
| 25 | n/a | / | |
| 26 | n/a | / | |
| 27 | n/a | / | |
| 28 | / | 8.87 (s) | |
| 29 | 21.1* | 1.28–1.34 (m), 1.44–1.51 (m) | 30 |
| 30 | 13.1* | 0.92 (t, 7.0 Hz) | 29 |
| 31 | 17.7* | 0.86 (d, 7.1 Hz) | 17 |
| 32 | / | 5.09 (bs) | 3 |

* $^{13}$C shift taken from HSQC. ** $^{13}$C shift taken from HMBC.

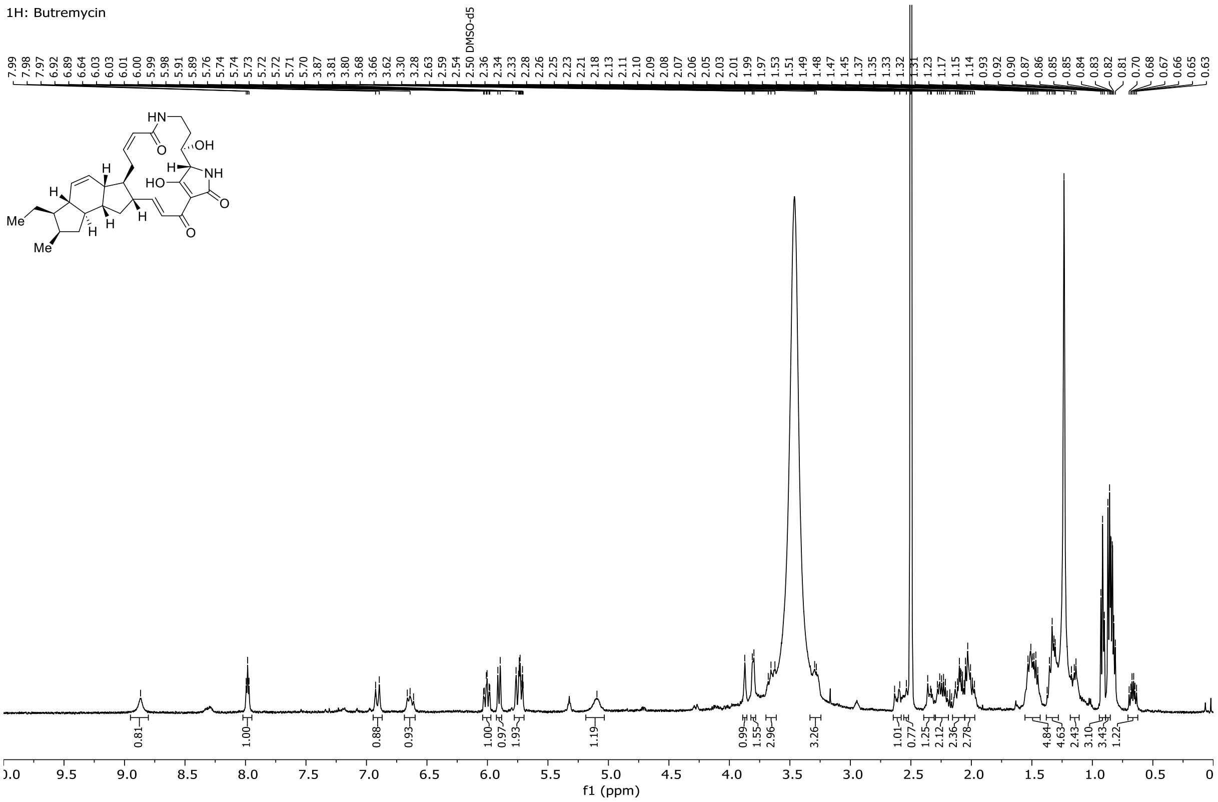


**Figure S18.** $^{1}$H NMR of butremycin (**9**).

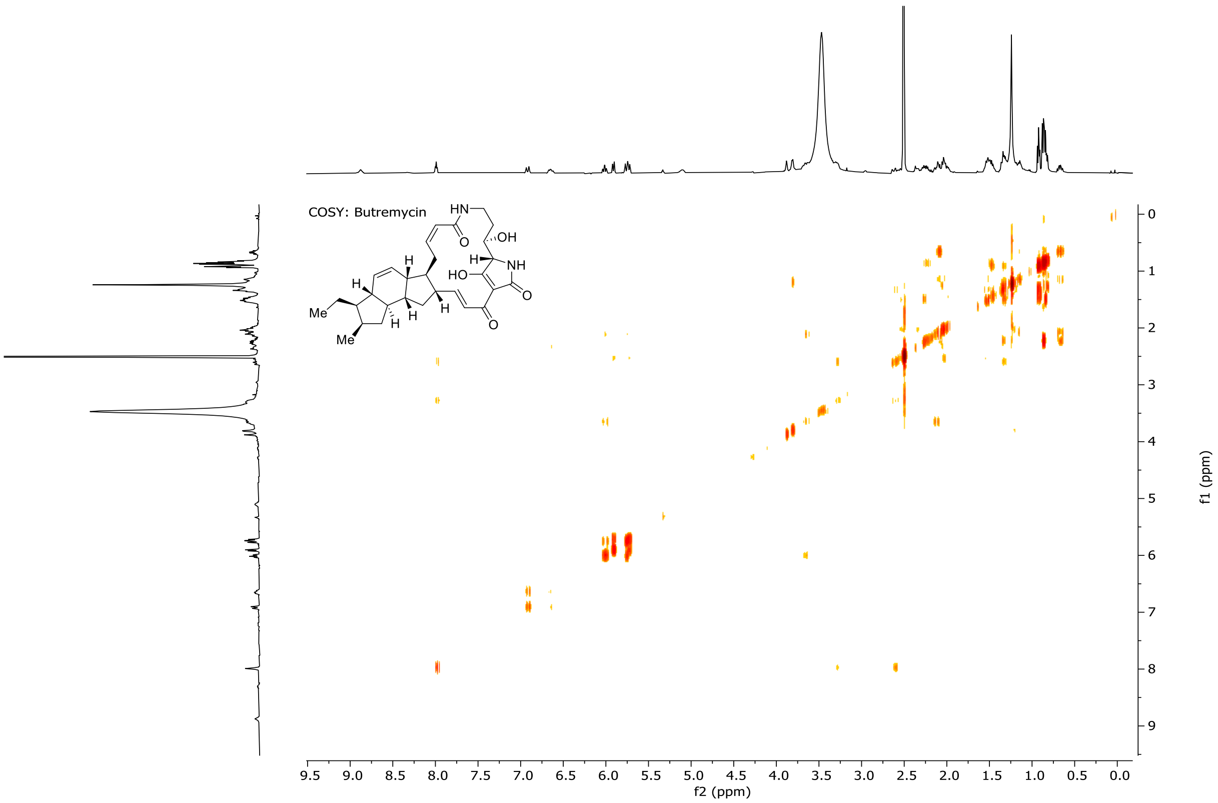


**Figure S19.** COSY NMR of butremycin (**9**).

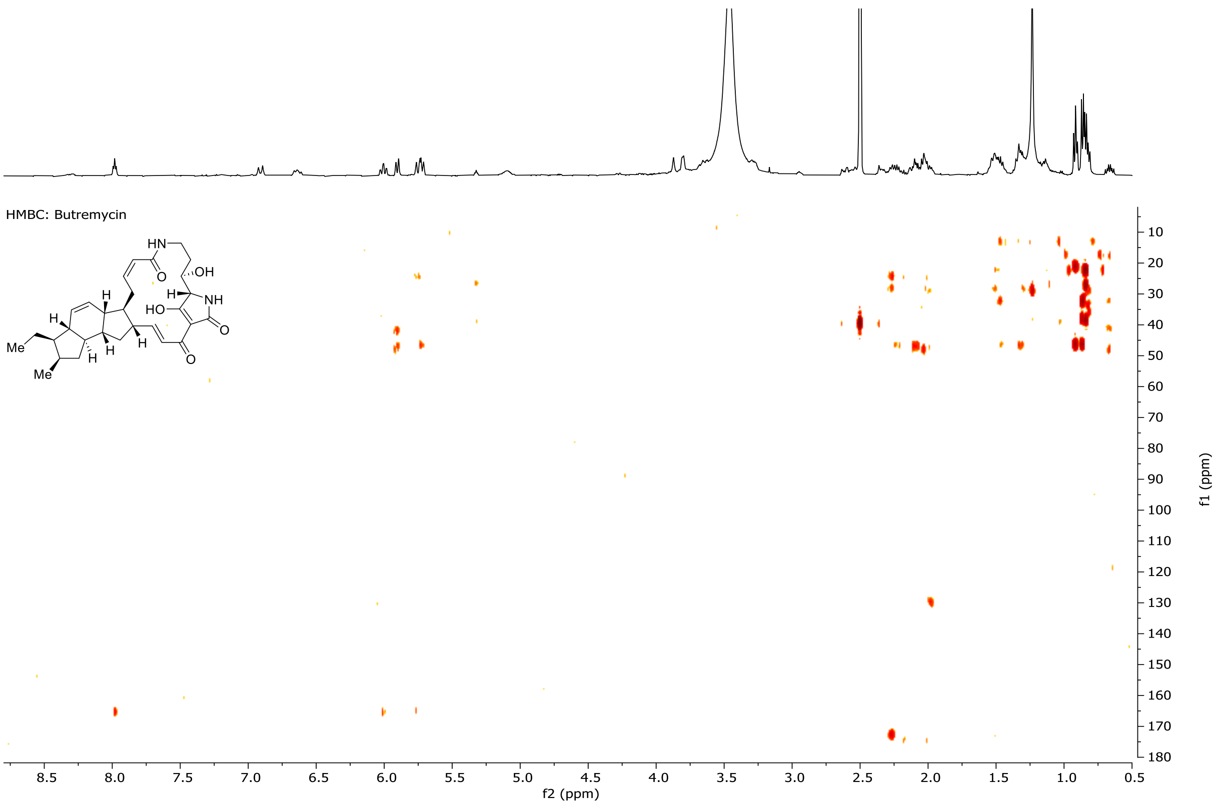


**Figure S20.** HMBC NMR of butremycin (**9**).

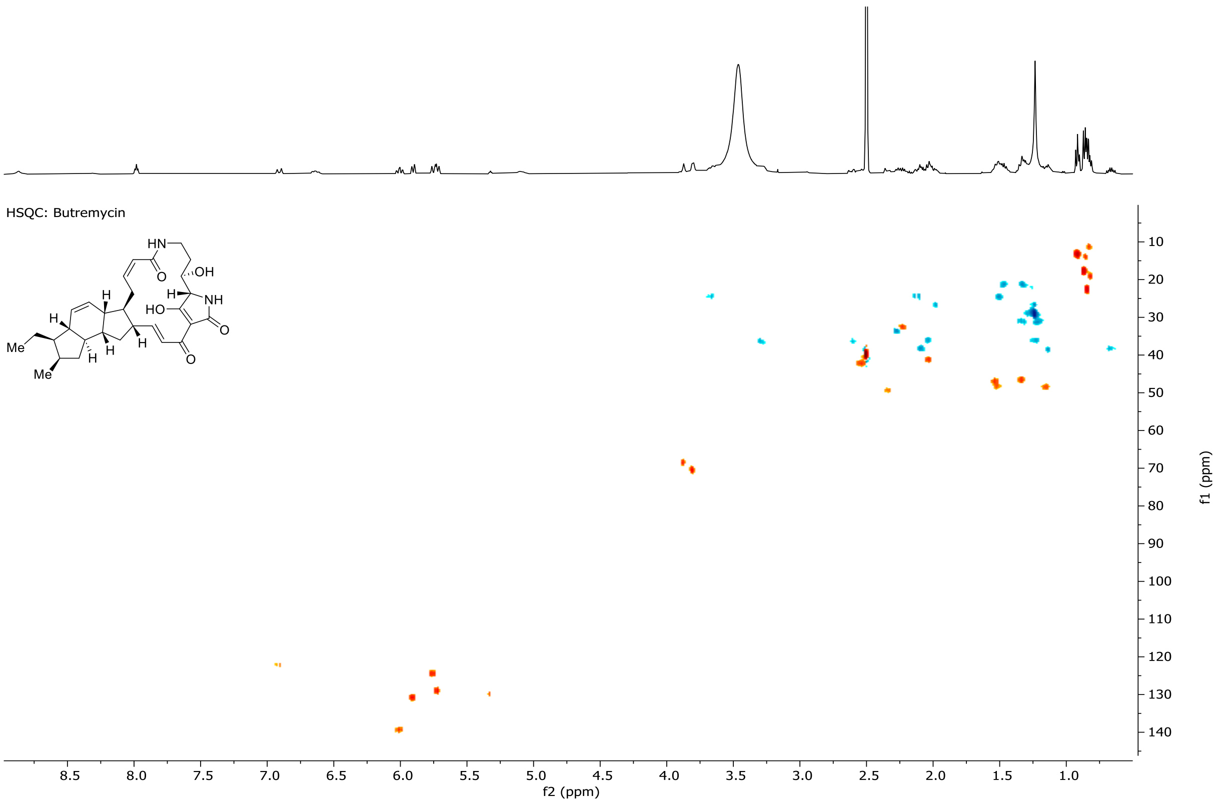


**Figure S21.** HSQC NMR of butremycin (**9**).

**Clifednamide A (10)**

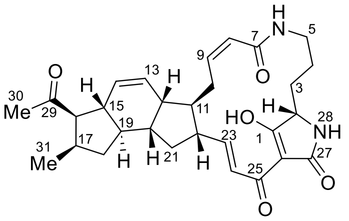


**Table S4.** $^{1}$H, $^{13}$C NMR (600 MHz, Pyridine-$d_5$) of clifednamide A (**10**).

| | **δC** | **δH (m, *J*)** | **COSY** |
|---|---|---|---|
| 1 | 197.2 | / | |
| 2 | 62.4 | 4.10 (d, 5.2 Hz) | 3 |
| 3 | 28.3 | 2.20–2.11 (m) 1H, 2.08–1.94 (m) | 2, 4 |
| 4 | 22.1 | 1.81–1.71 (m) 1H, 1.58–1.49 (m) | 3, 5 |
| 5 | 39.6 | 3.90–3.86 (m) 1H, 2.83–2.75 (m) | 4, 6 |
| 6 | / | 8.85 (bs) | 5 |
| 7 | 167.3 | / | |
| 8 | 125.8 | 6.25 (d, 11.4 Hz) | 9, 10 |
| 9 | 140.1 | 5.96 (td, 11.0, 3.4 Hz) | 8, 10 |
| 10 | 26.0 | 4.14 (ddd, 15.7, 10.3, 4.2 Hz), 2.62–2.51 (m) | 8, 9, 11 |
| 11 | 49.3 | 1.49–1.43 (m) | 10, 12, 22 |
| 12 | 43.2 | 2.47–2.41 (m) | 11, 13, 14, 20 |
| 13 | 129.6 | 5.67 (dt, 9.8, 2.8 Hz) | 12, 14, 15 |
| 14 | 130.3 | 5.77 (d, 9.8 Hz) | 12, 13 |
| 15 | 44.1 | 2.71–2.62 (m) | 13, 16, 19 |
| 16 | 54.4 | 2.95 (dd, 12.1, 10.3 Hz) | 15, 17 |
| 17 | 35.0 | 2.71–2.62 (m) | 16, 18, 31 |
| 18 | 39.7 | 2.08–1.94 (m) 1H, 0.71 (td, 12.0, 7.7 Hz) | 17, 19 |
| 19 | 48.3 | 1.18–1.11 (m) | 15, 18, 20 |
| 20 | 41.7 | 2.08–1.94 (m) | 12, 19, 21 |
| 21 | 37.1 | 2.08–1.94 (m) 1H, 1.11–1.05 (m) | 20, 22 |
| 22 | 50.0 | 2.40–2.31 (m) | 11, 21, 23 |
| 23 | 150.6 | 6.92 (dd, 15.5, 10.2 Hz) | 22, 24 |
| 24 | 124.2 | 7.72–7.80 (m) | 23 |
| 25 | n/a | / | |
| 26 | 102.5 | / | |
| 27 | 167.3 | / | |
| 28 | / | 9.51 (bs) | |
| 29 | 213.0 | / | |
| 30 | 70.7 | 4.65–4.50 (m) | |
| 31 | 19.5 | 0.91 (d, 7.2 Hz) | 17 |

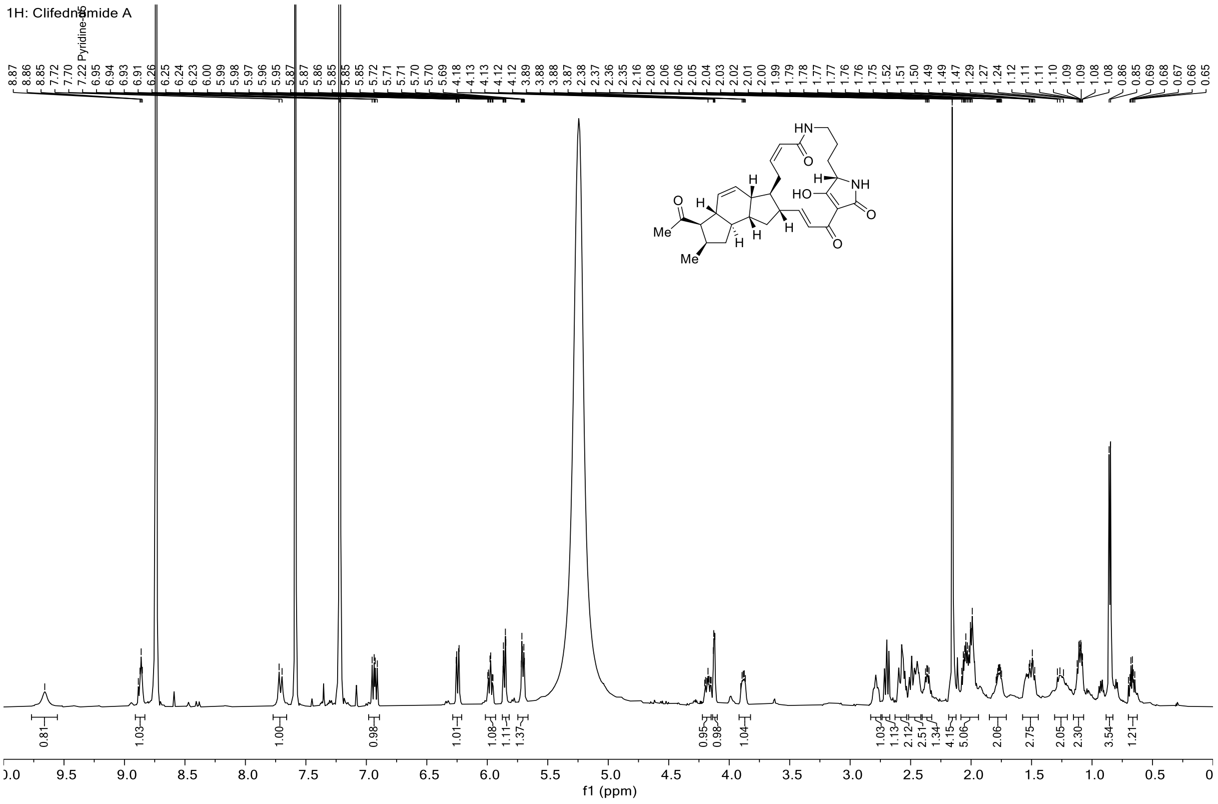

**Figure S22.** $^{1}$H NMR of clifednamide A (**10**).

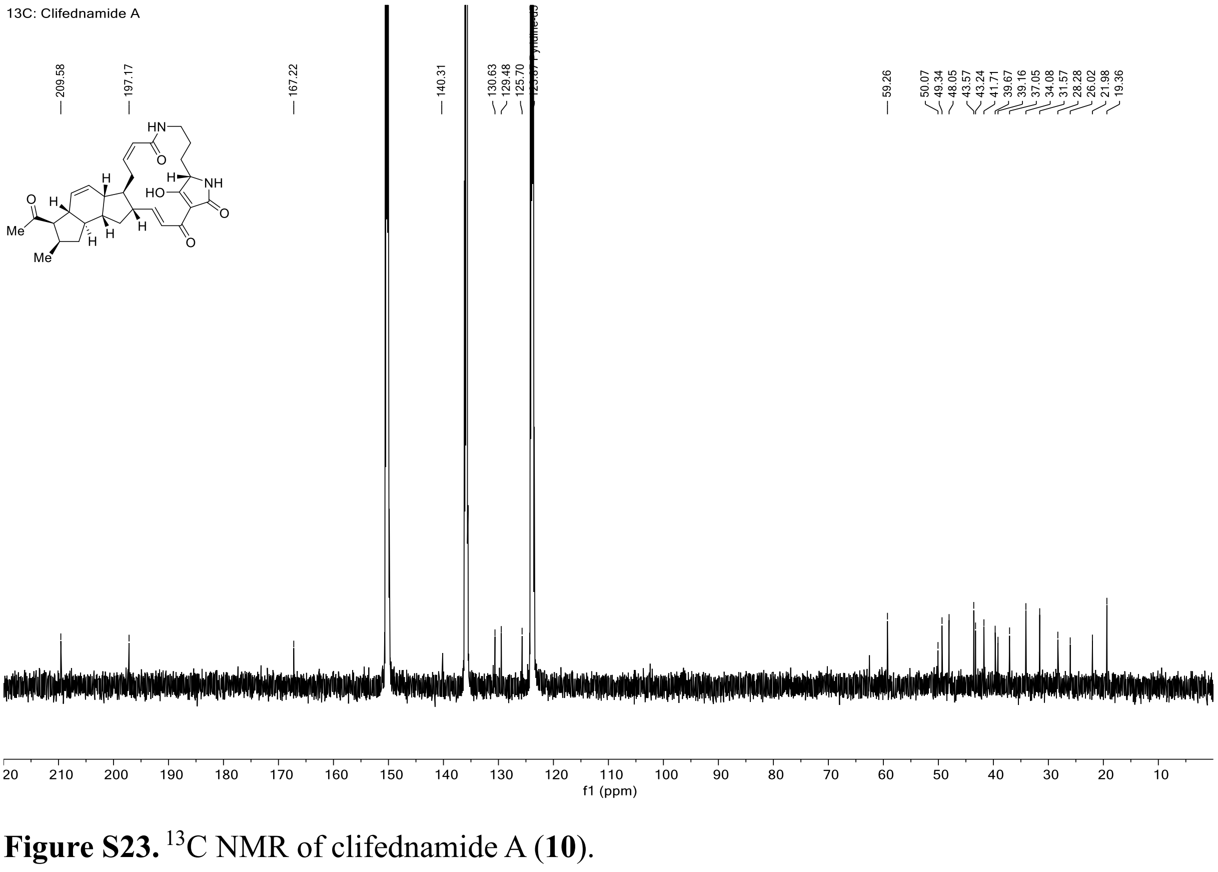

**Figure S23.** $^{13}$C NMR of clifednamide A (**10**).

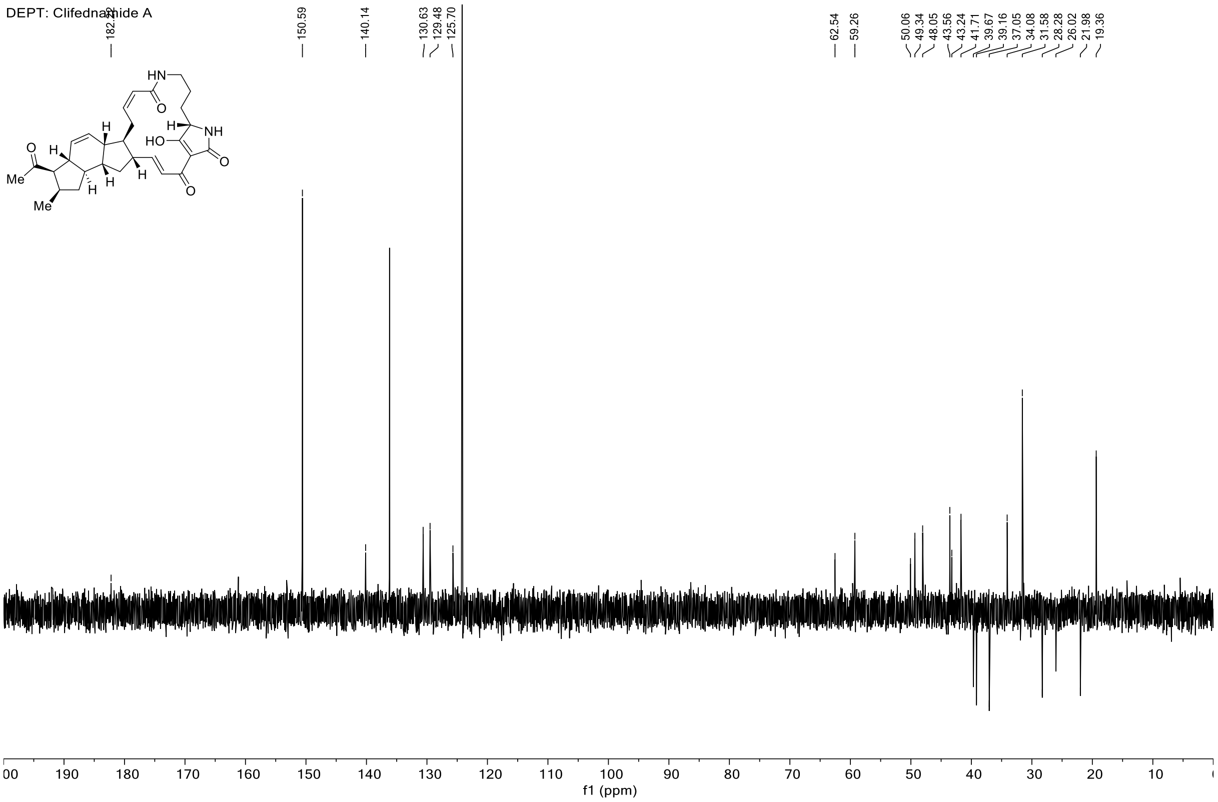


**Figure S24.** DEPT NMR of clifednamide A (**10**).

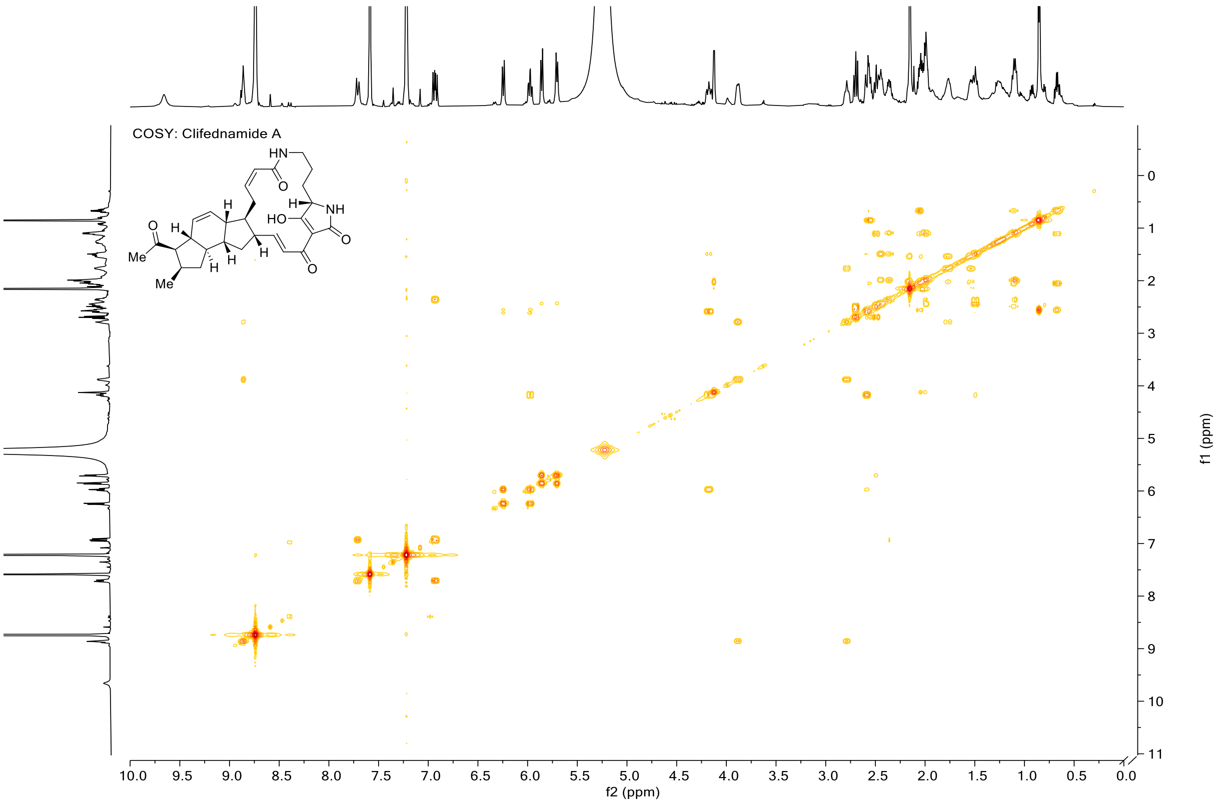


**Figure S25.** COSY NMR of clifednamide A (**10**).

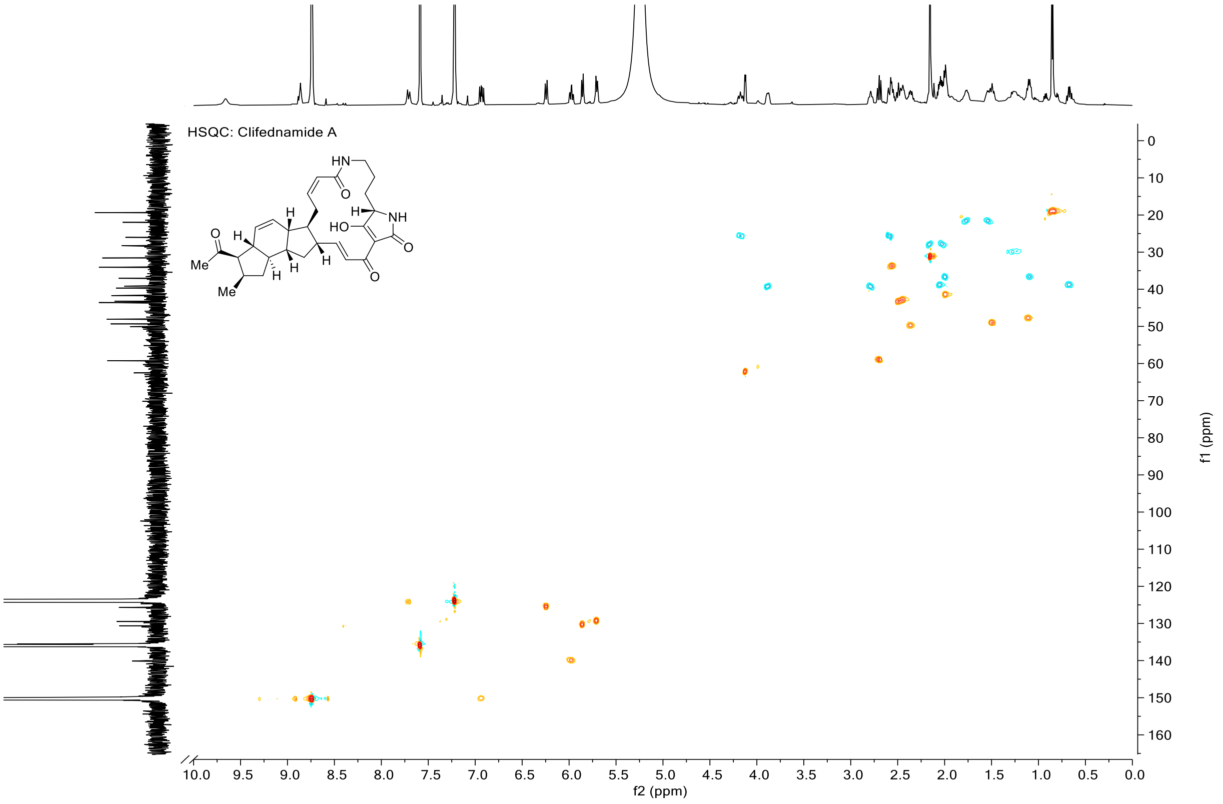


**Figure S26.** HSQC NMR of clifednamide A (**10**).

**Clifednamide C (11)**

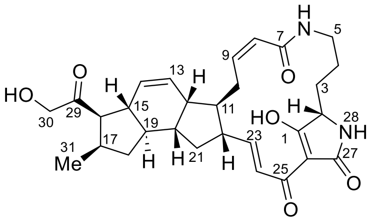


**Table S5.** $^{1}H$, $^{13}C$ NMR (600 MHz, Pyridine-$d_5$) of clifednamide C (**11**).

| | $\delta_C$ | $\delta_H$ (m, $J$) | COSY |
|---|---|---|---|
| 1 | 197.2 | / | |
| 2 | 62.5 | 4.12 (d, 4.9 Hz) | 3 |
| 3 | 28.3 | 2.19–2.12 (m), 2.08–2.01 (m) | 2, 4 |
| 4 | 22.0 | 1.84–1.73 (m), 1.58–1.51 (m) | 3, 5 |
| 5 | 39.2 | 3.91–3.85 (m), 2.82–2.76 (m) | 4, 6 |
| 6 | / | 8.86 (t, 5.7 Hz) | 5 |
| 7 | 167.2 | / | |
| 8 | 125.7 | 6.25 (d, 11.3 Hz) | 9, 10 |
| 9 | 140.1 | 5.97 (td, 11.0, 3.4 Hz) | 8, 10 |
| 10 | 26.0 | 4.18 (ddd, 15.8, 10.6, 4.4 Hz), 2.62–2.53 (m) | 8, 9, 11 |
| 11 | 49.3 | 1.52–1.46 (m) | 10, 22 |
| 12 | 43.2 | 2.47–2.42 (m) | 13, 14, 20 |
| 13 | 129.5 | 5.72–5.69 (m) | 12, 14 |
| 14 | 130.6 | 5.88–5.84 (m) | 12, 13 |
| 15 | 43.6 | 2.49 (t, 11.0 Hz) | 16, 19 |
| 16 | 59.3 | 2.70 (dd, 12.0, 10.1 Hz) | 15 |
| 17 | 34.1 | 2.62–2.53 (m) | 18, 31 |
| 18 | 39.7 | 2.09–2.01 (m), 0.67 (td, 12.0, 7.1 Hz) | 17, 19 |
| 19 | 48.1 | 1.13–1.06 (m) | 15, 18, 20 |
| 20 | 41.7 | 2.01–1.96 (m) | 12, 19 |
| 21 | 37.1 | 2.01–1.96 (m), 1.13–1.06 (m) | 22 |
| 22 | 50.0 | 2.36 (dd, 10.9, 6.5 Hz) | 11, 21, 23 |
| 23 | 150.4 | 6.93 (dd, 15.5, 10.3 Hz) | 22, 24 |
| 24 | 124.3 | 7.71 (d, 14.8 Hz) | 23 |
| 25 | n/a | / | |
| 26 | n/a | / | |
| 27 | 150.3 | / | |
| 28 | / | 9.66 (bs) | |
| 29 | 212.0 | / | |
| 30 | 31.6 | 2.1 (s) | |
| 31 | 19.4 | 0.85 (d, 7.2 Hz) | 17 |

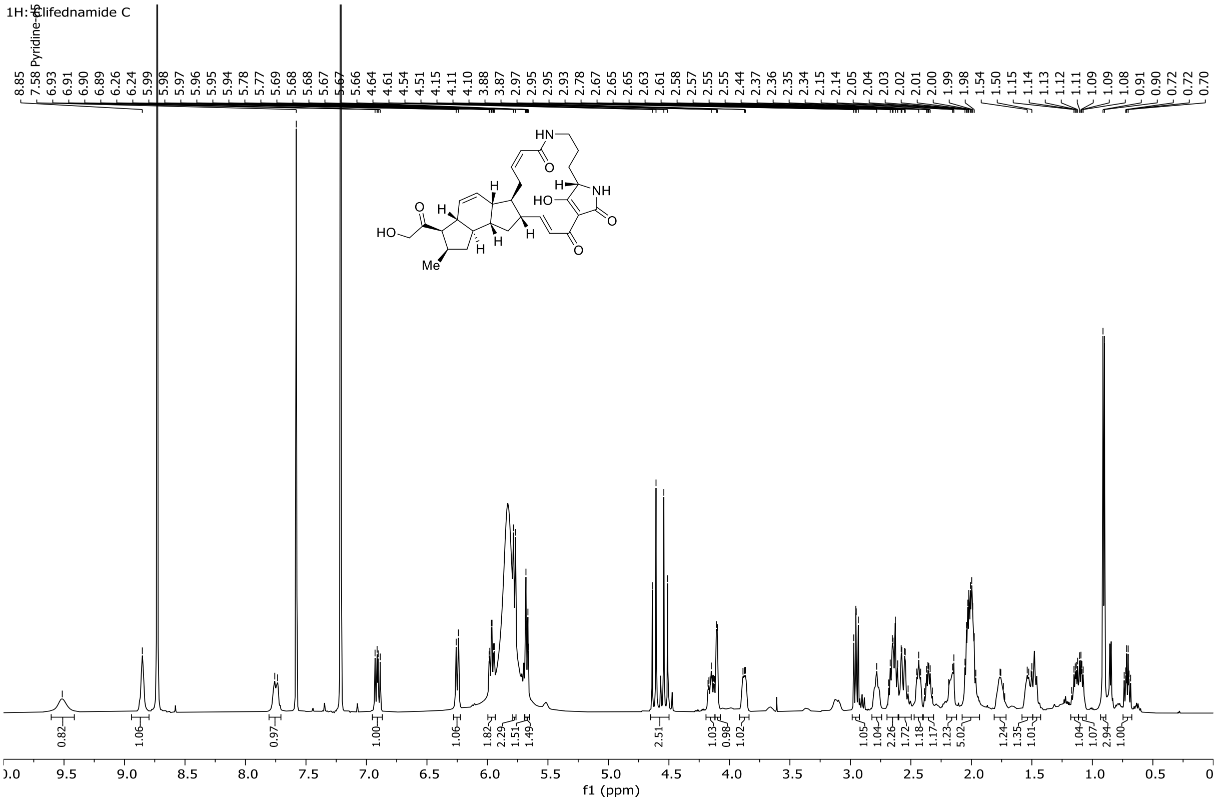


**Figure S27.** $^{1}$H NMR of clifednamide C (**11**).

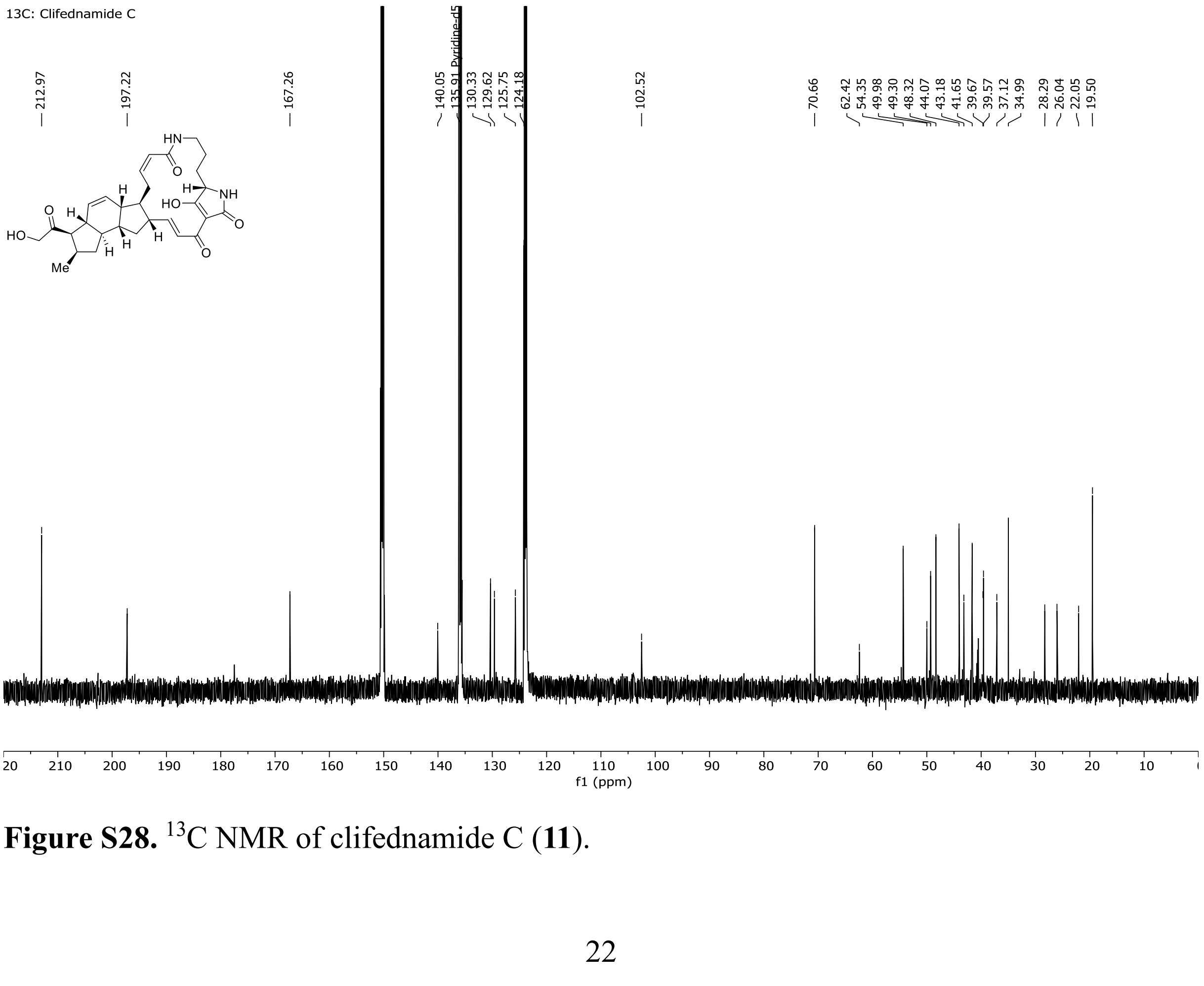


**Figure S28.** $^{13}$C NMR of clifednamide C (**11**).

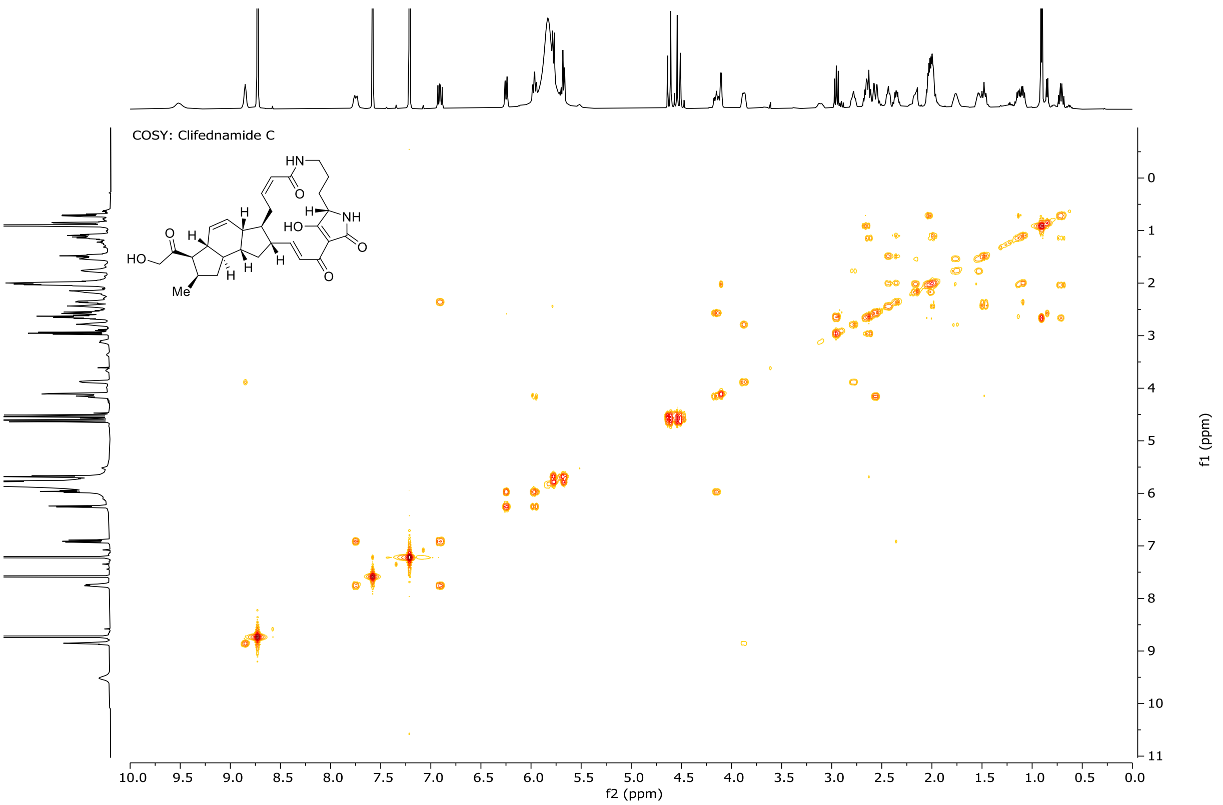


**Figure S29.** COSY NMR of clifednamide C (**11**).

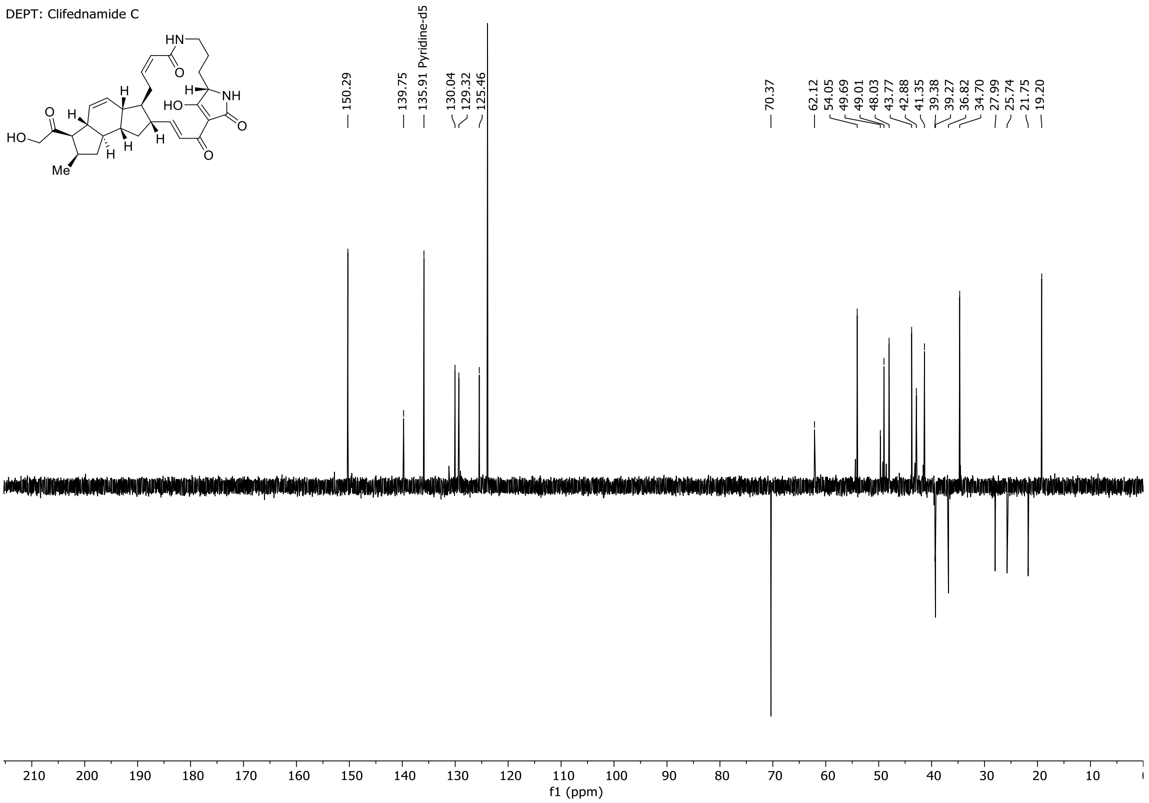


**Figure S30.** DEPT NMR of clifednamide C (**11**).

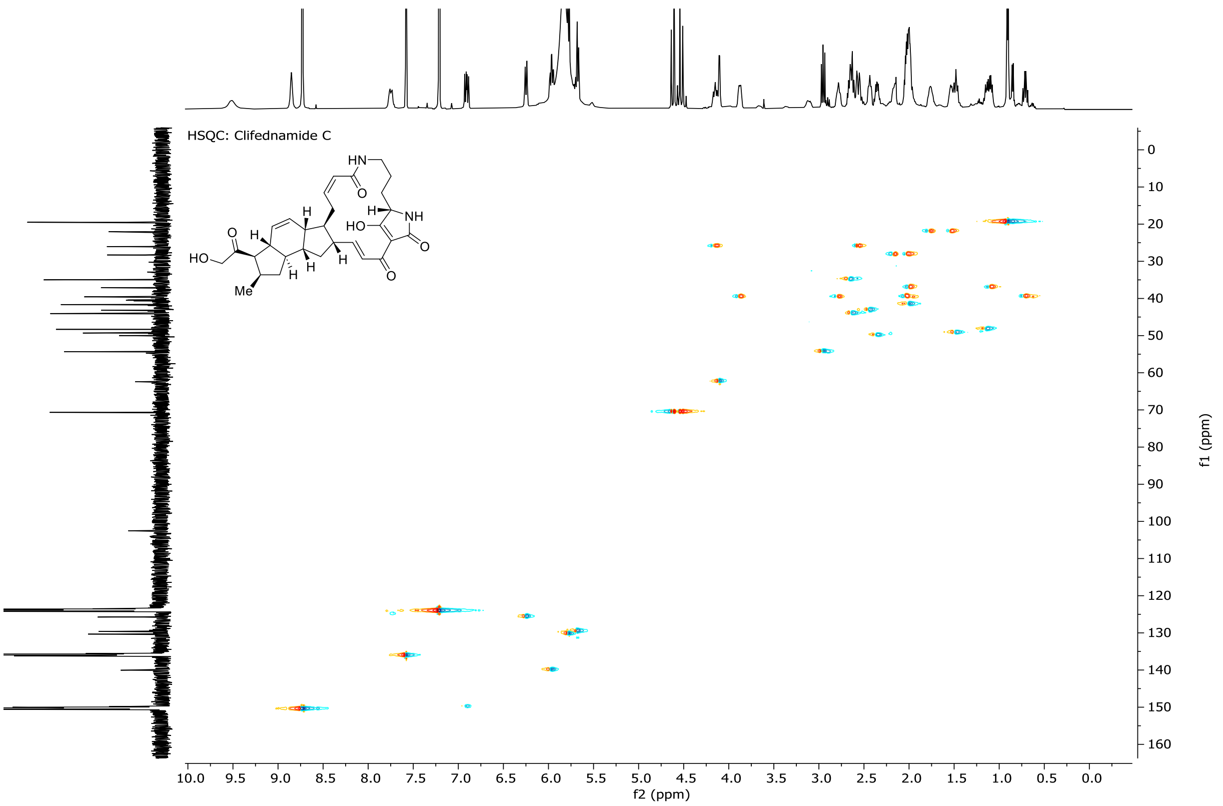


**Figure S31.** HSQC NMR of clifednamide C (**11**).

#### 7. Bacterial strains, primer, plasmids, and sequences

##### Strains

**Table S6.** Bacterial stains used in this study.

| Strain | Usage | Source/Reference |
|---|---|---|
| *E. coli* DH5α | Cloning | NEB |
| *E. coli* ET12567 (pUZ8002) | Donor stain for conjugation | [1] |
| *S. albus* DSM40313 | Heterologous PoTeM expression | DSMZ [2] |
| *S. lividans* TK24 | Heterologous PoTeM expression | [3, 4] |
| *S. coelicolor* M1154 | Heterologous PoTeM expression | [5] |

##### Primers

**Table S7.** Primers used in this study.

| Name | Sequence (5′ → 3′) | Function |
|---|---|---|
| ikaA-fwd | GCAGGTCGACTCTAGAGAGGCCTTCATACC TCGCCATCACC | *ikaA* amplification for pSET152_ermE |
| ikaA-rev | CAACGGAGGTACGGAAGGATGTATTCATGG ATTCCATGCACCACCCTGC | *ika* amplification for pSET152_ermE |
| ikaBC-fwd | CCGTCAAGATCGACCGCAGGCTACAGGGCGA CCAGGAC | *ikaBC* amplification for plug-and-play |
| ikaBC-rev | GTGATGGCGAGGTATGAAGGATGACGCCTT TCGTTCAGC | *ikaBC* amplification for plug-and-play |
| ikaD-fwd | GTACTCTAGACTACCAGGCGACCGGCAGT | *ikaD* amplification |
| ikaD-rev | GCTATCTAGAATGCCCGGACAGCAGGAACA | *ikaD* amplification |
| ptmD-fwd | GTACTCTAGACTACGCGGTGTGGGTCGG | *ptmD* amplification |
| ptmD-rev | GCTATCTAGAATGGAGATTCTCCGCATGGAAG | *ptmD* amplification |
| cftA-fwd | GCTTGGGCTGCAGGTCGACTTCACCAGGCG ACCGGGAG | *cftaA* amplification |
| cftA-rev | GAGCAACGGAGGTACGGACTATGTCGGATC AACACCCCCC | *cftaA* amplification |
| Lac-promoter | CTTCCGGCTCGTATGTTGT | sequencing primer |
| M13-fwd | GTAAAACGACGGCCAGT | sequencing primer |
| ikaA-screen | TTCAACTACCTGATGGGCGA | sequencing primer |
| ikaC-screen | AGCCATCTGGTGTCCACC | sequencing primer |
| ikaC-screen2 | TGTCGGTGGACCCGTATCT | sequencing primer |

**Plasmids**

The basic plasmid of the plug-and-play system only contained the iPKS/NRPS (*ikaA*) as producer of lysobacterene A (**5**) under the ermE* promoter. A second promoter (Table S7) was located downstream flanked by two unique restriction sites (StuI and XbaI) to enable a fast insertion of modifying enzymes. The construct for functional investigation of modifying enzymes on ikarugamycin additionally contained *ikaB* and *ikaC* upstream to the second promoter to enable production of ikarugamycin (**1**).

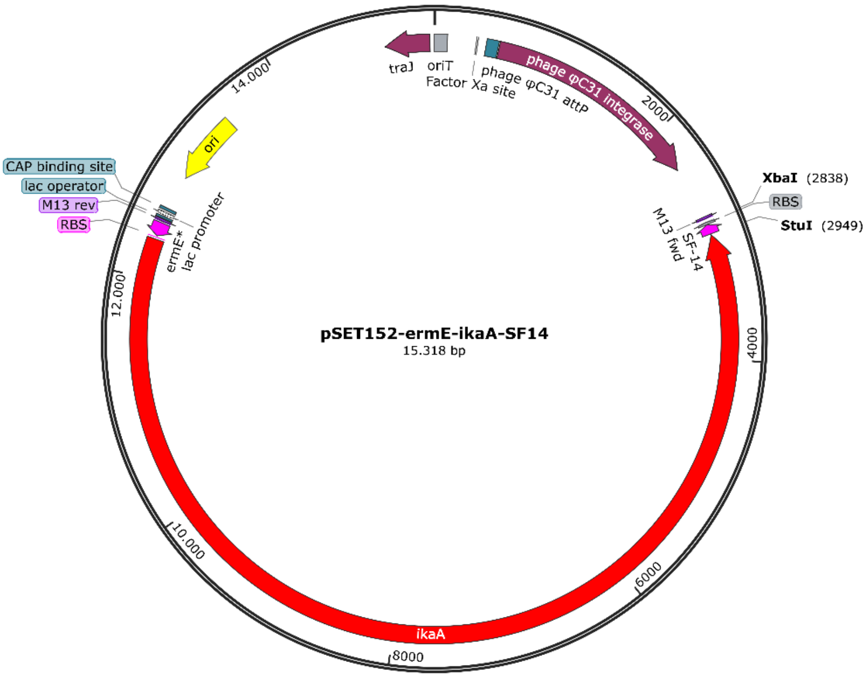


**Figure S32.** Basic expression construct of the plug-and-play system. The plasmid contained *ikaA* as reliable producer of lysobacteren A (**5**), the common precursor of all PoTeMs. Additionally, a second promoter was present to enable sufficient transcription of modifying genes.

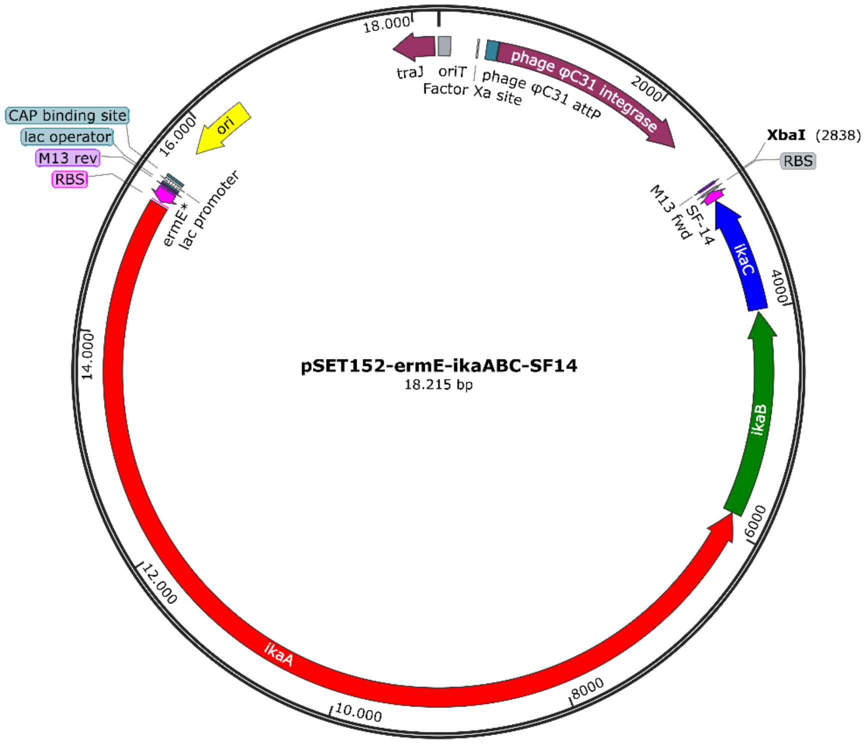


**Figure S33.** Plug-and-play system for the investigation of the influence of modifying enzymes on ikarugamycin (**1**). The plasmid contained *ikaABC* and produced ikarugamycin (**1**) in high yields. No modifying gene was included.

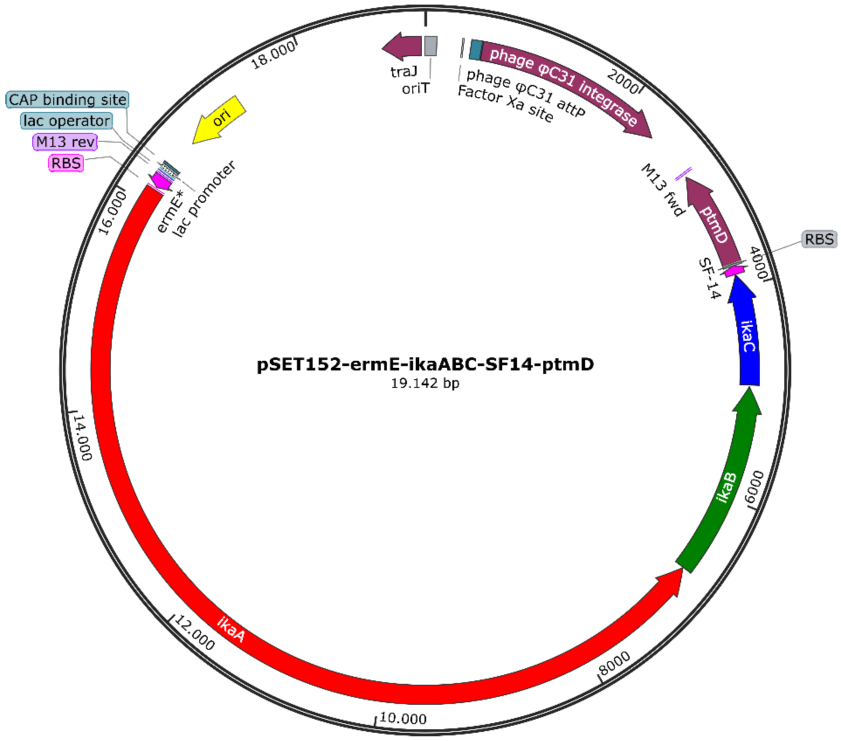


**Figure S34.** Exemplary expression construct for the investigation of the monooxygenase IkaD. The vector contained *ikaABC* as well as *ikaD* under the control of the SF14P promoter.

#### Gene sequences

Gene sequence of the iPKS/NRPS *ikaA* from *Streptomyces* sp. Tü6239 (9375 bp; 336.7 kDa):

```
ATGGATTCCATGCACCACCCTGCCCCCGTCCCCGTACCCGAAGTCCCCGCGCCCGTCCCGTCCCAGGACGACGCG
TTCGCCATCGTCGGCATCGGCTGCCGGCTGCCCGGCGGCGCCAGCGACTACCGGACCTTCTGGCGCAACCTCCTC
GACGGCAAGGACTGCATCACCGACACCCCCGCCGACCGCTACGACACCCGCACCCTGGGCAGCGGCGACAAGGCC
AAGCCCGGCCGGCTGGTCGGCGGACGCGGTGGATACATCGACGGCTTCGACGAGTTCGACCCCGCCTTCTTCGGC
ATCAGCCCGCGCGAGGCCGAGCACATGGACCCCCAGCAGCGGAAGCTCCTGGAGGTCGCCTGGGAGGCGCTGGAG
GACGGCGGCCTCAAGCCCGCCGAGCTGGCCGGCAGCGATGTCGGGGTGTACGTCGGGGCGTTCACCCTCGACTAC
AAGATCCTGCAGTTCGCCGACCTCGGCTTCGAGACCCTGGCCGCGCACACCGCCACCGGCACCATGATGACGATG
GTGTCCAACCGGATCTCGTACTGCTTCGACTTCCGCGGACCCTCGGTCTCCGTCGACACCGCGTGCAGCGGCTCC
CTGGTCGCCGTCCACCTCGCCTGCCAGAGCCTGCGCCGCGGCGAGACCTCCGTCGCCCTGGCCGGCGGCACCCTG
CTGCACATGGCGCCGCAGTACACCATCGCCGAGACCAAGGGCGGGTTCCTCTCCCCCGACGGCCGCTCCCGCGCC
CTGGACGCCTCCGCCAACGGCTACGTGCGCGCCGAGGGCGTCGGCATGGTCGCCATCAAGCGCCTCGCGGACGCG
CAGCGCGACGGCGATCCCATCCACGCCGTCATCATCGGCAGCGGCGTCAACCAGGACGGCCGCACCAACGGCATC
ACCGTGCCCAACCCCGACGCGCAGGTCGCCCTGATCGAGCGGGTCTGCGCCGCCGCCGGCGTCACCCCCGGCAGC
CTCCAGTACGTCGAGGCGCACGGCACCTCCACCCCCGTCGGCGACCCGCTGGAGGCCAACGCCCTCGGCCGCGCG
CTCTCCATCGGCCGCGAGCCGGGCGCCCGGACGTACGTCGGCTCGGTCAAGACCAACATCGGGCACACCGAGTCC
GCCGCCGGCATCGCCGGGCTGATCAAGACGGTGCTCAGCCTCAAGCACAAGGTCATCCCGCCGCACATCAACCTG
GAGAAGCTCAACCCGCAGATCGACGAGGCGTCCCTGCCGTACGAGATCCCCCGCGAGCCCACCCCCTGGCCCGAG
CACAGCGGGCCGGCCCGGGCCGGCGTCAACTCCTTCGGCTTCGGCGGGACCAACGCCCACGTCCTGCTCCAGGAG
GCACCGCCGACCGTCGGGGAGCCCGCGCCACCGGCCACCGACGGGTACTCCGTGCTGCCGCTCAGCGCCCGCGAC
CCCGAAGCCTTTCCCGCCATCGCCACCGGCCTGCGCGAACGGCTCGCCGAGGGACTGCCGGTGGGCGACGCCGCC
TACACCCTCGCCCACCGGCGGCAGCATCTGGAGCAGCGGCTGTCCGTCGTGTACGACTCCCCCGAGGCCCTCGAC
GAGGTGCTCGGCGCCGTCGCCCGCGGCGAGAGCCACCCGCGTGCCGTCGCCGGCACCCAGCGGGAGGGCCTGGAC
CGCAGGCTGGTGTGGGTGTTCACCGGCATGGGCCCGCAGTGGTGGGCCATGGGCCGCCAGTTGTACGCGAGCGAG
CCCGTCTACCGGGAGGTCATCGACCGCTGCGACCAGGAGATCGCCGCGCTCACCGGCTGGTCCCTCACCCAGGAG
CTGAACGCCGACGAGGCCGACTCCCGGATGAGCGAGACCTGGCTCGCCCAGCCCGCCAACTTCGCCGTCCAGATC
GCCCTGGCCGCCCTGTGGCGCAGCAAGGGGATCCAGCCCGACGCCGTCACCGGGCACAGCACCGGTGAGGTCGCC
GCGTTCTACGAGGCCGGGGTGTACACCCTCCCCGAGGCCGTGAAGATCGTGGTGCACCGCAGCCGGCTCCAGCAG
AAGCTCATCGGCACCGGCTCCATGCTCGCCGTCAGCCTCACCGAGGCGGAGGCCGCCCGCCGGGTGCGCCCGCAC
GGCGACCGGGTCTCCATCGCCGCCGTCAACAGCCCCACCTCCATCACCCTGGCCGGGGACACCGAGGCGCTGGAG
GTGATCGCCGCCGAGCTGGGCGCCGAGGACATCTTCGCCCGCTTCCTGGAGGTCGGCGTCCCGTACCACAGCCCC
CGCATGGAGCTGATCAAGGACGAGCTGCTGACCTCGCTCGCCGATCTCAAGCCGCAGCAGGCGAAGTTGCCGCTG
TACCTCACCGCGCTGCCGGGCACCGTCGCGCAGGGCACGGAGCTGGACGCCGACTACTGGTGGCGCAATGTGCGC
GAGGCCGTGCACTTCCGGGCCGCCGTGGACCGGCTGCTGGACGACGGCTACGGCGTCTTCCTGGAGATCGGCCCG
CACCCCGTGCTCGCCCACTCCCTGCGCGAGTGCTGCGAGGCCCGCGACGCGCACAGCGTCACCCTGGCCTCCATC
CGCCGCAAGGCGGACGAGCGCGAACGCCTCACCCTGTCGCTCGCCGCGCTGCACAGCCTCGGCTTCGCCGTGGAC
TGGCACGCCCTGCACCCCGCCGGGCGGCCGGCCGAACTGCCGCGCTACCCGTTCCGGCGCGACCGGTACTGGGTC
GAGCCGGCCCCGGTCGCGCAGATCCGGCTCGGCCACCGCGACCACCCGCTGCTGGGCCGCCGCACCGCGAGCGCC
GAGCCGGTGTGGGAGGTGAAGCTGGACGCGGAGGCCGCCCCGTACCTGGAGGACCACCGCATCCAGGGCACCGTG
CTGTTCCCGGCCGCCGGCTATCTGGAGATGGCCGCGCAGGCCATGCGGGCGCTGACCGGTGATGAGCACAGCACC
GCCGCGCTGGCCGGCATCGAGCTGCGCAAGGCGCTGTTCCTGCCGGACGGCGAGCCGCAGACGGTGCAGCTGTCC
TTCTCCTCCGACGCCGCCGCGTTCTCCATCGCCACCGTGGGCGCCGCCGGCGCCGAGCCGACCGTGCACGCCACC
GGTACGGTACGGGCCGCCCAGCGCCGCCGGCTGACCGCGCCGCTGGACACCGTCGCCGTCCGGGCCCGCGCCGCC
CGCCACCTGAGCGGCCCCGACTGCTACGCCGAACTGGCCGCGCTCGGCTACCACTACGGCCCCGCCTTCCAGGGC
ATCGAGGAGGTGTGGATCGGCGAGGGCGAGGCCCTGGCCCGGATCCGTCCGCCGCAGGGGCTCACCCCGGACGCG
GCGGCGCACCACATGCATCCGGTGCTGCTCGACTCCTGCTTCCAGTCGCTGCTGACCCCGCAGCTGCTCACCGCG
CCCGCCGGGCCCGGGGGCACCGGCATCCGGCTGCCGCTGTCCATCGCCGAGGTACGGCTGGACCCGGTCGGCGAC
CGCGAACTGTGGGTGCACGCCACCGTCACCGGCGACGACGAGGACGAACTCACCGGTGACATCGCCGTGTACGAC
GGCGCCGACGGTACGCCGCTGGGCCGCGTCGCCGGCTTCCGCGCCGCCGATGTGGAGAAGGCCGCCACCACCGTG
GGGCTGTCCACCATCGACAGCTGGCTCACCGAACCGAGCTGGGTGCCGTGCCCGCTGCCCGAGGCGGCGTCCGCC
GCGCCGGCGGCCGGGCGGCACGTACTGTTCGCCGACGCGGGCGGGGTCGCGCAGCGGCTGGCCGCGCTGATCGGC
GAGGCCGGCGGGGAGGCCCATCTGGTCCGGCCCGGTGCCGCGTACGGCCTGGACCGCACGGCGAGGACCGCCACC
GTCGTCCCCGGATCCGCGGATGACCTGCGGCGGTTGCTCACCGATCTCGGGCAGGTGGACGGCGTCGTCCACCTG
TGGAACCTGGACCGGCCGGCGCTGGCCGACGCCCCGCGCGGACGGTTCGCGGACATCGCCTCCACCGGCGCGTAC
GCCCTGATCGCCCTCACTCAGGCCCTGCTCGCCGACCCGGAGCGGCACGGCGGCACCCCGGTGCACATCGTCACC
AGAGCCGCCCAGTGCGTGGTCCCCGGTGAGCCGGTGGAGCCGCTGGGCGCGCCCGCCTGGGGCATCGGCCGGGTG
CTGTGGCAGCAGGAACTGGCCGGGCGCGGCGGCAAGCTGATCGACCTGGCGGCCGACGGCGGCGTCGAGGAGGAC
GCGTACGCGCTGCTGCGCGAGCTGGCCGACCCCACCGGCGCGGCCGAGCGCGAGGACGAGATCGCGCTGCGCGCC
GGGGAGCGGCACACCAGCCGGCTGGTGGCCGCCGAGGGGCTGAGCAGGCCGCTGCCCCTGCGGCTGCGCCCGGAC
```

GGCAGCTATCTGGTGACCGGCGCGTTCGGCGCGCTCGGCAGGCTGCTGTGCCGCACGCTGGTCAGGCGCGGGGCG
CGGCGGCTGATCCTGGTGGGCCGCACCCGGCTGCCGGAGCGCGAGCGCTGGGCCGACCAGGACCCGAACTCGCCG
GCCGGGCGGCACGTGGCCTTCCTCAAGGAGCTGGAGGCGCTGGGCGCGCAGCCGATTCTCGCGCCGCTGGACATC
ACCGACGAGGACGCGCTGGCCGGCTGGCTCGCCGGGTACCGGCGCGCCCAGGGGCCGCCGATCCGCGGGGTGTTC
CATCTGGCGGGGCAGGTGCGCGACACCCTGGTGCCGGAGATGGACCGGGAGGTGTTCGACGCCGTCCACGACCCG
AAGGTGGTGGGCGCGGCGCTGCTGCACCGGCAGCTGAGCGGCGAACCGCTGGAGCACTTCGTGCTGTTCGCCTCG
GTCGCGGCCTGGCTGACGACGGCCGGACAGACCAACTACGCGGCGGGGAACGCCTTCCTGGACGCGCTGGCGCAC
CACCGCCGCGCGCAGGGGCTGCCGGCGCTGGCGCTGGACTGGGGCCCGTGGGCCACCGGCATGATCGAGGAACTG
GGCCTGATCGACCACTACCGCAACAGCCGGGGCATGTCCTCGCTGGCGCCCGAGGCGGGCATGGCGGTGCTGGAG
CGGGTCATCGGGCAGGACCGGGCACAGCTGCTGGTGGCCACGGTCGTGGACTGGCCGGTGTTCATGTCCTGGTAC
GCGGCGCCGCCGCGGCTGGTCACGGAGCTGGCGGCCACCGCCCAGGGACCGGGGTCCGAGGGCGACGGCAGTTTC
CTGGACGCGTTCCGGGAGGCCACCGCGGACAAGCGGCGGCTGCTGCTGACCGAGCGGTTCACGACGCTGGTGGCG
GGTGTGCTGCGGGTGCGGGCCGAGCAGGTGGATCCGGCGGTCAGCCTGAATCTGCTGGGGCTCGACTCGCTGCTG
GCGATGGAGCTGCGAGCGCGGGTGGTGGCCGAGGTGGGCATCGCGCTGCCGGTGGTGGCGCTGCTGTCCAGCGCG
CCGGCCGGGGACCTGATCACCCAGCTGCACGAGGGCCTGGAGGAGTTGCTGGCCGAGGAGGGCAGCGGCGCCGCG
GTGACGGCGGTGGAGCGCTTCGAGGACGAGGCCGAGTTCCCGCTGACGCAGAACCAGAAGGCGCTGTGGTTCCTG
AAGCAGCTGAACCCGGACGGCTTCGCGTACAACATCGGCGGCGCCGTCGAGGTGCGGGTCGAGCTGGACCCGGAC
CTGATGTTCGAGGCGTTTCGCCGGCTGCTGGCCCGGCATCCCGTGCTGCGGGCGAACTTCCTGCTGGTGGAGGGG
CAGGCGGTGCAGCGGATCTCCCCGGAGATCAAGGAGGACATCGCGCTCTTCGACGTCGAGGACCGCGCGTGGGAC
GACATCTACCGGATGATCATCGAGGAGTACCGCAAGCCGTACGACCTGGCGACCGATCCGCTGATCCGGTTCCGC
CTCTTCCGGCGCGGCCCGGACCGCTGGGTCATCACCAAGGCCGTCCACCACATCATCTCGGACGCCATCTCCACC
TTCACCTTCATCGAGGAACTGCTGTCCCTGTACGAGGGGCTGCGGCAGGGCCACGACGTCGAACTGCCGCCGGTG
TCCGCCCGCTATCTGGACTTCCTCAACTGGCAGAACGCGTTCCTGGCCGGCCGCGAGGCGCAGAAGATGCTCGCG
TACTGGCGGGGGCAGCTGCCGGACGAGGTGCCGGTGCTGGCGCTGCCCACCGACAAGCCGCGCCCGGCGGTGCTC
ACCCACAACGGGGCGTCCGAGTTCTTCGCCCTGGACGCGGAGTTGAGCGCCCGGGTGCACGCGCTGGCGCGGGAG
CACAACGTCACCGTCTTCATGGTGCTGCTGAGCGCGTACTACCTGCTGCTGCACCGCTATGCGGGGCAGGACGAC
ATCATCGTCGGCTCCCCCGTCACCGGCCGCACCCAGGAGGAGTTCGGCGCCGTCTACGGGTACTTCGTGAACCCG
CTGCCGCTGCACGCCTCGCTGGCCGGTGACCCCACGGTCGCCGAGCTGCTGGACCAGGTGCGCACCACGGTGCTG
GGCGGCCTGGACCACCAGGAGTACCCGTTCACGCTGCTGGTGGAGCAGCTGGGGCTGGCCCACGACCCGAGCCGG
TCGGCGGTCTTCCAGGCGATGTTCATCCTGCTGCACCACAAGGTGGCCACCGAGAAGTACGGCTACAAGCTGGAG
TACATCGAGCTGCCCGAGGAGGAGGGCCAGTTCGACCTGACGCTGTCCGCGTACGAGGAGGAGGCGGACGGGCGG
TTCCACTGCGTCTTCAAGTACAACACCGACCTCTTCGAGGCGGAGACGATCCGGCGGCTCGCCGGGCACTACACG
CAGCTCCTGGAGTCGCTGACCGCGGCGCCCGCCGACGCCGCCACCGGTGGACTGCGGATGCTGTCGGGCGGCGAG
CGGGAGCGGATCCTCACCGAGTGGAGCGGGGCCGGGCAGGGCGCGCAGGACGCGCCGGTGCCGGTGCACCGGCTG
ATCGCCGAGGCGGCGCACCGTACCCCGCAGGCGATCGCGGTGGCCGCGCCCGCCGAGAGCGGGGAGACCCGGCGG
CTGACGTACGGCGAACTGGAGGAGCGCGCCGGCGAACTGGCCGGGCGGCTGCGGGCGCGCGGCGTGCGCGAGGGC
ACCGTCGTCGCGCTGTGCCTGGAGAAGTCGCCCGAGCTGATCACCGCCCTGCTGGCGGTCCTCAAGGCGGGCGGC
GCCTATCTGCCGCTGGACCCGGACTATCCGGCCGACCGGCTCGCGTACATGGTGCGCAACGCCGGGGCCACGCTG
GTGATCGGCGGGACGGGCGGCGCGGCCGAGGGGCTGCCGGGCACCGTGGTCACCCTGGAGGAACTGCTCGCGGGC
GAGGCCGGCGAAGCGGGGCCGGACGCCGAGCCGGGGCCCGACTCCCCCGCCTACGTCATCTACACCTCGGGCTCC
ACCGGGCGCCCCAAGGCGGTCGCGGTCAGCCACCGCAATCTGGCCTCGGTGTACGCCGGATGGCGCGACGCCTAC
CGCCTGGAGGAGGGCGGCATCCGGGTCCATCTCCAGATGGCCAGCCCCTCCTTCGACGTCTTCACCGGCGACCTG
ACCCGAGCCCTGTGCTCGGGCGGCACGCTGGTGCTGGTCGGCCGGGAGCTGCTGTTCAACACCGCCCGGCTGTAC
GAGACGATGCGCGCCGAACGGGTGGACTGCGGCGAGTTCGTGCCCGCCGTGGTGCGCACCCTGGTGCGGCACTGC
GAGGACACCGGCGCCCGGCTGGACTTCCTGCGGCTGCTGATCGTGGGCTCGGACTCCTGGAAGGCCGAGGAGTAC
GAGCGGCTGCGCGCGCTGGGCGCACAGCGCCTGGTGAACTCGTACGGGCTCACCGAGGCCACCATCGACAGCGCC
TGGTTCGAGGGTCCCGCGGATGACCTGGAGGGCGGCCGGATGGTGCCCATCGGGCGGCCGTTCCCGGGCAGCGCG
CTGTACATCCTGGACTCGCGCGGCGAGCCGGTGCCGCCCGGTGTCCCCGGCGAGCTGTGGATCGGCGGCACCGGG
GTGGCGCTCGGCTACCTCGGCGACGAGGCGCTGACCGGGGAGCGGTTCCTCACCCGCGCCCTGGCCGGCGACGCT
CCGGTACGGCTGTACCGCACCGGTGACCTCGCGCGCTGGGACGCGGCCGGCACCGTCCATCTGCTGGGCCGGGCC
GACTCGCAGATCAAGGTGCGCGGGCACCGCATCGAGATCGGGGAGATCGAGTCGCACCTGGCGGCCTGCCCCGAG
CTGGCCCAGGCGCAGGTCACCGTGCGGCCGGACGCGGGCGGCGAGAACGTGCTGTGCGCGTACGGGGTGGCGGCC
CCGGGCGCCGTGCTGGACTGGCGCGAGGTGCGCCGGCGCCTGGCGGACTATCTGCCGACGTTCATGATCCCCACC
CACTTCACCGAGCTGCCCGCCCTGCCGCTCACCCCGAACGGCAAGGTGGACGTGGCGGCGCTGCCCGCCCCGCGC
ACCGGCGACGGCGCGGACGGGCCGGTGTACGAGGCCCCCGTCACGCTGTACGAGACCCGGATGGCCGAGCACTGG
CAGCGGCTGCTGGGCATCGAGGCCCCCGGGCCCGGTCTGGGCCACGACTTCTTCGAGACCGGTGGCAGCTCCATC
CGGCTGATCGAGCTGATCTACCACCTGCAGGCCGAGTTCGGGATCTCCATCCCGGTCAGCCGGCTGTTCCAGGTG
ACGACGCTGCACGGCATGGCCAAGACGGTCGAGCGGATCGTCACCGGGGAGATCGAGGGGTCGCTGCCGTATCTG
CGGTTCAACGAGAACGCCGCGGCGGGCACGGTGTTCTGCTTCCCGCCGGCCGGTGGCCACGGCCTGGTCTACCGG
GAGTTCGCGGCGCGGCTGCCGGAGTTCGAGTTCCTCGCCTTCAACTACCTGATGGGCGAGGACAAGGTAAGCGGG
TACGCCGACCTGGTGGCCGGGCACCGGCCGGAGGGCGAGATCGACCTGCTCGGCTACTCGCTGGGCGGCAACCTC
GCCTTCGAGGTGGCCAAGGAGCTGGAGCGGCGCGGCCGCACCGTGCGCCACGTCGTCATCATGGACTCGCTGCGG

GTGACGGAGTCCTACGAGCTGGGCCCGGAGCACCTGGCCGTCTTCGAGCGCGAGCTGGCCGAGCATCTGCGCAAG
CACACCGGCTCGGCGCTGGTCGCGGAGAAGACGCGCGAACAGGCCAAGGACTACCTGGAGTTCACCGGCCGCACC
GCCAACCCCGGCACCACCGGGGCCCGGATCGCGGTGATCAGTGACGAGGAGAACGCGGCCGCGTACGACAGCGGC
GCCGAGGGCAGCTGGCACGGCGCCTCCCGTACCGGAACCGACGTGCTGCGCGGGGTGGGCCGGCACGCCGACATG
CTCGATCCGGGGACGGTCGAGCACAACGCGCGCCTGGCGCGCGGCATTCTCACCGGCGGTGATGGCGAGGTATGA

Gene sequence of the FAD-dependent oxidoreductase *ikaB* from *Streptomyces* sp. Tü6239 (1833 bp; 67.5 bp):

ATGACGCCTTTCGTTCAGCCGGCGGTCGACACCAAGGAGCACAGCGCCATGTCATCCCCCACCACCTCCGGCACC
CCGGGCAGGCAGTCGATGATCATCATCGGCGGCGGCCTGGGGGGCCTGTCCACCGGCTGCTACGCGCAGATGAAC
GGCTACGCGACGCGGGTCTTCGAGATGCACGAGATCCCGGGCGGTTCCTGCACCGCCTGGGAGCGCGGGGACTTC
ACCTTCGACTGGTGCGTCAGCTGGCTGCTGGGCAGCGGTCCCGGCAACGAGATGTACCAGATCTGGATGGAACTG
GGGGCGTTGCAGGGCAAGGAGATGCGCCAGTTCGACGTCTTCAACATCGTGCGGGTGCGCGGCGGCCAGCCGGTG
TACTTCTACTCCGACCCGGACCGGCTCCAGGCGCACCTGCTGGAGATCTCCCCGGCCGACGCCCGCCGCATCAAG
AACTTCTGCGAGGGGGTGCGCACCTTCCAGAAGGCGCTGTCGGTCTACCCGTTCCTCAAGCCGGTGGGGCTGATG
GGGCGGTGGGAACGGTGGAAGATGCTGGCCTCGTTCCTGCCGTACTTCAACGCCATCCGCAAGTCCATCACCGAG
CTGATGACGGACTACGCGGAGAAGTTCCAGCACCCGGTGCTGCGCGAGGCCTTCAACTACGTGCTGTACGAGAAG
CACGCCGACTTCCCCGTCCTGCCGTTCTGGTTCCAGCTGGCCTCGCACGCCAACGGCTCGGCGGGGGTGCCCGAG
GGCGGCTCGCTGGAGCTGGCCCGGTCCGTGGAGCGGCGCTACCTGGGGCTCGGCGGGGAGATCACCTACAACGCC
AAGGTGGAGAAGATCCTCGTCGAGCACGACAAGGCGGTGGGAGTGCGGCTCACCGACGGCCGCGAGTTCCGCGCG
GACATCGTGGTGTCGGCGGCCGATCTGCACACCACCGCCATGGAGATGCTCGGCGGCCGGTATCTCAACGACACC
TGGCGCAAGCTGCTCACCGAGACGATCGACGAGGTGGGCACGATCTCCCCCGGCTATGTCTCGCTGTTCCTGGGG
CTGCGCCGGCCGTTCCCCGAGGGCGAGCCGTGCACCACGTACGTGCTGGAGGACAGCATGGCGGAGAAGCTCACC
GGCATGCGGCATCCCAGCATGAACGTGCAGTTCCGCAGCTGCCACTACCCGGAGCTGTCGCCGCGCGAGACCACG
GTCATCTTCGCCACGTACTTCTCGGAGGCCGAGCCGTGGCGGGCGCTGCGCGACGACGTGCCGGAACAGGCGGGC
CGGGTGCGGCGCGGTCAGGTGCTGCACACCCTGCCGGTGAAGCACGGCAAGGCGTACACCCAGGCCAAGCGGCAG
GCGCGGATCACCATCGAGAACTTCCTGGACGAGCGGTTCCCCGGTCTCAAGGACGCGGTCGCCGTGCGGGACGTG
TCCACGCCGCTGACGCAGGTGCGCTACACGGGCACCTACAACGGCGGGTTCCCCGGCTGGCAGCCGTTCGTGGAC
GGCGGGGAGACCGTGGAGGTGGAGATCAACAAGAACGGCCCGGTGCTGCCGGGGCTCTCCAACTTCTATCTGGCC
GGGGTGTGGGTCACCGTCGGCGGGCTGATCCGGGCGGTGGCCTCGGGCCGGCAGGTCACGCAGGTGATCTGCCGG
GACGACGGGCGGGAGTTCACGGCGAGCGTGGACGAGAGCGCGCCGCCGCCCACCCAGGTCGCCATCCCGGTGGGC
AAGCAGCCGGGCGTGCCGGATCTGGCGGCCGGGTTCCCCGCCCAGACCGCCGGCGGCGAGACCGCCACCGGCGCC
AACAACACCGTCACATCGTCGAGGAGCGTGTAG

Gene sequence of the alcohol dehydrogenase *ikaC* from *Streptomyces* sp. Tü6239 (1047 bp; 37.5 kDa):

GTGATCGCCGAGCACATCCCGGGGGTCCCGGACACGGACCGGATCTACCGGAAGGTGACGCGGGAGTTCGATCCC
GCCTCGCTGGCGGACGATCAGATGCTGCTGCGCACCCGGTACGTGTCGGTGGACCCGTATCTGGTGGGGCTCTCG
CTCCAGACGCCGATCGGGGACACCGTGCGCGGTGACTCGATCATGGAGGTGGCCGTGGCGGGGCCGCGCGCCCGC
TTCCAGGTCGGGGACCTGGTGCAGGGGTACGGCGGCTGGTGCAGCCATCTGGTGTCCACCGGGGGGCCCAGCGGA
TGGAACGACGACGGCGCCGAGTTCGCCGTCCAGTTGCCGCCGTTCCGCAAGCTGGACCCGCGGCGGTACGACGAG
GCGCTGCCGCTGTCCACGGCGCTGGGCGTGATGGGCACCCCGGGCATCACCGCGTTCGGCGCGATGAAGACGTTC
CTGACCGTGGGCTCCGAGGACACGGTGGTGATCAGCGGGGCGTCCGGGACGGTGGGCACCCTGGTGGGCCAGCTC
GCCAAGCGGGCCGGGGCCCGGGTGGTGGGCACCACCTCCTCGCCGGGGAAGGCCGCGTATCTGACGCAGCTGGGC
TTCGACGCGGTGGTGAACTACCGGCAGGGCGACGACACGGACACGGTGCGCGAGGCGCTGGCGGCGGCAGCGCCC
AACGGAATCGACAAGTACTTCGACAACCTGGGCGGCACCGTGACGGACGCGGTGTTCACGATGCTCAACGTGCAC
TCCCAGGTGGCGGTGTGCTGGCAGTGGGCCACCACGGTCAACGGGGACTGGACGGGGCCGCGGCTGCTGCCGTAC
ATCATGTTCCCGCGCACCACGATCCGGGGGATCTTCGCCGACGAGTGGTACACGGAGGAGATGGTCGACGCGCTG
CACGAGGAGGTGGGCGGGCTGATCCGCAAGGGTGAGCTGGCCTACCACCAGACCATCCACCAGGGCTTCGACGCC
CTCCCGGACGCGTACCGCTCCCTGTACACCGGCCAGGAGGGCAACCGCGGCAAGGTCCTGGTCGCCCTGTAG

Gene sequence of the monooxygenase *ikaD* from *Streptomyces* sp. Tü6239 (1224 bp; 45.7 kDa):

```
ATGCCCGGACAGCAGGAACAGCAGGCACCGTCGGAGCACCCGGAGCAGGAACTCCTCACCTTCCCCTTCCCCTCG
ACGGGCCTGGAGTTTCCCCCCGTCTACCACGAGCTGTACCAGCAGCGGCTCACCAAGGTCCGGCTCCCGTACGGC
GACGACGCCTATCTCGCCATCCGGTACGCCGATGTGAAGACCGTGCTGTCCGACTCCCGGTTCTCCATCGTCGCC
TCCCTCGGCCAGGACCAGCCGCGCACCCGGGCCGGGGCCCGCGTCGGCAACGGCCTGTTCTCCCTCGACCCCCCG
CAGCACTCCCGGCTGCGCTCCGTCCTGGGCCGGGACTTCACCCCGCGCCGGGTGGAGAAGCTGCGCGAGCGGGTG
CGGGAGCTGACCGACCAGTGCCTGGACCGCATGGAGGCCGCCGGGTCCCCCGCCGATCTCGTCGCCCACCTCGCC
GTGCCGATGCCCACCGCCGTGGTCTGCGAGATGATGGGCGTACCCGAGCCCGACCACCACCTGTTCTGGGGCTGG
GCCGAGACGATCCTGTCGAACGACACCACGCCCGACGACCTCATCCGGCGCTACCAGGAGTTCACCGCCTACATG
GGCGGCATGGTCGAGGAGCGCCGCGCCCGTCCCACCGACGACATGTTCGGCATGCTGGTGCGGGCCTGCGACGAG
GAGGGCCGGATCACCGAGATCGAGATGCACGCGCTCGCCTCCGACCTGCTCAGCGCCGGCTTCGTCAGCACCGCC
CACCAGATCGCCAACTTCACGGCCATGCTGCTGGCCCGCCCCGAACGGCTCCAGCCCCTGGTGGACAAGCCGGAG
CAGATCCCGGCCGCCGTCGAGGAACTCATGCGCCACGTCCCGATCCTCAGTGGCTTCTCCTTCCCCCGCTACGCC
ACCGAGGACCTGGAGATGAGCGGGGTGACCGTCCGCCGCGGCGAGGCCGTCATCCCGGTGATCGCCGCCGCCAAC
CGCGACCCCGACGTCTACCCGGACGCGGGCCGCCTCGACCTCGAACGCAACGGGCTTCCGCACCTCGGCTTCGGC
CAGGGCCCGCACTTCTGCATCGGCGCCCATCTGGCCCGGGTCGAGCTGCAGGTGGTCCTCGAAGCCCTCACCGAG
CGCTTCCCCGACCTGCGCTTCGGCGTCCCCGAGAACGCCCTCAAGTGGAAGCAGGGCCACTTCATGAACGGCCTG
CACGAACTGCCGGTCGCCTGGTAG
```

Gene sequence of the sterol saturase *ptmD* from *S. pactum* SCSIO 02999 (921 bp; 33.4 kDa):

```
ATGGAGATTCTCCGCATGGAAGATCGACCCGATACGGGGTCGCTGCCGGCGAATGTGTGGACCGCCGGCGCGTCC
GCCCGCGAGTGTCTCCGCTATGCCGCGTACCCGATTCTGCTCTTGTCCGCAGTCTGGCTCTTCGCCTCGGTTCTG
CGCTTCGGCTGGGACCGGGGCCAGGCCATCCAGCTCTTCCTGATCGGCACCATCGTCTATCTCGCGGCGCTGGAG
CGGCTGATCCCGCACCGGACCGACTGGCACCCGAGCGGTCGGGAACTGTGCTGGTACGCAGCCTATTTCGGGTTC
ACCATGGTCGGCGCCGTGCTCGGCCAGTCGGTCGTCGCGGCCGTTCTGGCCGCGACCCCCGTCACGGGACCGGGG
CTCGCGCTCGGGGCCGAGGTGCCCCTCGCGCTGCTGGCCTCCTCACTGACCGGCTACCTCGCCCACCGCTGGGCG
CATTCCAATCGGTGGCTGTGGAAGGTACACGGAATTCATCACGTCCCCGGAAAGGTGAACGTCGCCAATAACGGC
GTCAACCACCTACTGGATGTCGCCTTCAAACAGGGCGCGGTCCAATTGACACTGGGATTTCTCGGATTCTCCGCC
GACTGCCTTTTCGCGGTGGCGCTCTTCACCCTTGTCCAAGGATATTTCGTGCACGCGAACGTGGACGTCCGACTC
GGTCCGCTCCACCACGTACTGGCCAGCCCCGAGCAGCACCGGCTGCACCACAGCGCCGACCTGGCCGAGGCCGGT
CACTTCGGCGTGGACCTCTCGGTCTGGGACCGCCTCTTCGGCAGTTTCACCTGGCGACCGGACCGCCGCCCGGCG
ATCGTCGGTGTCAAGGACCCCGCCACGTTCCCGGAGACCGGCTCGATCGTCGCCAGCCTGCTCCACCCCGTGCGC
CGCCCGACCCACACCGCGTAG
```

Gene sequence of the monooxygenase *cftA* from *Streptomyces* sp. JV178 (44.0 kDa; 1191 bp):

```
ATGTCGGATCAACACCCCCCACTCCCCTACCCCTTCGCCCCACGCGGCCTCGACCTCGACCCCACCTACGCGGAG
TTGCGCGACCGCGCACCGGCCCGTATCCGTATGCCGTACGGCGACGACGCCTGGCTGGTCACCCGGTACGAGGAC
GTACGGACGGTCCTGGCCGACCCGCGGTTCAGCCTGGCCGCCTCGATGGGCCGGGACCAGCCGCGGATGCGCCCG
GTGGCCCGTACCGGCGCCGGCCTGTTCTCCACGGAGCCGCCCGACCACACCCGGCTCCGCTCACTGGTCGCACGG
CAGTTCAGCGCCCGCCGGGTCGAGCCGCTGCGCGCACGGGCCGGGGAGCTGGCCGACGAGCTGATCGACGGGATG
GTGGCCGCAGGCCAGCCGGCCGACCTGGTCGAGGACTTCGCCATCCCCATGCCGACCACCATCATCTGCGAGGTG
CTCGGCATCCCCGCCAAGGACCACCGGATGCTCTGGCACTGGGCGGAGACGGTGCTCTCCGCCATCACCCCCCAG
GAGGTGCTCGCCACCGAGGGCCGGGCCTTCATGGAGTACATGGTCGGCGTGCTGGAGCTGCGGGGCCGCGAGCCG
GGCGACGACCTGCTCACCACCCTGGTCCGGGCCTGCCGCGAGGAGGGCCTGATCAGCGAGGAGGAGCTGCTGTCG
ATCGCCTGCGACCTGCTGATCGCCGGCTTCGTCTCCACCACCAACCAGATCGGCAACTTCTTCCACCAACTCCTC
GTACATCCAACGGAGTTGACGCGGCTACGGGAGCGCCCCGAGCTGATCCCGAAGGCCGTAGAGGAGCTGATGCGC
TACGTGCCACTGCTCACCGGCTTCAACCTGGCACGGTACGCGACCGCCGATGTCGAGCTGGGCGGCATCACGATC
CGGGCCGGCGAGGCGGTGATGATCGCCACCGCCGCCGCCAACCGCGACCCGGGGGTGTTCCAGGAGCCGGAGCGC
CTCGTCTTGGACCGCGACGCCAACCCGCACATCGGCTTCGGCCACGGCGTGCACTACTGCGTCGGCGCCCATCTG
GCCCGGCTGGAGCTCCAGGTGGCGATCGAGCGGGTGCTGCACCGGCTGCCCGGCCTACGGCTGGCCGTCCCCGAG
AGCGAACTGAGCTGGAAGCAGGACGCGATGGTCAACGGTCTCCAGGCACTCCCGGTCGCCTGGTGA
```

#### Sequences of the used promoters

**Table S7.** Strong, constitutive active promoter sequences used for plug-and-play system. The promoter sequences were elongated by restriction sites (underlined; StuI: AGGCCT and XbaI: TCTAGA), and RBS (bold) were added if not already present.

| Promoter | Sequence |
|---|---|
| **ermE** | AGGCCTGCGGTCGATCTTGACGGCTGGCGAGAGGTGCGGGGAGGATCTGACCGACGCGGT<br>CCACACGTGGCACCGCGATGCTGTTGTGGGCACAATCGTGCCGGTTGGTAGGATCCAGCG<br>GAGCAACGGAGGTACGGATCTAGA |
| **gapdhP(EL)** | AGGCCTGCTGCTCCTTCGGTCGGACGTGCGTCTACGGGCACCTTACCGCAGCCGTCGGCT<br>GTGCGACACGGACGGATCGGGCGAACTGGCCGATGCTGGGAGAAGCGCGCTGCTGTACGG<br>CGCGCACCGGGTGCGGAGCCCCTCGGCGAGCGGTGTGAAACTTCTGTGAATGGCCTGTTC<br>GGTTGCTTTTTTTATACGGCTGCCAGATAAGGCTTGCAGCATCTGGGCGGCTACCGCTAT<br>GATCGGGGCGTTCCTGCAATTCTTAGTGCGAGTATCTGAAAGGGGATACGC**CGAGCAACG**<br>**GAGGTACGGAC**TCTAGA |
| **rpsLP(XC)** | AGGCCTGCCCTGCAGGCGGAAGTCAGGTAGACACGACTTCCGCTAGTCCTTGCAAGGTCT<br>GCTGACGTGAGGCGGGGCGGTCGTTTTTGACCGCCCCGCCTTCGTCATGTAGGCTCGCTC<br>GCTGTGCCTGGCGTGTCTTCAGACGCCCAGGTCCCGGTGCCGTGAGGCCCGGGCCATCGA<br>GCCGGTGGTACGTGGCTGCGGTCCCCTTGTGAGGGCTGCGCGCCGTGTGCTGTCCGGCGC<br>GCACAGCCTTGAATCCACCCGCGGGGGCCGGCCGGTCTCCGTGAGCTCGAGAAGACGACG<br>GAGACGTA**CGAGCAACGGAGGTACGGAC**TCTAGA |
| **kasOP*** | AGGCCTTGTTCACATTCGAACGGTCTCTGCTTTGACAACATGCTGTGCGGTGTTGTAAAG<br>TCGTGGCCAGGAGAATACGACAGCGTGCAGGACTGGGGGAGTT**CGAGCAACGGAGGTACG**<br>**GAC**TCTAGA |
| **SF14P** | AGGCCTCCTATCCAGGAGATATTATGAGTTACGTAGACCTACGCCTTGACCTTGATGAGG<br>CGGCGTGAGCTACAATCAATACTCGATTA**CGAGCAACGGAGGTACGGAC**TCTAGA |
| **P-2** | AGGCCTGCCCGGCCATATCCGGCCCGGCCAAATCTCGGCCGGCCACCTCGGCCTGGCCAG<br>CCTGGCCCGGCCAATCTCGGCCCGACCAACTTCAGCCCGGCCGGCGCTTGAGGCCGATGA<br>GCCGCGGAGCGGCGAGTCTTCCGCCCGGCCGGTCCGGGTGGCCTCAAGCGCCGGCCGGGC<br>TGGTTTTGGTGCGGACACGTCTGACCGTGCCGGTCACCGATGGCCTCAAGCGCCGGCCGG<br>GCTGGGAGTGGTGGCCGAGGCTTCGGGCGTACGTGCCAGCCCGCAAGGGGCTGCGGTGGG<br>GTGGCCTCAAGCGCCGGCCGGGCTGAGGTTGGCTGGCTGGGCCGGGTTCGGCCGGTGGGT<br>CGAGGTGGCCTGGCCGGGCTCGCCAGGGTGAGTTGGCCGACGGGCCGAGGCGGCCCGCCC<br>GGGCTCCCCGGGCCGAGTTGGCGCGGCCAGGCCAGGGCTCAGCAGGGTGGGGGAGTGGGG<br>CAGGCGGCCCGGTAGGGGAGTGCGGGAGGGCAGCGCGCGCCGCGCGCATTGGCACTCCGC<br>TTGACCGAGTGCTAATCGCGGTCATAGTCTCAGCTCTGGCACTCCCCGCAGGAGAGTGCC<br>AACACAGCGACGGGCAGGTCCGGCACCCGCGACGACGGATCGACCTGGTCGCCACACTCA<br>GATCAGTTAACCCCGTGATCTCCGAAGGGGGAGGTCGGATCTCTAGA |
| **P-6** | AGGCCTGGCGCCGACCGCACCACACTCACGAGGGCCCGCCCCACCAACAGGGGGCGGGCC<br>CTCTGTGCTGGCCTCAGGCGCCGACCGGGCTCGGTGCCCTCAAGCGCCGGCCGGGCTCCA<br>AGGGTGGCCTCAAGCGCCGGCCGGGCTGAGTTGGGCCGGTCTGGGCCCGCACGCGCGCCT<br>CACTGACGGCCTCAAGCGCCGGCCGGGCTATCTATAGCCCGGCCGGCGCTTGAGGCCGTC<br>TTTGGCGCGCGCCTGTGAGCGGACGGCCCGTCAAAGATCAGCCCGGCCGGCGCTTGAGGC<br>CATCTTTCGAGCCCGGCCGGCGTTTGAGGCCACCCCACCCCCGCCCCGGCAGGGGCGGCC<br>TGACCTCCGCATCCGCCGGCGCGGACAGGGCACCCCCAGTAGACGGGCGCGGGGCCGGAG<br>GCCCCTAGCGCCTTGCACTCTCCTACCCCGAGTGCTAATTATTGGCGTTAGCACTCTCCG<br>AGTGAGAGTGACAGAAGGACCGGGTCGGTGAGGCCCGCTGGCCACGCGGGGCAAGGAACC<br>GCGAGGCAGGCAGGCCGTCCGTCGCGGGCGCCAGCACGGTCCGGAGTATCCACCCTCCCC<br>CAGACAGAGTCCGGGGGGACCCCCAGTCCTGGGAGGACCACTTCACTCTAGA |
| **P-15** | AGGCCTTCCGCGCCGCCGGCCCGACGGTGCCCGGCCCCGTACCCCCCCCGGGTGGTGCGG<br>GGCCGGGCACCGGCCTTTTGGCGCTGCGGAGTTGACGGAAGTTGGCCGAACCGGATGCGC<br>TCGGCGCCCGGGGGCTGAAAGATGCTCACAGCCCCTTTCCACGGCGGTCCGGGAGGGGAG<br>GCCGGGCAACCGGTTTTCGGGGGCGGAGTGTCCGGTATGCGGACGGCCGCGCCCGATAGA<br>TGTGTAACGAGTCCGTTTCGCAACCATCTATCTCGGATCGGTTTGTCCGGATTTTGGAAG<br>ATGTGAGTGTCAGGTGTGATCGAACCGAGACCAAAAGGGTGTGGTCGGGCCGAACACCAT<br>GGCTAATAGTTGAGCGCGTAGAGCTCGGGTCAATGGGTCACGCGCTGTGGGGAGCGCCGA |

CTCACGAGCACACTGGGGCACTCGATCTTCGCCGTCAGGGGTGTCGGCGGATCGTCCTGT
GCCCTCTCTTGCAGTGAACAAGTGGACTCATGAGGAGGAACCCTCTAGA

**P-31**

AGGCCTCCGGACCTCTCCTCACGCTCACCCTGCGCGCTTCCGCGCGACAGGCACAATTAC
CCGTATATGTCCCGACTCGCCCACAGTCTCCGCCTTCGGCCGGGTCATTCCCCCGACCGA
CCCGGCCCGGCCCACCCATTTCCGGCCCGGCCGGCGTTTGAGGCCGACCGGTGACGGACA
CCCGAAGCCCTCGGAGCGCGCTCGGCATCAGCCCGGACGACGCTTGAGGCCACCTCGACC
GCCGCCGGACGGCTTCATCCGAAGTGCCTCTGAACTGGTAAAACGAGCCGTGCTGGCAGC
TCTCTGCACAACCAGGCAGAACAAAACTTGAGCCCGTCCGACTCAACCGCATTGACGCGC
CGCGTCCCCTCGTGCATCCTTGAGTGAGTTCCACTCAAGTAGTCAGCTGGAGGAATTGAC
TCTAGA

### 8. Cloning

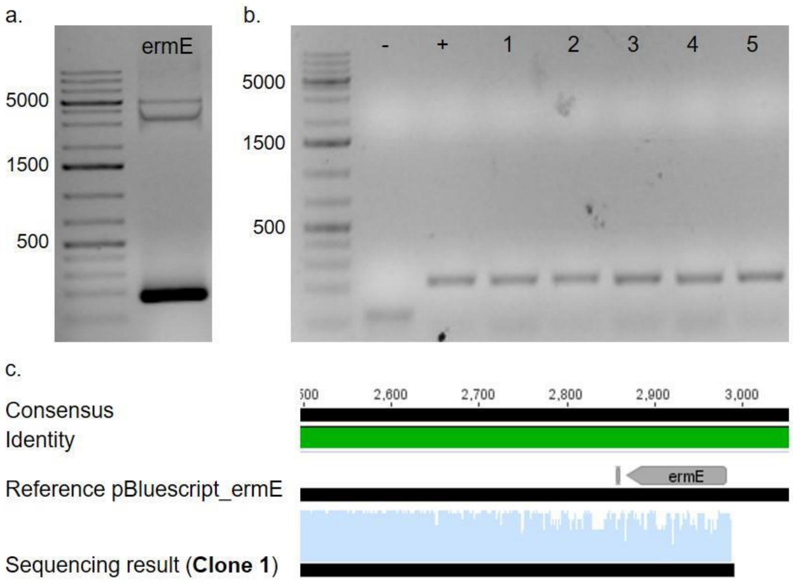


**Figure S35.** Assembly of pSET152-ermE expression vector. **a.** Amplification of the ermE promoter (183 bp). **b.** Colony PCR of clones obtained after transformation (positive: 266 bp; negative: 151 bp). **c.** Sanger sequencing revealed 100% identity with the reference sequence (green).

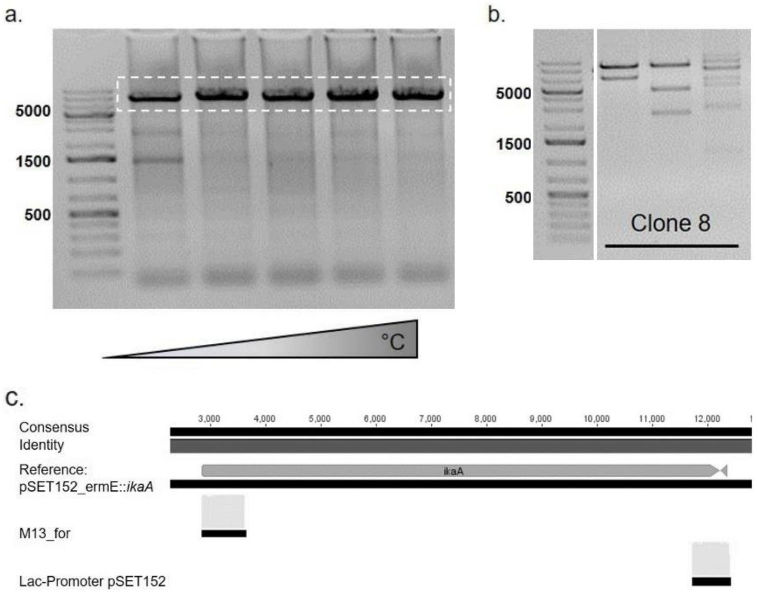


**Figure S36.** Assembly of pSET152-ermE-*ikaA* expression vector. **a.** Gradient PCR (50-70 °C) for the amplification of *ikaA* (9375 bp), suitable for Gibson assembly into pSET152-ermE. **b.** Analytical restriction digest (NsiI/StuI, NcoI/PciI, KpnI/BciVI) with clone 8 showing the predicted restriction pattern of pSET152-ermE-*ikaA*. **c.** Sanger sequencing revealed 100% identity with the reference sequence.

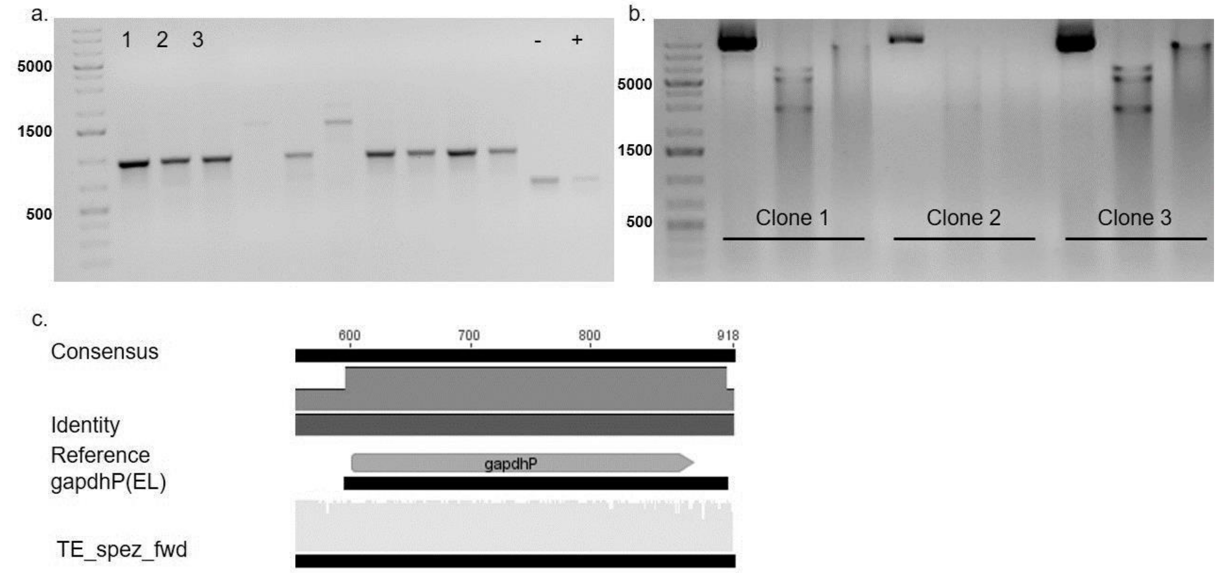


**Figure S37.** Verification of the cloning of pSET152-ermE-*ikaA*-gapdhP(EL). **a.** Colony screening PCR with expected bands for clones 1, 2, and 3 (919 bp). **b.** Analytical restriction digest (uncut, StuI/XbaI, PvuI/EcoRI) with clone 1 and 3 showing the predicted restriction pattern. **c.** Sanger sequencing of clone 1 revealed 100% identity with the reference sequence.

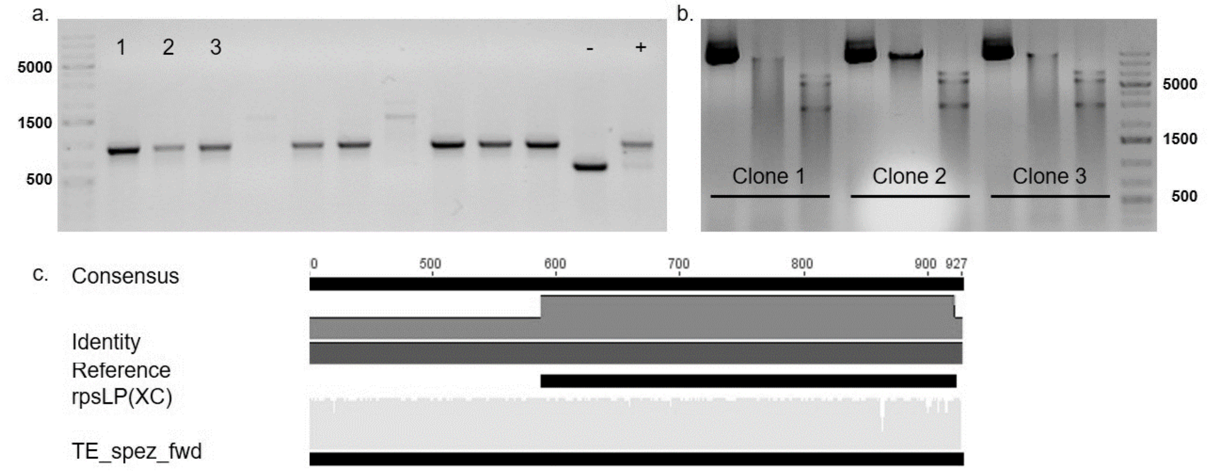


**Figure S38.** Verification of the cloning of pSET152-ermE-*ikaA*-rpsLP(XC). **a.** Colony screening PCR with expected bands for clones 1, 2, and 3 (936 bp). **b.** Analytical restriction digest (uncut, StuI/XbaI, PvuI/EcoRI) with all clones showing the predicted restriction pattern. **c.** Sanger sequencing of clone 1 revealed 100% identity with the reference sequence.

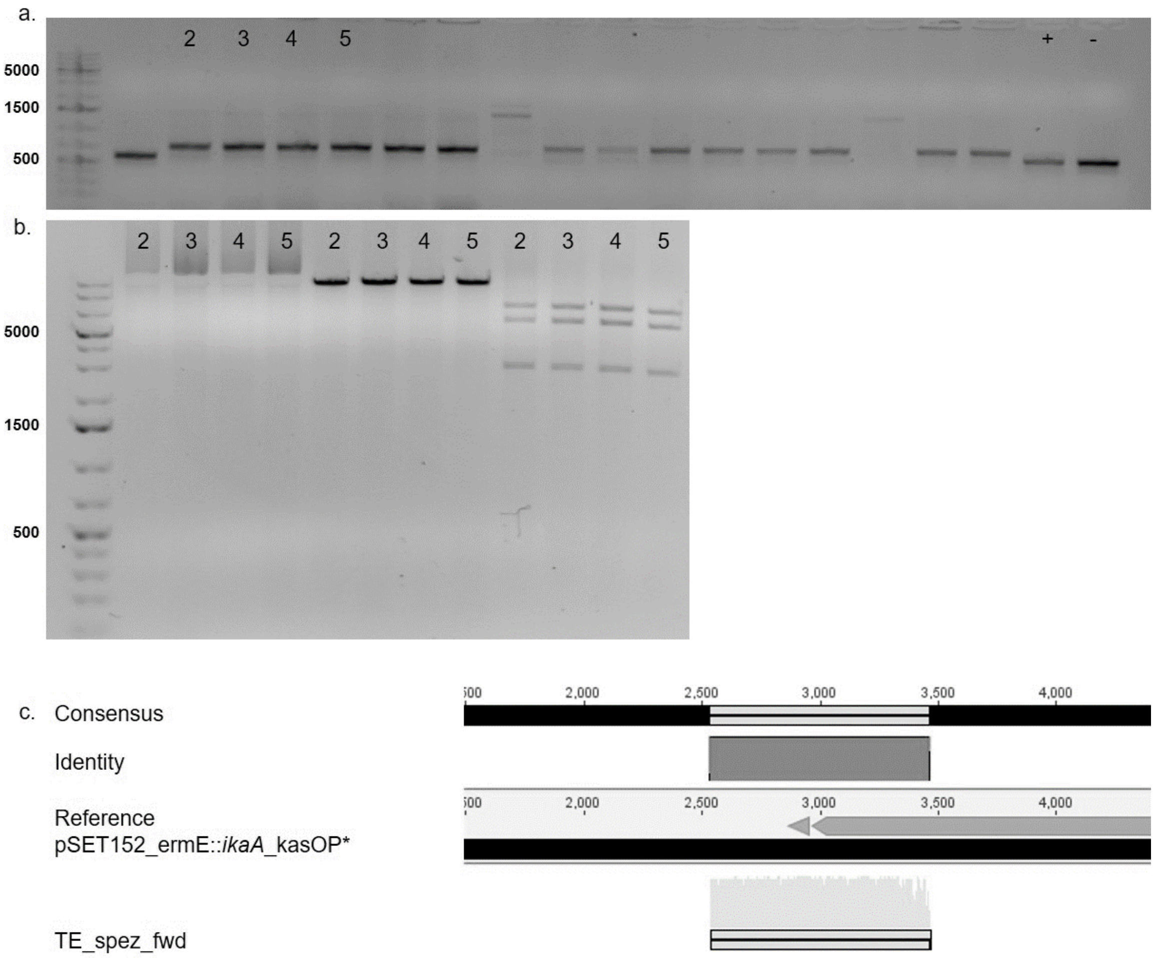


**Figure S39.** Verification of the cloning of pSET152-ermE-*ikaA*-kasOP*. **a.** Colony screening PCR with expected bands for clones 2, 3, 4, and 5 (731 bp). **b.** Analytical restriction digest (uncut, StuI/XbaI, PvuI/EcoRI) with all clones showing the predicted restriction pattern. **c.** Sanger sequencing of clone 2 revealed 100% identity with the reference sequence.

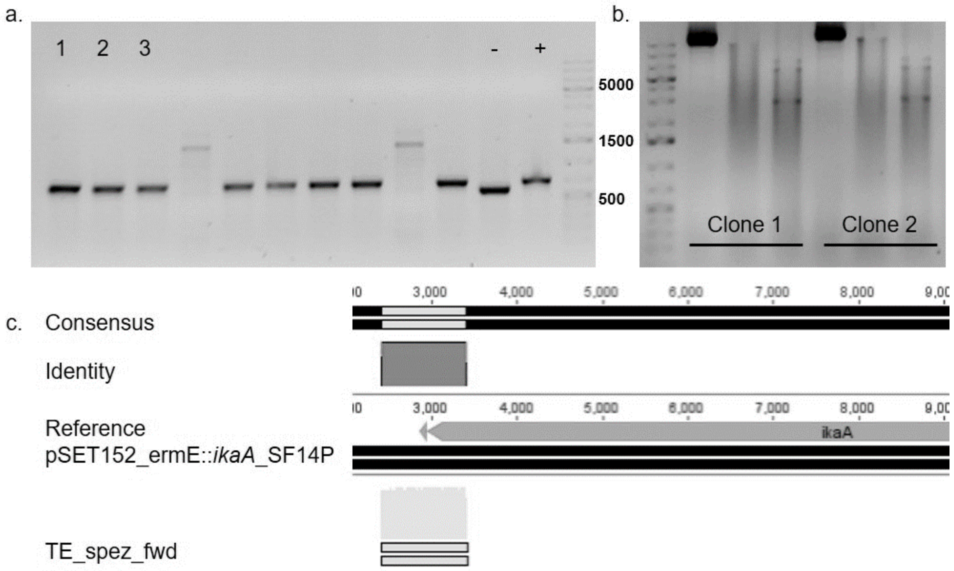


**Figure S40.** Verification of the cloning of pSET152-ermE-*ikaA*-SF14P. **a.** Colony screening PCR with expected bands for clones 1, 2, and 3 (717 bp). **b.** Analytical restriction digest (uncut, StuI/XbaI, PvuI/EcoRI) with all clones showing the predicted restriction pattern. **c.** Sanger sequencing of clone 1 revealed 100% identity with the reference sequence.

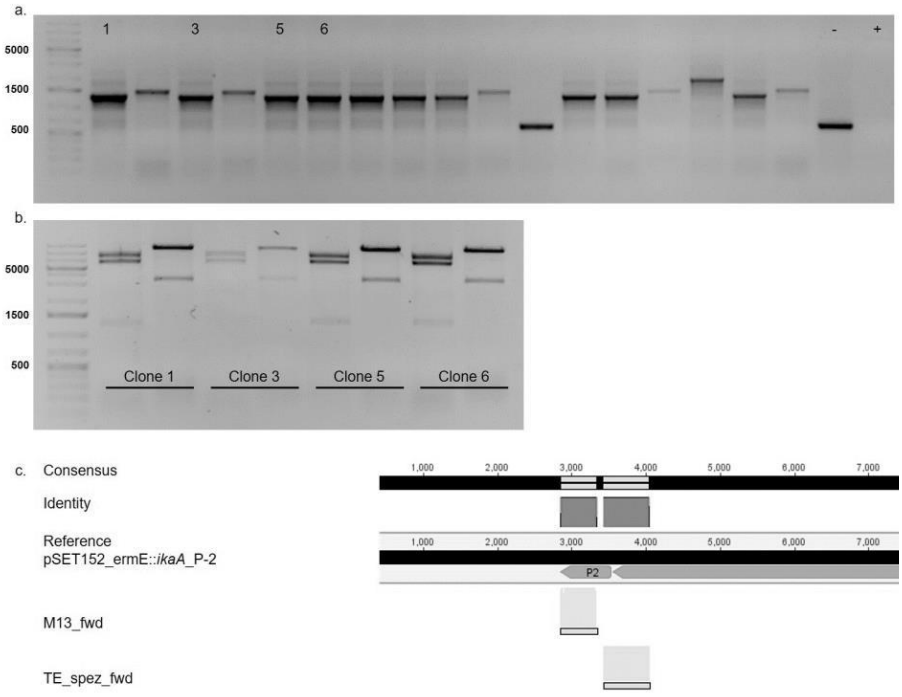


**Figure S41.** Verification of the cloning of pSET152-ermE-*ikaA*-P-2. **a.** Colony screening PCR with expected bands for clones 1, 3, 5, and 6 (1309 bp). **b.** Analytical restriction digest (NcoI/XbaI, PvuI) with all clones showing the predicted restriction pattern. **c.** Sanger sequencing of clone 1 revealed 100% identity with the reference sequence.

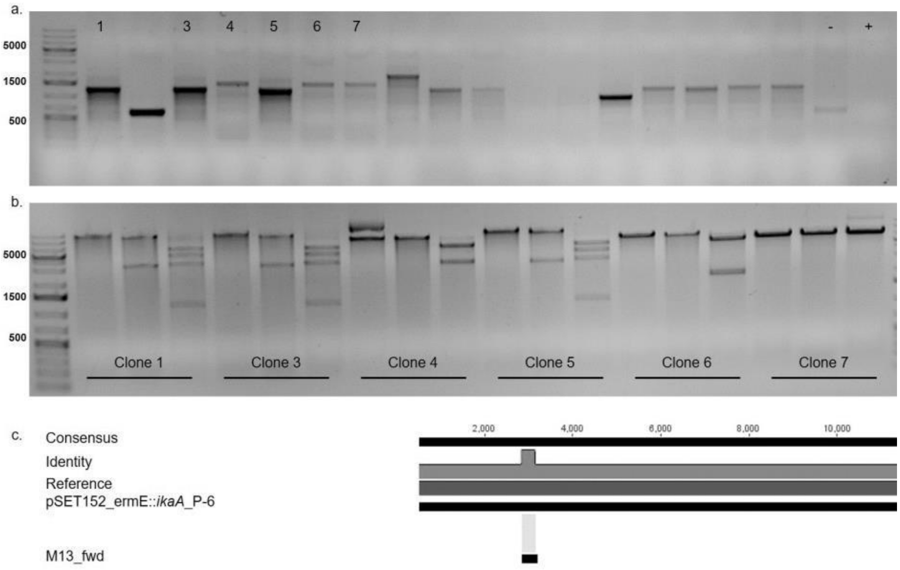


**Figure S42.** Verification of the cloning of pSET152-ermE-*ikaA*-P-6. **a.** Colony screening PCR with expected bands for clones 1, 3, 4, 5, 6 and 7 (1254 bp). **b.** Analytical restriction digest (XbaI/StuI, PvuI, XmnI) with all clones showing the predicted restriction pattern. **c.** Sanger sequencing of clone 1 revealed 100% identity with the reference sequence.

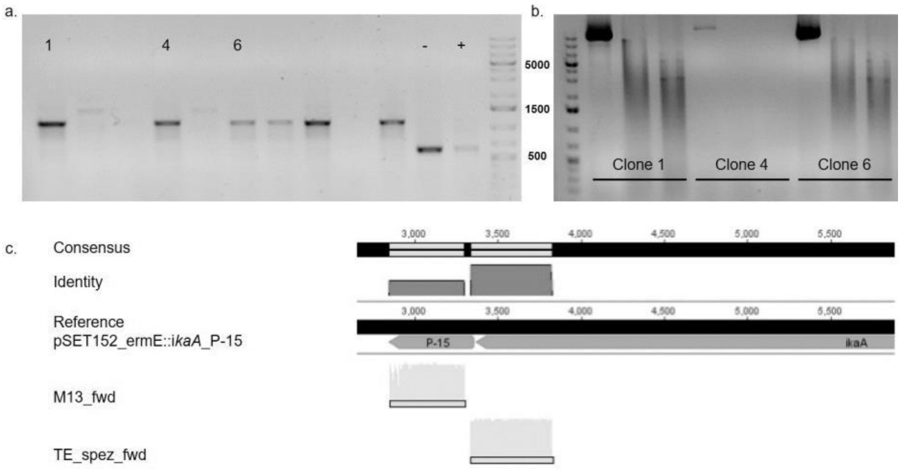


**Figure S43.** Verification of the cloning of pSET152-ermE-*ikaA*-P-15. **a.** Colony screening PCR with expected bands for clones 1, 4, and 6 (1131 bp). **b.** Analytical restriction digest (uncut, StuI/XbaI, PvuI/EcoRI) with all clones showing no conclusive bands, but **c.** Sanger sequencing of clone 1 revealed 100% identity with the reference sequence.

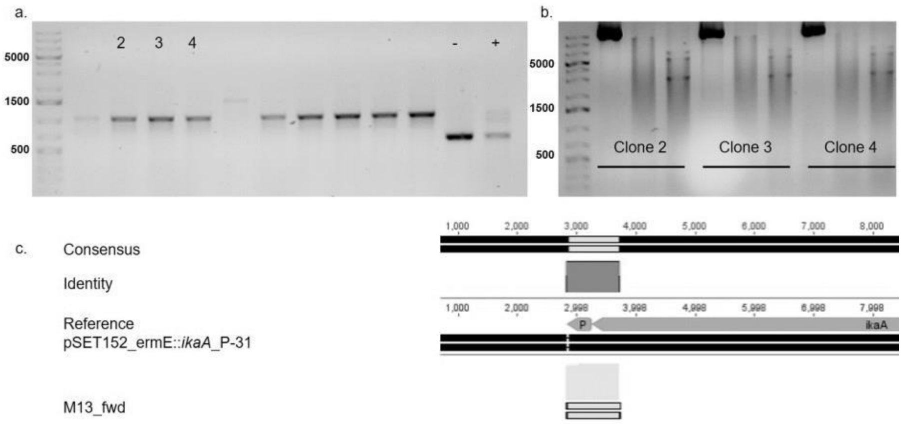


**Figure S44.** Verification of the cloning of pSET152-ermE-*ikaA*-P-31. **a.** Colony screening PCR with expected bands for clones 2, 3, and 4 (1028 bp). **b.** Analytical restriction digest (uncut, StuI/XbaI, PvuI/EcoRI) with all clones showing the predicted restriction pattern. **c.** Sanger sequencing of clone 2 revealed 100% identity with the reference sequence.

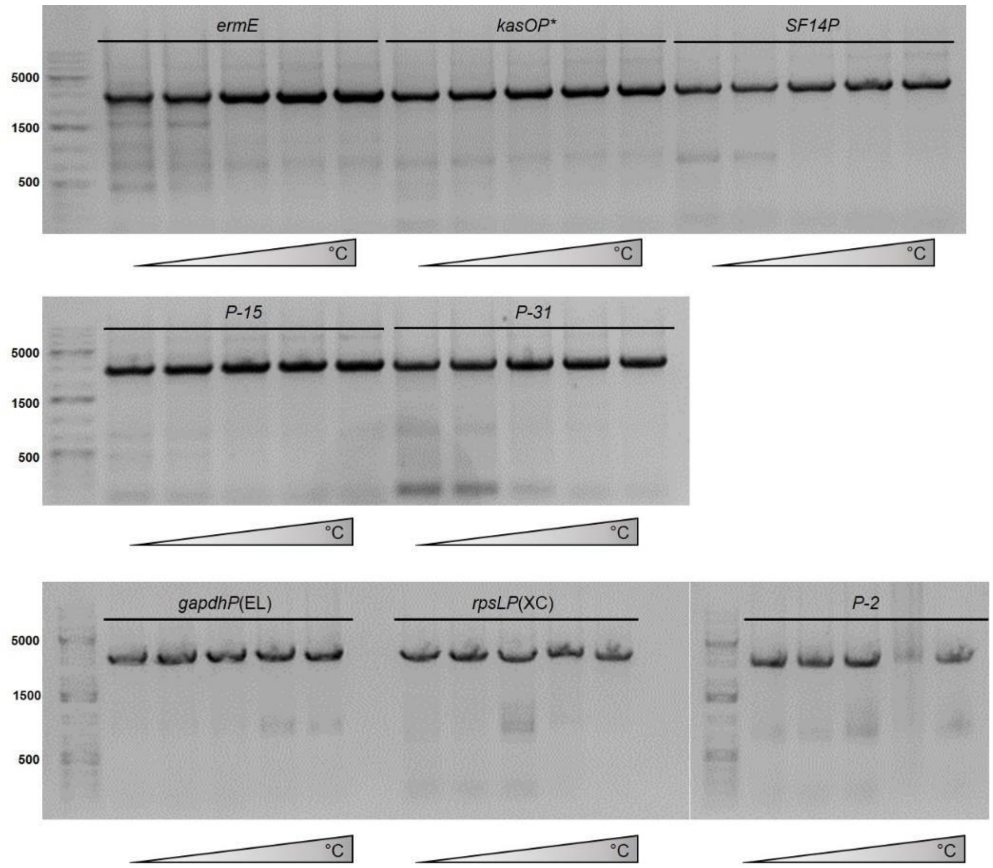


**Figure S45.** PCR amplification of *ikaBC*. The two genes (2937 bp) were amplified together for the eight different expression constructs and simultaneously homologous arms were added for SLIC cloning.

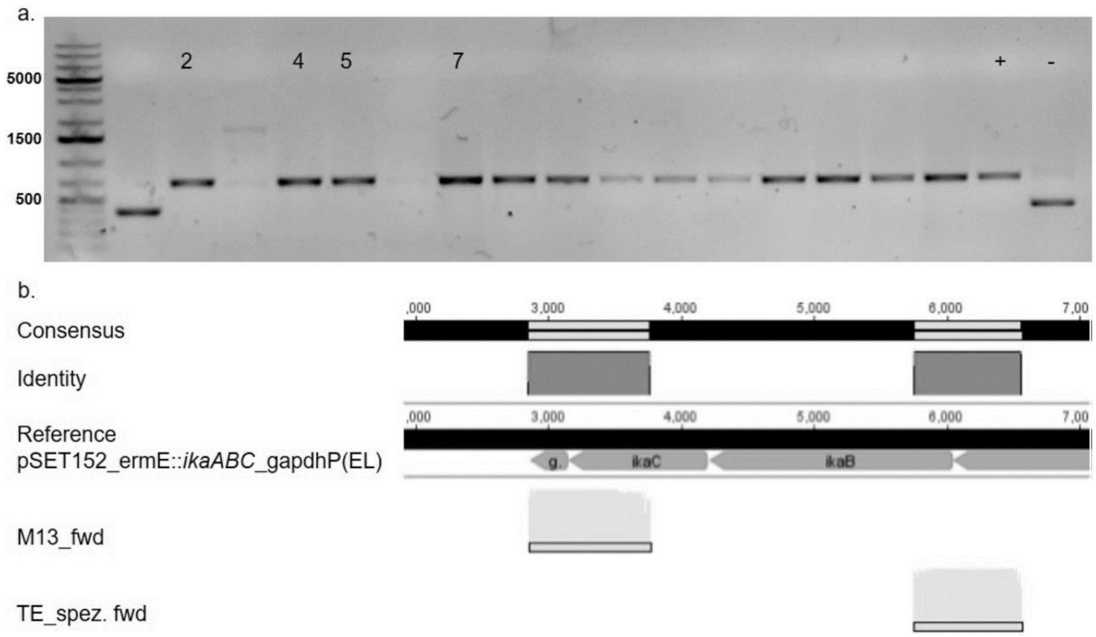


**Figure S46.** Verification of the cloning of pSET152-ermE-*ikaABC*-gapdhP(EL). **a.** Colony screening PCR with expected bands for clones 2, 4, 5, and 7 (677 bp). **b.** Sanger sequencing of clone 2 revealed 100% identity with the reference sequence.

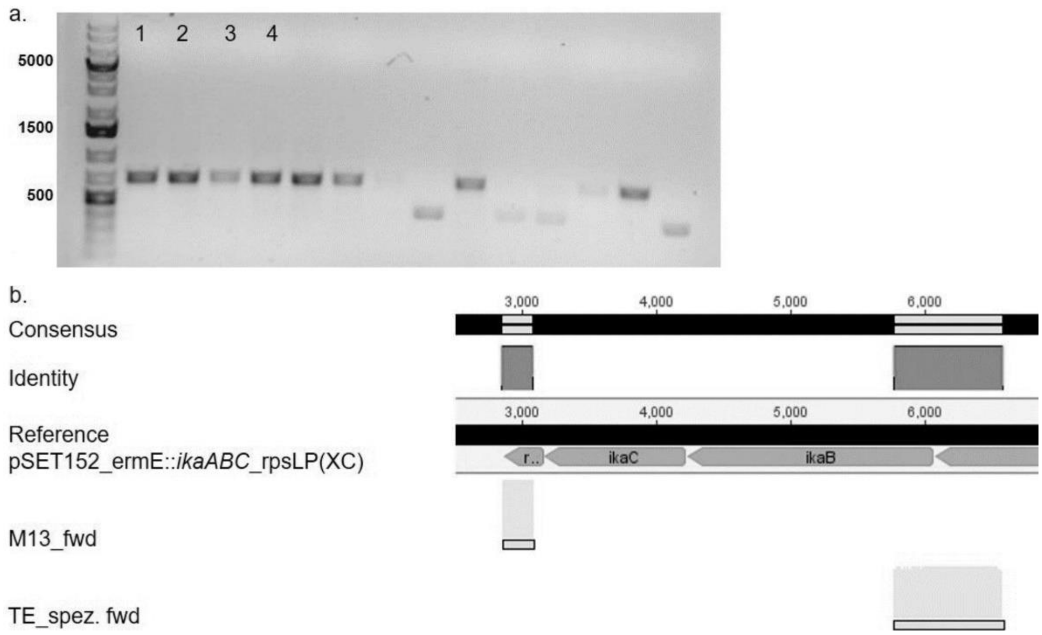


**Figure S47.** Verification of the cloning of pSET152-ermE-*ikaABC*-rpsLP(XC). **a.** Colony screening PCR with expected bands for clones 1, 2, 3, and 4 (694 bp). **b.** Sanger sequencing of clone 1 revealed 100% identity with the reference sequence.

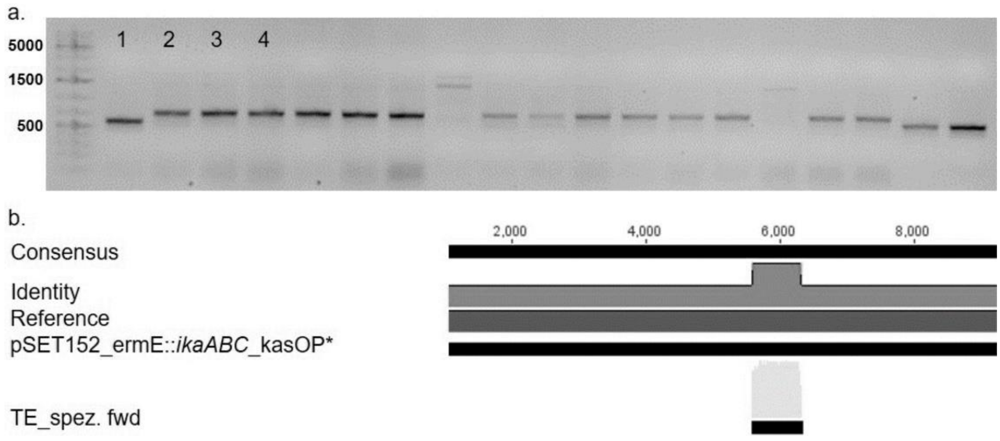


**Figure S48.** Verification of the cloning of pSET152-ermE-*ikaABC*-kasOP*. **a.** Colony screening PCR with expected bands for clones 1, 2, 3, and 4 (731 bp). **b.** Sanger sequencing of clone 3 revealed 100% identity with the reference sequence.

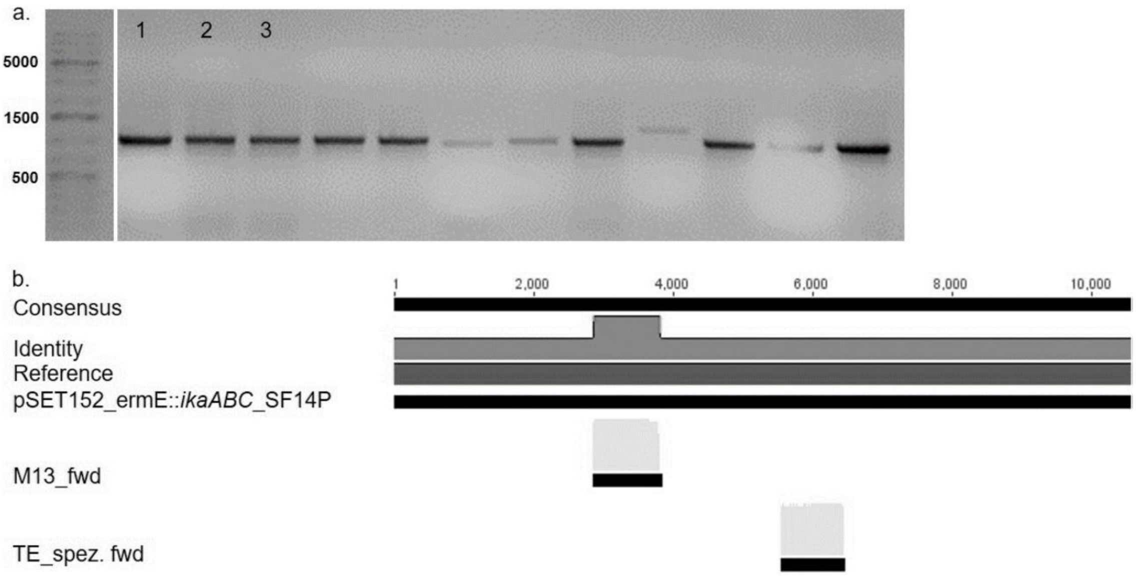


**Figure S49.** Verification of the cloning of pSET152-ermE-*ikaABC*-SF14P. **a.** Colony screening PCR with expected bands for clones 1, 2, and 3 (1082 bp). **b.** Sanger sequencing of clone 1 revealed 100% identity with the reference sequence.

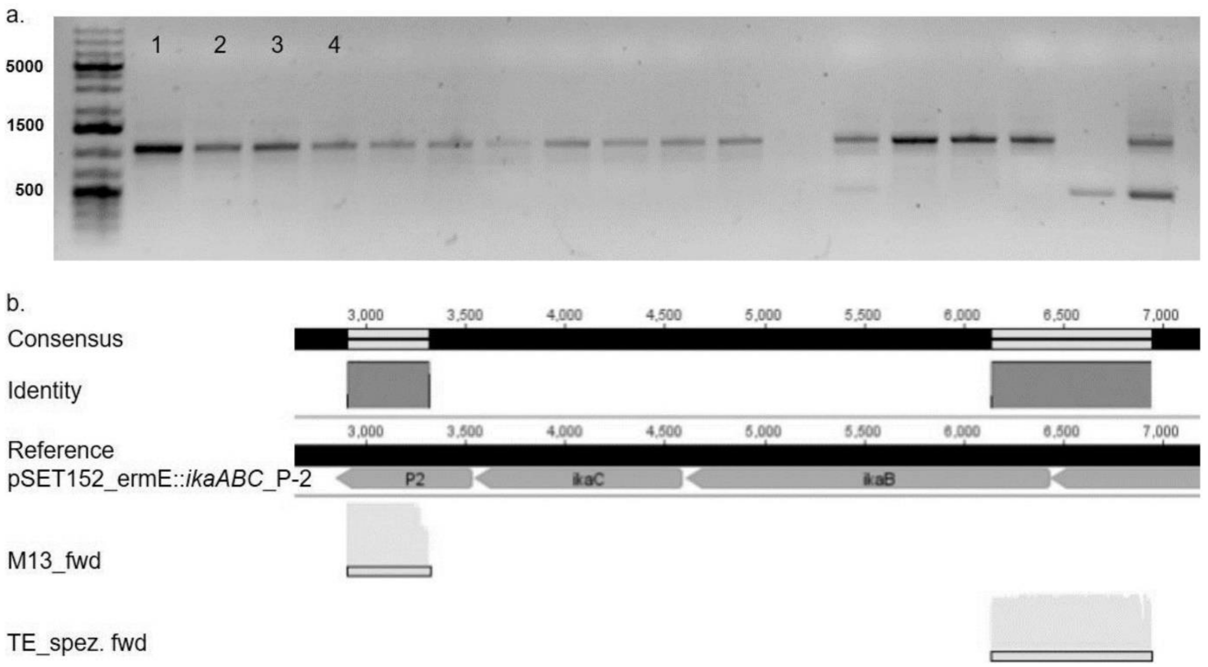


**Figure S50.** Verification of the cloning of pSET152-ermE-*ikaABC*-P-2. **a.** Colony screening PCR with expected bands for clones 1, 2, 3, and 4 (1067 bp). **b.** Sanger sequencing of clone 1 revealed 100% identity with the reference sequence.

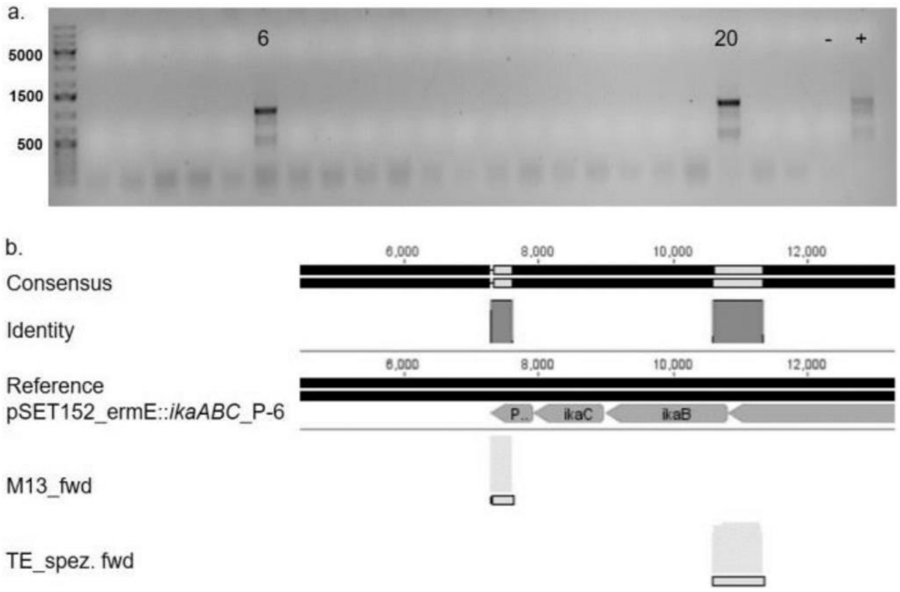


**Figure S51.** Verification of the cloning of pSET152-ermE-*ikaABC*-P-6. **a.** Colony screening PCR with expected bands for clones 6 and 20 (1012 bp). **b.** Sanger sequencing of clone 20 revealed 100% identity with the reference sequence.

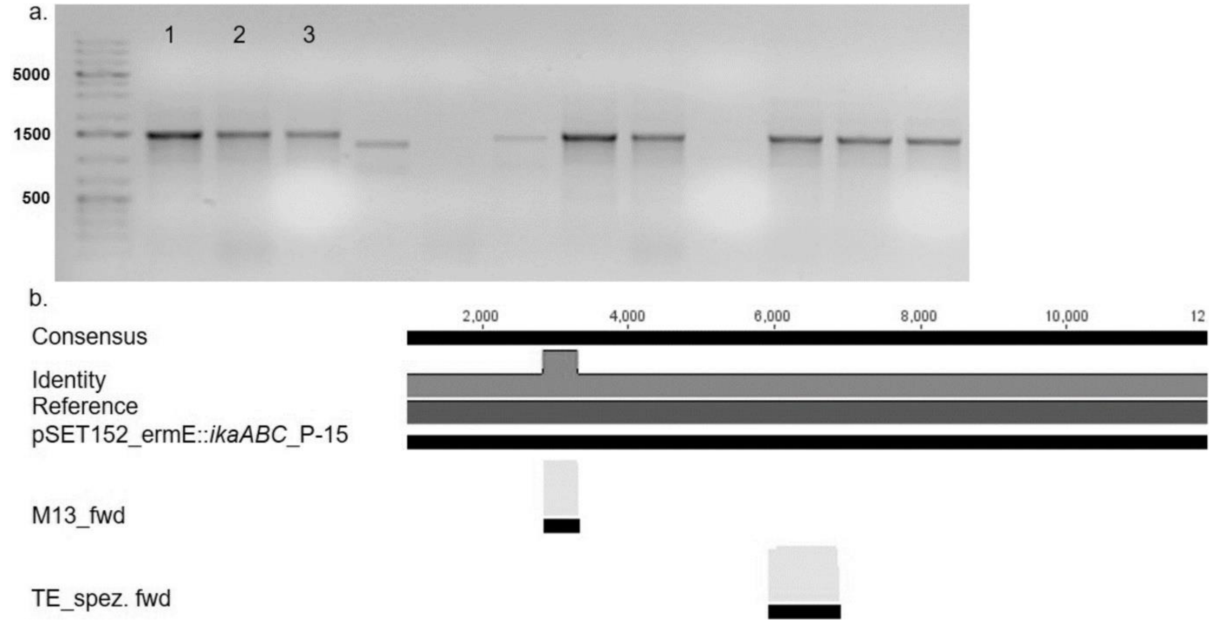


**Figure S52.** Verification of the cloning of pSET152-ermE-*ikaABC*-P-15. **a.** Colony screening PCR with expected bands for clones 1, 2, and 3 (1496 bp). **b.** Sanger sequencing of clone 1 revealed 100% identity with the reference sequence.

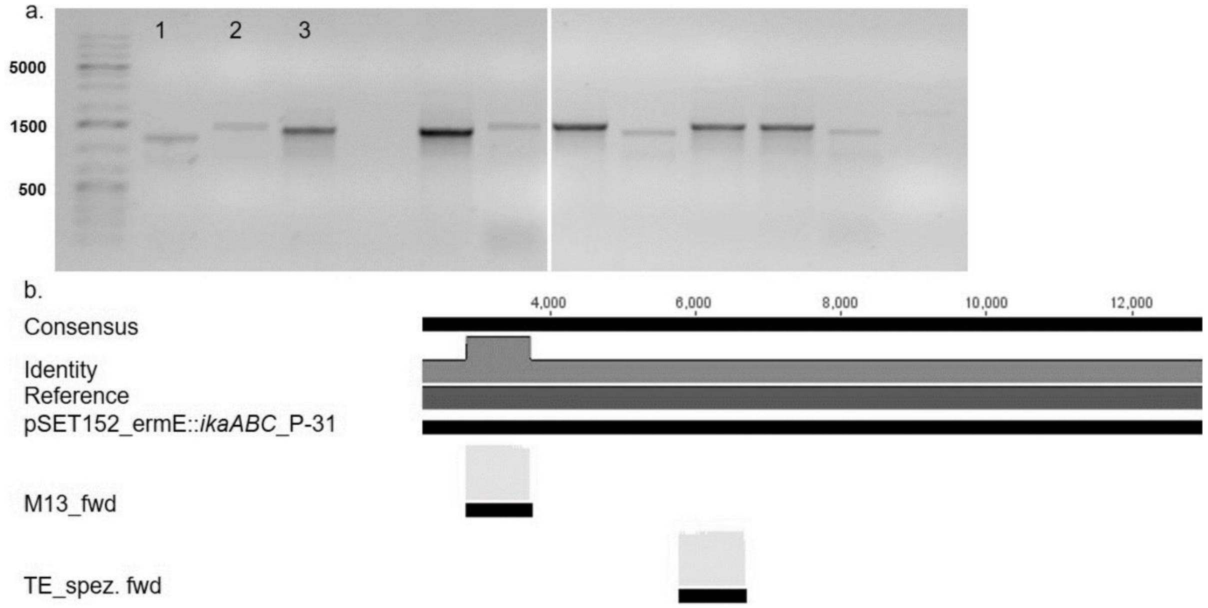


**Figure S53.** Verification of the cloning of pSET152-ermE-*ikaABC*-P-31. **a.** Colony screening PCR with expected bands for clones 1, 2, and 3 (1393 bp). **b.** Sanger sequencing of clone 1 revealed 100% identity with the reference sequence.

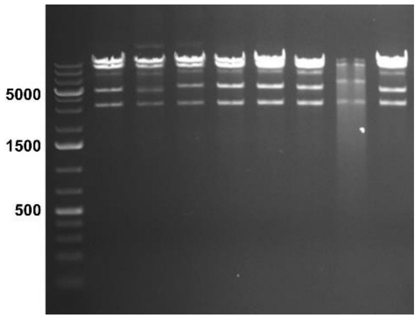


**Figure S54.** Analytical restriction digest of the plasmids pSET152-ermE-*ikaABC*-P2. The second promoters (P2) were gapdhP(EL), kasOP*, P-2, P-6, P-15, P-31, rpsLP(XC), and SF14P (from left to right). All plasmids were digested with SphI/EcoRI and showed the predicted restriction pattern.

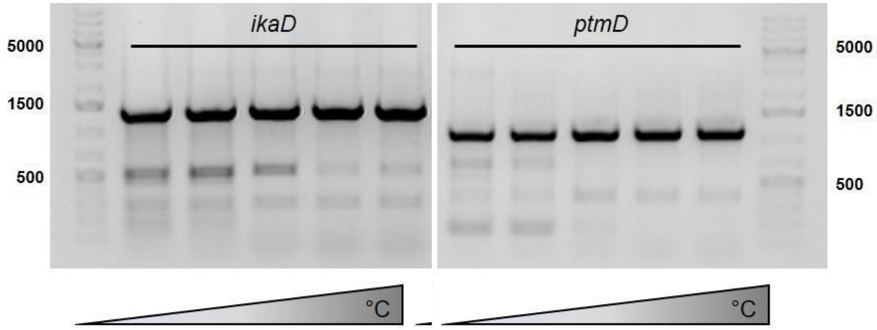


**Figure S55.** Amplification of *ikaD* (1388 bp) and *ptmD* (1085 bp).

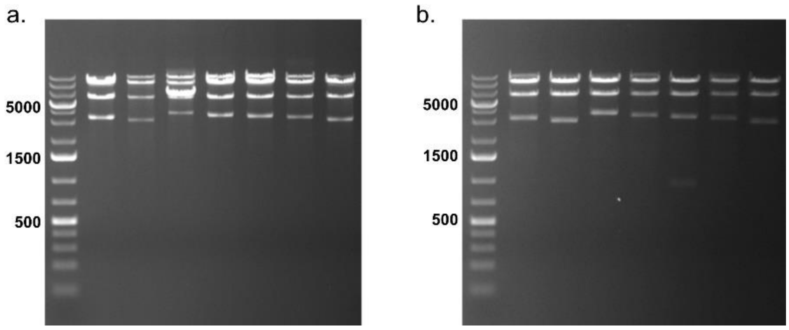


**Figure S56.** Analytical restriction digest of the final expression plasmids containing the modifying genes. **a.** Digest of pSET152-ermE-*ikaABC*-P2-ikaD with PciI/StuI. The second promoters (P2) were gapdhP(EL), kasOP*, P-2, P-15, P-31, rpsLP(XC), and SF-14 (from left to right). All plasmids showed the predicted restriction pattern. **b.** Digest of pSET152-ermE-*ikaABC*-P2-*ikaD* with EcoRI/StuI. The second promoters (P2) were gapdhP(EL), kasOP*, P-2, P-15, P-31, rpsLP(XC), and SF-14 (from left to right). All plasmids showed the predicted restriction pattern.

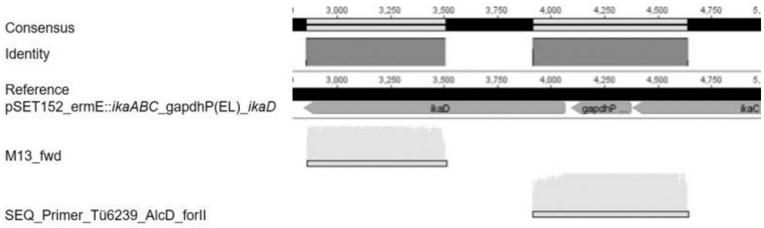


**Figure S57.** Sequencing result pSET152-ermE-*ikaABC*-gapdhP(EL)-*ikaD*.

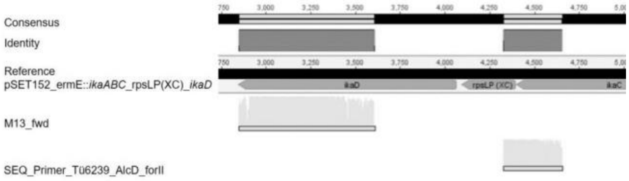


**Figure S58.** Sequencing result pSET152-ermE-*ikaABC*-rpsLP(XC)-*ikaD*.

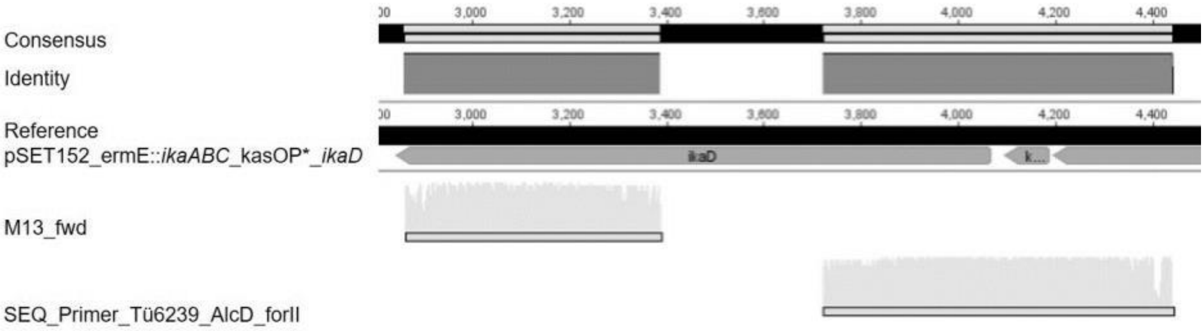


**Figure S59.** Sequencing result pSET152-ermE-*ikaABC*-kasOP*-*ikaD*.

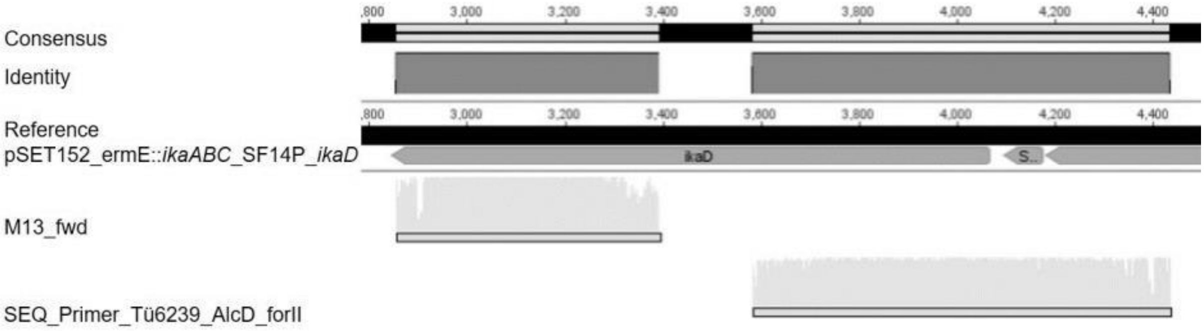


**Figure S60.** Sequencing result pSET152-ermE-*ikaABC*-SF14P-*ikaD*.

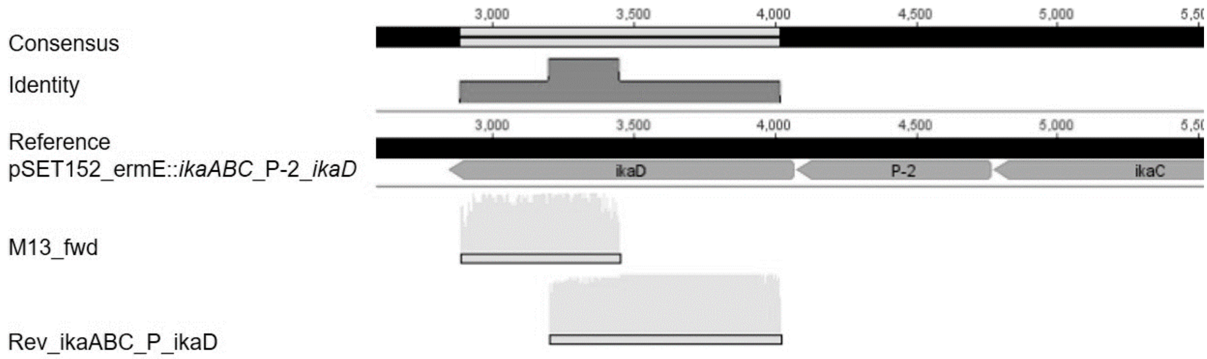


**Figure S61.** Sequencing result pSET152-ermE-*ikaABC*-P-2-*ikaD*.

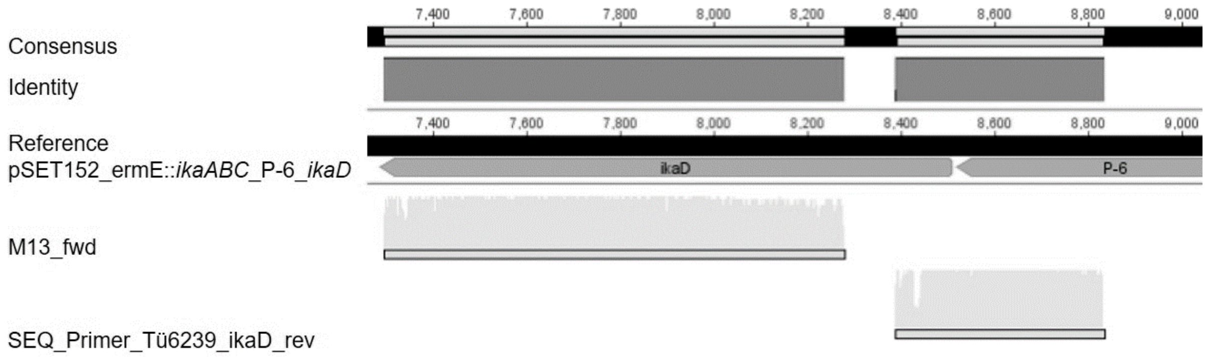


**Figure S62.** Sequencing result pSET152-ermE-*ikaABC*-P-6-*ikaD*.

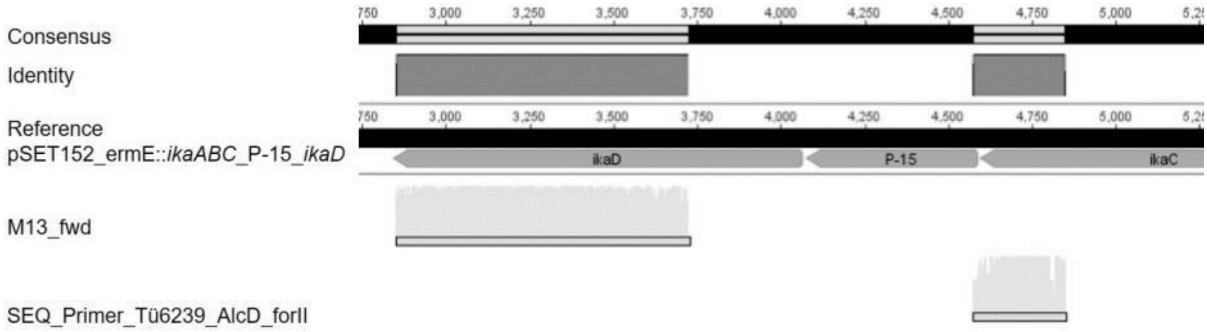


**Figure S63.** Sequencing result pSET152-ermE-*ikaABC*-P-15-*ikaD*.

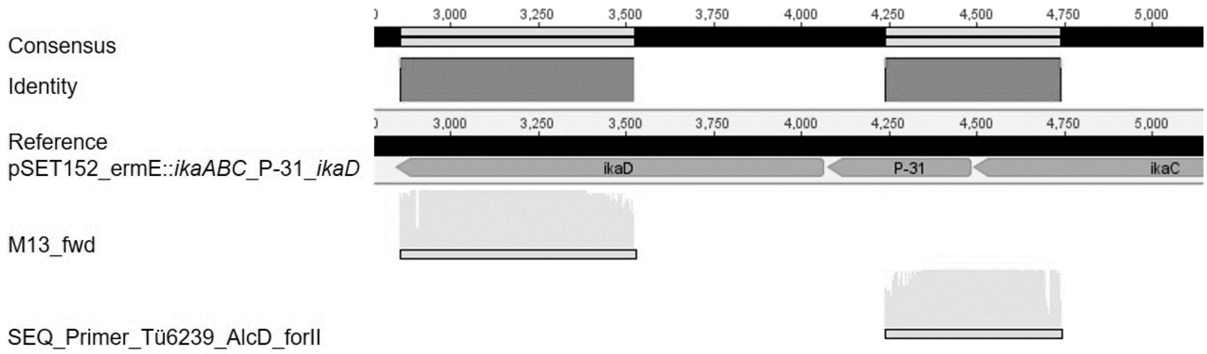


**Figure S64.** Sequencing result pSET152-ermE-*ikaABC*-P-31-*ikaD*.

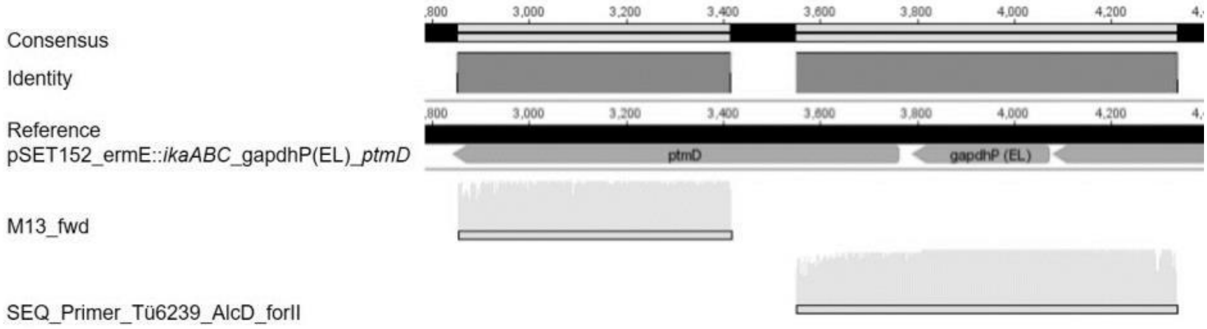


**Figure S65.** Sequencing result pSET152-ermE-*ikaABC*-gapdhP(EL)-*ptmD*.

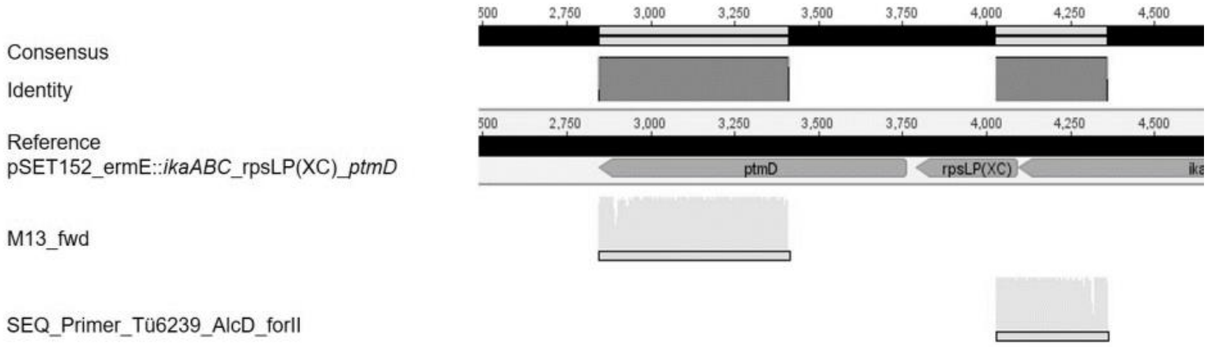


**Figure S66.** Sequencing result pSET152-ermE-*ikaABC*-rpsLP(XC)-*ptmD*.

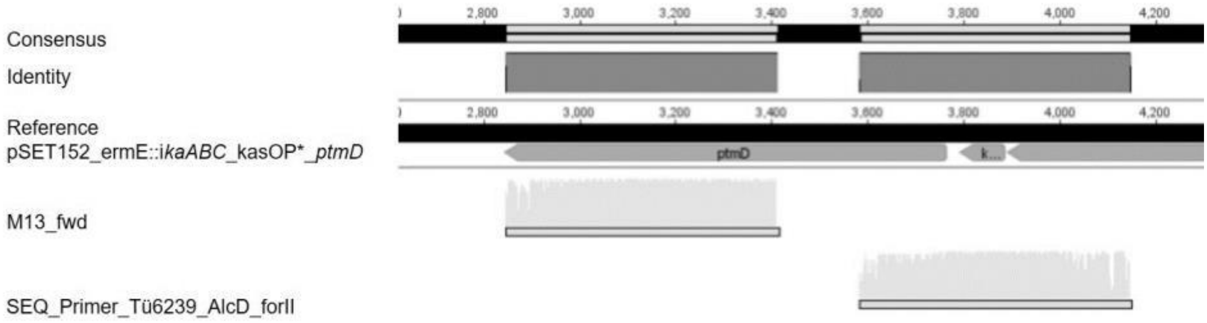


**Figure S67.** Sequencing result pSET152-ermE-*ikaABC*-kasOP*-*ptmD*.

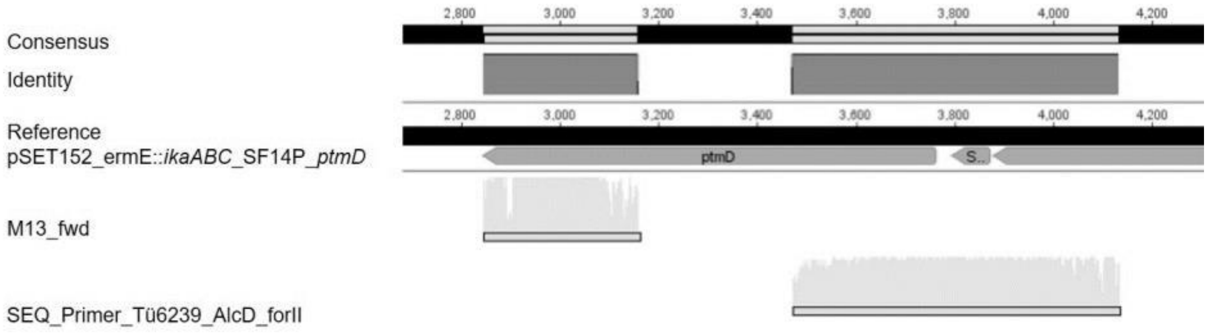


**Figure S68.** Sequencing result pSET152-ermE-*ikaABC*-SF14P-*ptmD*

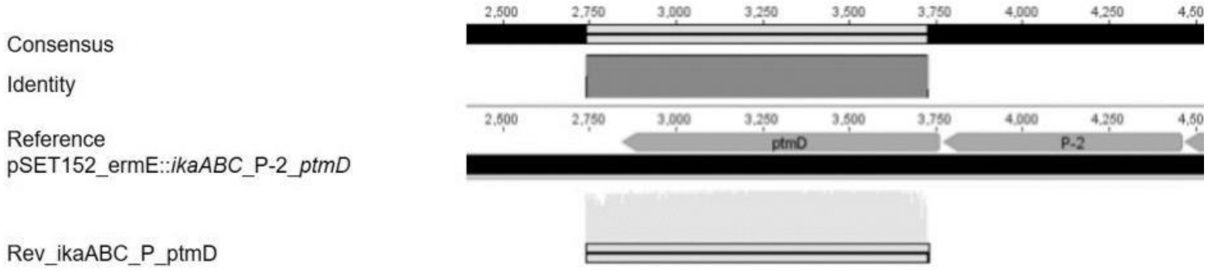


**Figure S69.** Sequencing result pSET152-ermE-*ikaABC*-P-2-*ptmD*.

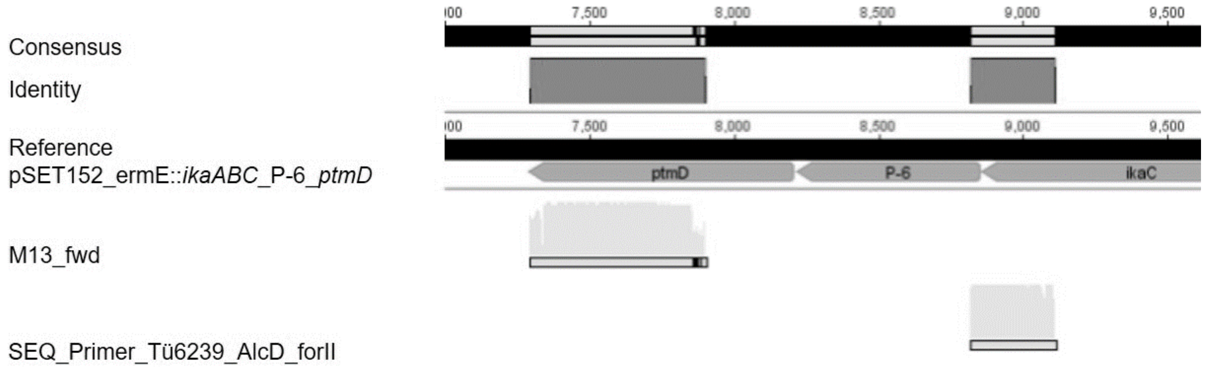


**Figure S70.** Sequencing result pSET152-ermE-*ikaABC*-P-6-*ptmD*.

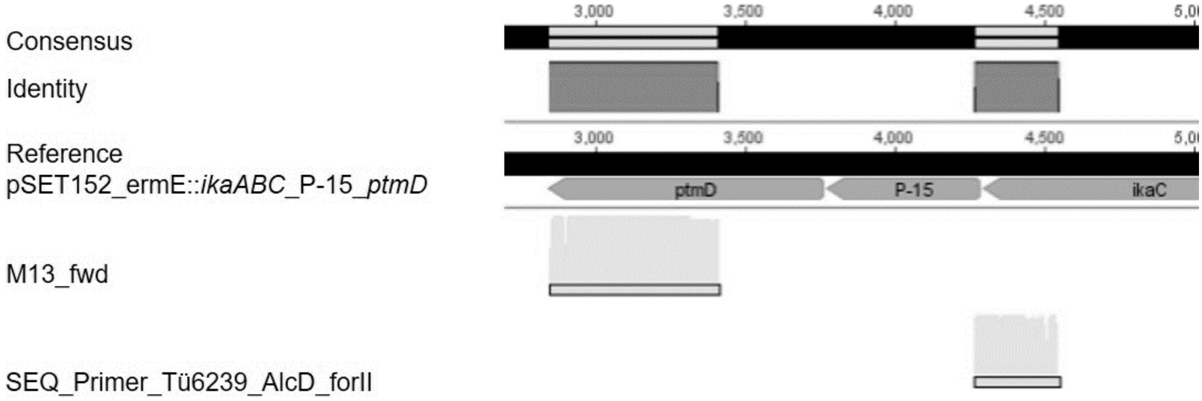


**Figure S71.** Sequencing result pSET152-ermE-*ikaABC*-P-15-*ptmD*.

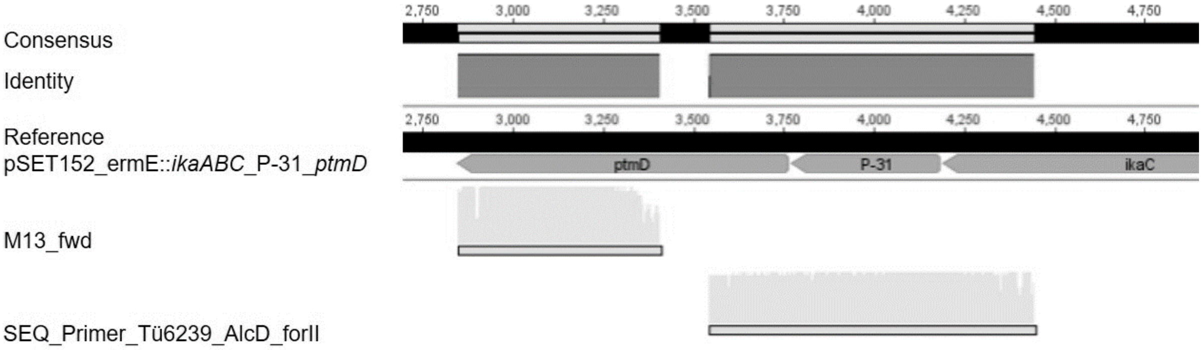


**Figure S72.** Sequencing result pSET152-ermE-*ikaABC*-P-31-*ptmD*.

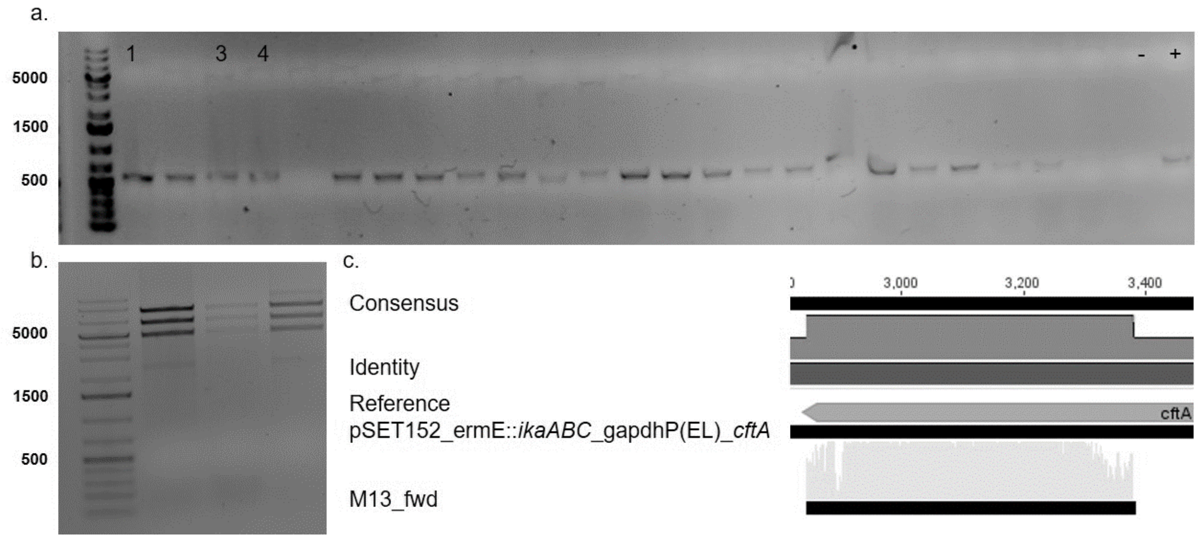


**Figure S73.** Verification of the cloning of pSET152-ermE-*ikaABC*-gapdhP(EL)-*cftA*. **a** Colony screening PCR with expected bands for clones 1, 3, and 4 (480 bp). **b.** Analytical restriction digest (NcoI) with all clones showing the predicted restriction pattern. **c.** Sanger sequencing of clone 1 revealed 100% identity with the reference sequence.

#### 9. Literature


1. MacNeil, D. J.; Gewain, K. M.; Ruby, C. L.; Dezeny, G.; Gibbons, P. H.; MacNeil, T., Analysis of *Streptomyces avermitilis* genes required for avermectin biosynthesis utilizing a novel integration vector. *Gene* **1992,** *111*, 61–68.
2. Labeda, D. P.; Doroghazi, J. R.; Ju, K.-S.; Metcalf, W. W., Taxonomic evaluation of *Streptomyces albus* and related species using multilocus sequence analysis and proposals to emend the description of *Streptomyces albus* and describe *Streptomyces pathocidini* sp. nov. *Int. J. Syst. Evol. Microbiol.* **2014,** *64*, 894–900.
3. Hopwood, D. A.; Kieser, T.; Wright, H. M.; Bibb, M. J., Plasmids, Recombination and Chromosome Mapping in *Streptomyces lividans* 66. *J. Gen. Microbiol.* **1983,** *129*, 2257–2269.
4. Kieser, T.; Hopwood, D. A.; Wright, H. M.; Thompson, C. J., pIJ101, a multi-copy broad host-range *Streptomyces* plasmid: Functional analysis and development of DNA cloning vectors. *Molec. Gen. Genet.* **1982,** *185*, 223–238.
5. Wang, G.; Hosaka, T.; Ochi, K., Dramatic Activation of Antibiotic Production in *Streptomyces coelicolor* by Cumulative Drug Resistance Mutations. *Appl. Environ. Microbiol.* **2008,** *74*, 2834–2840.
